# A maize near-isogenic line population designed for gene discovery and characterization of allelic effects

**DOI:** 10.1101/2025.01.29.635337

**Authors:** Tao Zhong, Alex Mullens, Laura Morales, Kelly L Swarts, William C Stafstrom, Yijian He, Shannon M Sermons, Qin Yang, Luis O Lopez-Zuniga, Elizabeth Rucker, Wade E Thomason, Rebecca J Nelson, Tiffany Jamann, Peter J Balint-Kurti, James B Holland

## Abstract

In this study we characterized a panel of 1,264 maize near-isogenic lines (NILs), developed from crosses between 18 diverse inbred lines and the recurrent parent B73, referred to as nested NILs (nNILs). 884 of the nNILs were genotyped using genotyping-by-sequencing (GBS). Subsequently, 24 of these nNILs, and all the parental lines, were re-genotyped using a high-density SNP chip. A novel pipeline for calling introgressions, which does not rely on knowing the donor parent of each nNIL, was developed based on a hidden Markov model (HMM) algorithm. By comparing the introgressions detected using GBS data with those identified using chip data, we optimized the HMM parameters for analyzing the entire nNIL population. A total of 2,972 introgressions were identified across the 884 nNILs. Individual introgression blocks ranged from 21 bp to 204 Mbp, with an average size of 17 Mbp. By comparing SNP genotypes within introgressed segments to the known genotypes of the donor lines we determined that in about one third of the lines, the identity of the donors did not match expectation based on their pedigrees.

We characterized the entire nNIL population for three foliar diseases. Using these data, we mapped a number of quantitative trait loci (QTL) for disease resistance in the nNIL population and observed extensive variation in effects among the alleles from different donor parents at most QTL identified. This population will be of significant utility for dissecting complex agronomic traits and allelic series in maize.

**Significance Statement:** The study reports the characterization of a publicly available population of 1,264 maize near-isogenic lines largely derived from a single recurrent parent and 18 donor lines. This population is likely to be of significant utility for the characterization of allelic series at loci of interest.

## Introduction

Near-isogenic line (NIL) populations typically consist of a set of lines in which each line carries a small part of the genome of a “donor parent” in the genetic background of a “recurrent parent” (Szalma *et al*., 2007). Typically, each line within the population carries a different part, or “introgression”, from the donor genome, such that, over the population, the entire genome of the donor parent (or close to it) is represented. While NIL populations can be used for the identification of loci with effects on traits of interest (e.g. Lennon *et al*., 2016), they have lower statistical power for this purpose than other populations, such as recombinant inbred line (RIL) populations, in which allele frequencies are balanced (Kaeppler, 1997). NIL populations, however, provide the opportunity to identify pairs of lines that differ for specific alleles but that are otherwise almost genetically identical. As such they are very useful for the characterization of the effects imparted by specific alleles, including their mechanisms and dominance effects.

Here we characterize a population of 1,264 NILs, termed the nested NIL (nNIL) population, all of which use the B73 inbred as the recurrent parent. This population is divided into 18 subpopulations, based on the identity of the reported donor parent (CML103, CML228, CML247, CML277, CML322, CML333, CML52, CML69, Ki11, Ki3, M162W, Mo17, Mo18W, NC350, NC358, Oh43, Tx303, and Tzi8). All 19 of these lines are among founders of the widely-used and well characterized maize nested association mapping (NAM) population (Gage *et al*., 2020, Hufford *et al*., 2021).

We report the genotyping of 884 of the 1,264 nNILs, using a genotyping-by-sequencing (GBS) approach. Identifying introgressions from low-coverage GBS data can be difficult because of high rates of missing data and relatively low rates of informative markers. Most SNPs have low minor allele frequency, such that most pairs of individuals will have identical in state SNP genotypes at most SNPs (Hufford *et al*., 2021). Thus, without knowledge of parental genotypes, a set of informative SNPs cannot be determined *a priori*, such that even in regions of introgression, progeny line genotypes will match a recurrent parent at most SNPs. To verify accurate introgression calling from GBS data, we developed and optimized a novel pipeline for calling introgressions, which did not rely on knowledge of the donor parent of each nNIL. We also independently genotyped 24 of these lines and all parental lines using a high-density genotyping chip which yields datasets characterized by low missing data and low error rates to use as a validation data set to evaluate the accuracy of introgression-calling from GBS data. We phenotyped the entire nNIL population for resistance to the fungal foliar diseases gray leaf spot, northern leaf blight, and southern leaf blight, as well as for several other agronomic traits. We report the use of the nNIL population for identification of QTL for resistance to these diseases and for characterization of allelic series at these QTL with differing effects.

## RESULTS

### Optimization of introgression calling

The GBS and chip SNP genotype datasets are provided in Files S1 and S2. We tested our Hidden Markov Model (HMM) introgression calling pipeline across a range of parameters for non-informative rate (*nir*), genotyping error rate on homozygotes (*ger_m_*), genotyping error rate on heterozygotes (*ger_t_*), probability that genotyping error on a homozygote results in a heterozygous call (*p*), and probability of recombination between adjacent markers. The *nir* represents the overall probability that donor parent and B73 SNP alleles are identical-in-state, such that the SNP is not informative about introgression in lines derived from that donor parent. Ideally, the parameters set for the HMM should result in zero introgressions called on the B73 samples and 100% introgression calls in the donor parents included in the chip genotyping. The chip genotyping dataset included the recurrent parent, B73, as well as all the donor parents. Therefore, we searched for a combination of parameters that identified zero introgressions in B73 and as close to 100% homozygous introgressions as possible on the donor parents with the chip data. Then, we searched independently for the combination of parameters in the HMM applied to GBS data that resulted in the highest match rate to the optimized chip introgression calls for the subset of 24 nNILs genotyped with both methods. Of all the parameters tested, the *nir* had the largest impact on results (Figures S1 and S2). All parameter settings tested resulted in 0% introgression calls in B73 (Figure S1). We observed that increasing *nir* values increased the proportion of homozygous introgression calls in the donor parents (Figure S1). Non-informative rate values between 0.5 and 0.9 resulted in greater than 90% homozygous introgression calls in the donors. Changing the values for parameters reflecting genotyping error rates (*ger_m_* and *ger_t_*) and recombination frequency settings had only small impacts on the results. The highest rate of introgression calls for the donor founders was for *nir* = 0.9, *ger_m_* = 0.01, *ger_t_* = 0.001 or lower, and r = 0.0007, for which the HMM returned 98% homozygous and 0.3% heterozygous introgression calls (File S3). At this setting, the 24 nNILs included in the chip data set had 5% homozygous and 0.3% heterozygous introgression calls (Figure S1). This level of introgression is higher than the expected rate of 1.51% for this generation (BC_5_) if NILs were selected randomly, but some marker-assisted selection was applied during their development that likely increased the proportion of introgression. The level of introgression called in the NILs was somewhat robust to the parameter settings; the lowest level of introgression called in the NILs in this grid search was 2.1% (File S3).

Next, we used our HMM introgression calling pipeline with these same ranges of parameter settings to call introgressions in the nNILs using GBS data (but changing the mean adjacent marker recombination rate to reflect the larger number of GBS markers). For each combination of GBS parameter settings, we compared introgression calls between GBS data and the previously selected set of chip data introgression calls for the 24 nNILs genotyped with both methods (Figure S2). The markers in the chip data had little overlap with the GBS markers, so, to compare the introgressions calls of nNILs using chip markers to the GBS markers, we identified the nearest GBS marker to each chip marker and imputed the introgression calls on the GBS markers. The mean introgression call mismatch was lowest (0.078%) for GBS parameter settings *nir* = 0.9, *ger_m_* = 0.01, *ger_t_* = 0.0001 and r = 0.00047 (Figure S2; File S4). Across all 884 nNILs genotyped with GBS, we called a mean of 4.4% of markers as homozygous introgressions and 0.4% as heterozygous introgressions. These were similar proportions to those derived from the chip data.

We visualized the comparisons of haplotype calls from the two data sets for the common set of 24 nNILs (Figure 1). Large-scale differences in introgression calls were observed for three of the 24 nNILs. CMLL277/B73 NIL-1086 had a large introgression on chromosome 3 based on chip data that was not called from the GBS data. Tzi8/B73 NIL-1107 had introgression blocks called in different positions across the genome, with one small region of overlap on chromosome 8 and another block on chromosome 1 that was called as a single large introgression from GBS data but as several smaller introgressions disrupted by recombinations from chip data. Tzi8/B73 NIL-1220 had a similar discrepancy on chromosome 7 where chip data indicated a large homozygous introgression, but GBS data indicated a series of disrupted homozygous and heterozygous introgressions. This last discrepancy may represent some difficulty in distinguishing heterozygous from homozygous introgressions from low coverage sequence data. The other large differences in CMLL277/B73 NIL-1086 and Tzi8/B73 NIL-1107 are more likely due to heterozygous introgressions segregating out differently in distinct sub-lines, as the seed sources for the chip and GBS data sets had diverged for at least two generations of selfing.

**Figure 1.**
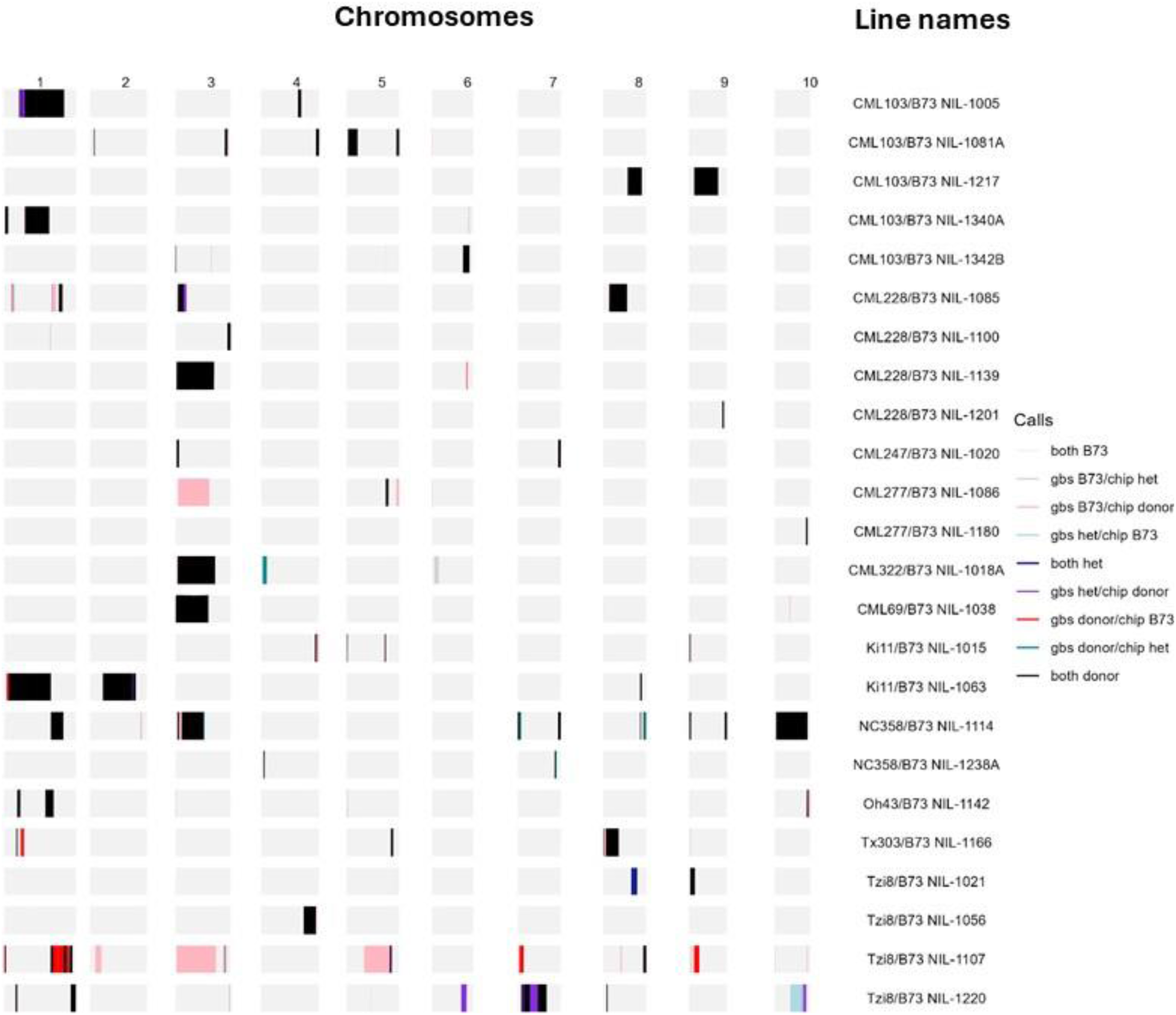
Comparison of 24 nNILs genotyped with both chip and GBS methods. The black and the light grey bars identify regions at which both the GBS and the chip data suggest that the DNA derives from donor or B73 parents respectively. Other colors, as described in the key, denote regions where the two genotypic datasets suggest diverging origins for the DNA at those loci.

We compared SNP calls within introgression regions to NAM founder HapMap 3.2.1 SNP calls independently for the introgression calls and SNPs based on the chip data vs the introgression calls and SNPs based on the GBS data to identify which founder sequence best matched the introgression sequences of each nNIL. The NAM founder with highest proportion of matching SNP calls agreed between chip and GBS data for 22 of 24 nNILs for which we had chip data. The two lines where GBS and chip data disagreed on the closest matching founder, Tzi8/B73 NIL-1107 and Tzi8/B73 NIL-1220, were also identified previously as lines where the introgression calls differed substantially between GBS and chip data sets (Figure 1; Table 1). For Tzi8/B73 NIL-1107, GBS data matched the founder expected by pedigree (Tzi8) best, whereas the chip data identified Mo17 as the closest founder match. For Tzi8/B73 NIL-1220, the best founder match from both genotyping methods disagreed with the expected founder based on pedigree (Table 1). In addition, both genotyping methods identified a common highest matching founder for introgression regions that was discordant with the expected donor founder based on pedigree in three of other 22 nNILs (Table 1).

**Table 1.**
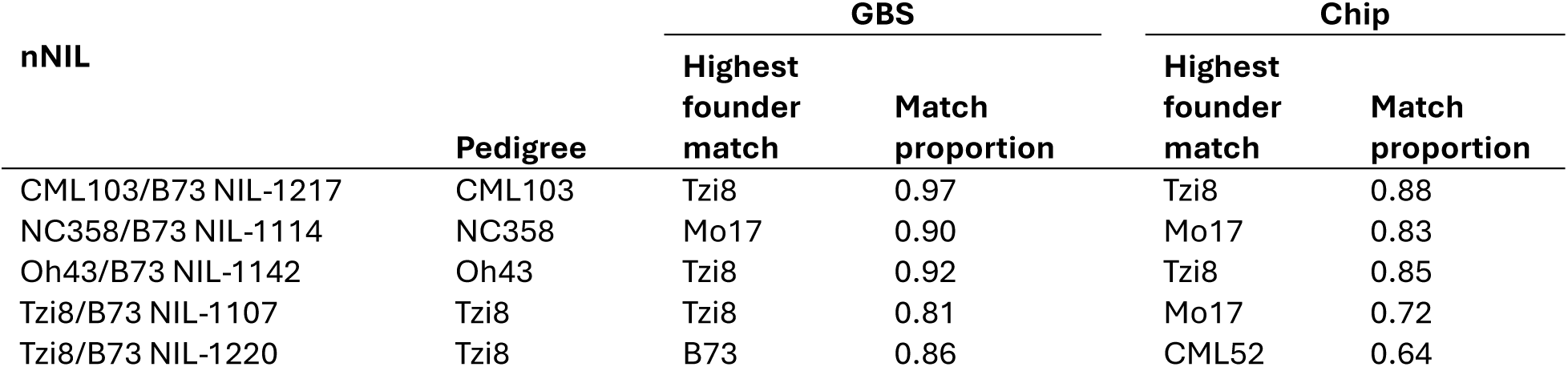
nNILs genotyped by both GBS and chip for which the highest NAM founder match is discordant with the reported pedigree.

We also identified closest founder matches within introgressions for the full set of nNILs genotyped by GBS. Twenty (2%) of lines had fewer than 20 SNPs called as donor introgressions; we did not attempt to match donor founders to these lines. Another 11% of lines had enough introgressed SNPs to test sequence matches with founders but did not match a donor parent with certainty. Finally, 20% of lines were matched with high probability to a founder that disagreed with their pedigree. In total therefore, 33% of lines had uncertain or discrepant matches to NAM founders within their introgressions. Introgression calling and imputation methods that rely on correct pedigrees would be expected to have difficulty with this data set. More than 30% of nNILs putatively derived from CML52, Ki3, NC350, NC358, and Oh43 did not match their pedigree parent (Table 2, File S5).

**Table 2.** Proportion of nNILs with discrepancies between the reported donor parents by pedigree and the best NAM founder SNP call matches within introgressed regions. Number of nNILs phenotyped includes all lines in the original population with the reported pedigree.

| Reported donor by pedigree | Number of nNILs<br>genotyped | Proportion of nNILs<br>matching different<br>founder | Number of nNILs<br>phenotyped |
| --- | --- | --- | --- |
| CML103 | 94 | 0.24 | 108 |
| CML228 | 42 | 0.17 | 50 |
| CML247 | 43 | 0.09 | 64 |
| CML277 | 45 | 0.20 | 56 |
| CML322 | 45 | 0.11 | 63 |
| CML333 | 74 | 0.09 | 93 |
| CML52 | 56 | 0.50 | 77 |
| CML69 | 47 | 0.17 | 57 |
| Ki11 | 44 | 0.16 | 61 |
| Ki3 | 29 | 0.31 | 64 |
| M162W | 18 | 0.00 | 73 |
| Mo17 | 27 | 0.11 | 65 |
| Mo18W | 75 | 0.04 | 92 |
| NC350 | 44 | 0.34 | 65 |
| NC358 | 38 | 0.34 | 69 |
| Oh43 | 59 | 0.34 | 72 |
| Tx303 | 47 | 0.30 | 62 |
| Tzi8 | 57 | 0.05 | 73 |

### Summary of Introgressions

File S6 details the introgressions detected in each of the genotyped nNILs. A total of 2,972 contiguous introgressions were identified across the 884 nNILs meaning that nNILs carried on average 3.36 introgression blocks per line. The total proportion of genome introgressed was computed for each line. Two NILs had very high rates of introgression: CML247/B73 NIL-1015 (63%) and CML247/B73 NIL-1005A (56%). Both of these lines were previously identified as having uncertain donor matches and, furthermore, it was noted during field evaluation (see below) that they did not closely resemble B73. It is almost certain therefore that these lines were contaminated by other pollen sources in later generations of the breeding process. NILs previously identified as having uncertain or incorrect donor founder matches had higher mean introgression rates (more than 3.5%) than the NILs that matched their pedigree donor parent inside introgressions (2.4%). These higher introgression rates and unexpected donor matches suggest that pollen contamination may also influence these two groups at a significantly higher rate than the NILs whose introgression blocks matched their pedigree donors. Therefore, the introgression block sizes and genome proportions summarized below use only the 593 lines with high probability matches to their pedigree donor, which have a mean of 3.02 introgression blocks per line.

The distribution of introgressions was uneven across the genome (Figure 2). The number of nNILs carrying an introgression at a given marker ranged from 6 to 72, with a mean of 40. Individual introgression blocks ranged in estimated size from 21 bp to 204 Mbp with a mean of 17 Mbp (Figure S3). As a proportion of the total length of the chromosome in which an introgression occurred, introgressions represented a range from < 1% to 90% with a mean of 8% of the whole chromosome (Figure S4). Summing introgressions within lines and computing the proportion of the whole genome represented by introgressions revealed that lines varied from almost 0% to 17% of their genome introgressed from a NAM donor, with a mean of 2.4% (Figure S5). This value is closer to the expected value of 1.5% for BC5-derived lines than the 4.6% proportion of introgressed markers estimated from the full nNIL set. The proportion of introgressed markers does not account for the uneven distribution of markers throughout the genome and, more importantly, the full set of nNILs includes a significant number of lines that may have contaminations. Therefore, we believe that this estimated proportion of the genome introgressed is a more reliable estimate of the extent of introgressions among those lines.

**Figure 2.**
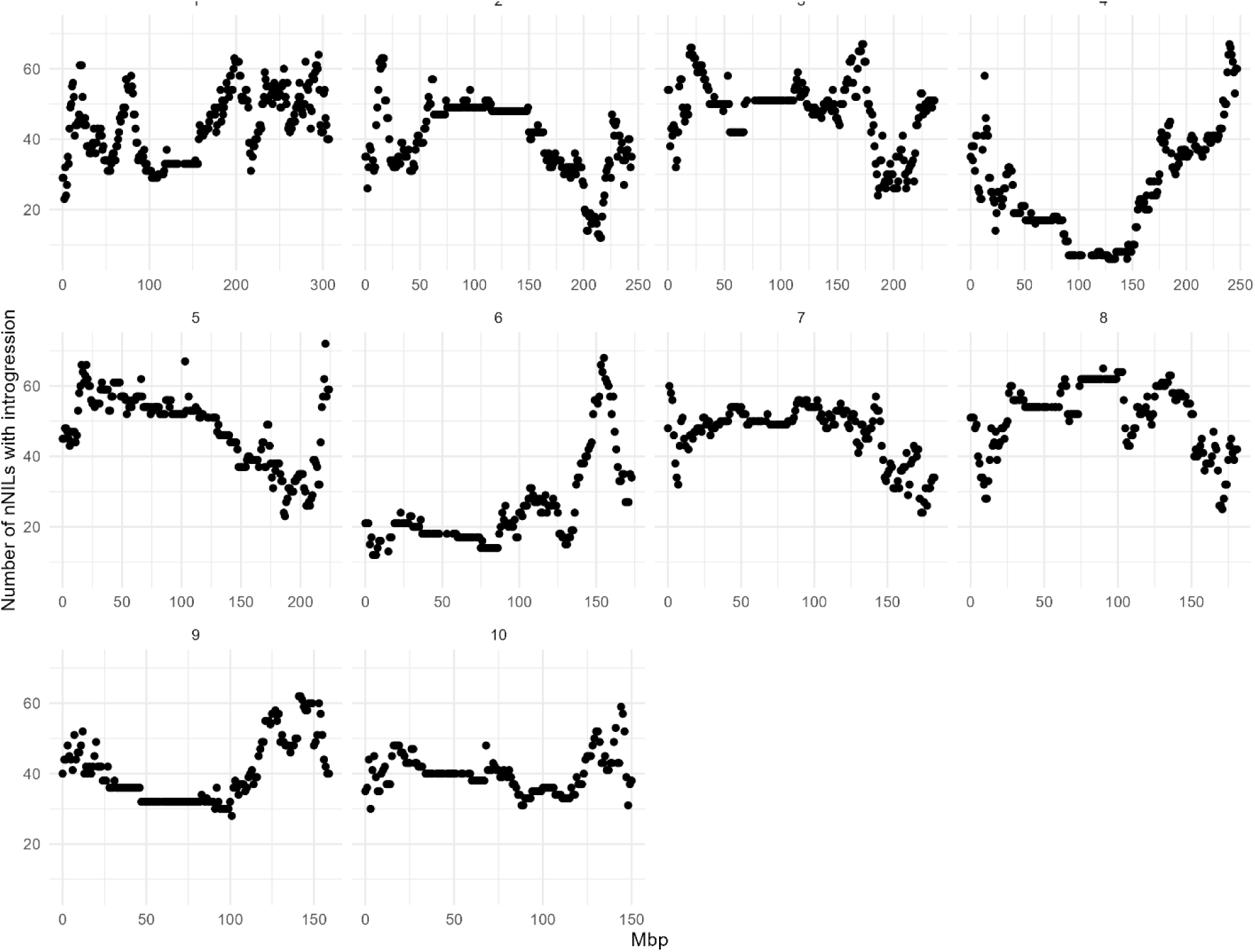
Introgression coverage by genome position among the 593 lines for which the pedigree was confirmed to be accurate across the 10 chromosomes of maize. Chromosome number indicated at the top of each of the ten graphs.

### Trait evaluations

File S7 includes all the phenotypic data recorded for the population. We evaluated the 1,264 nNILs for flowering time (DTA), height (EHT, PHT), and resistance to three foliar diseases (GLS, NLB, SLB) across four to six environments each. For each of the three diseases assessed, we separately estimated heritability of line means for each individual rating (first and second rating) across environments as well as for the mean rating. Heritabilities were greater than 50% for all three diseases, and the second rating had slightly higher heritability than the first rating for all diseases (increases of less than 0.1 to 3.8 percentage points). The heritability of average disease scores across ratings was highest for all diseases (0.60 for GLS, 0.53 for NLB, and 0.78 for SLB), but only slightly higher than the heritability of second ratings (Table 3). The genotypic correlation of ratings across time points was very high for all three diseases (0.92 to 0.98), indicating strong consistency of repeated ratings.

**Table 3.** Broad-sense heritability estimates on a line mean-basis (H) for each of two ratings and for the mean across ratings, and the genotypic correlation (r_g_) between ratings for gray leaf spot (GLS), northern leaf blight (NLB), and southern leaf blight (SLB) diseases measured on NILs at three environments each.

| Trait | H rating 1 | H rating 2 | H mean | $r_g$ between ratings |
| --- | --- | --- | --- | --- |
| GLS | 0.56 | 0.59 | 0.60 | 0.92 |
| NLB | 0.51 | 0.51 | 0.53 | 0.98 |
| SLB | 0.76 | 0.77 | 0.78 | 0.96 |

The NIL means for the three traits had low positive correlations (*r* = 0.20 for GLS and NLB, *r* = 0.36 for GLS and SLB, and *r* = 0.19 for SLB and GLS). The largest absolute values of correlation between disease traits and days to anthesis was 0.092, so we did not adjust disease means for differences in flowering time. Whereas flowering time and maturity tend to be strongly correlated with disease resistance scores in diversity panels with large ranges in maturity (Wisser *et al*., 2011), near-isogenic line population encompass much less variation in flowering time, reducing the confounding of maturity and disease resistance.

By chance we expect 63 of the 1264 phenotyped NILs to differ significantly from B73 for a trait at p < 0.05. We observed 106, 231, and 123 lines significantly different from B73 for GLS, SLB, and NLB, respectively (Figure 3). The largest tail of extreme values was a set of 173 NILs with significantly lower SLB scores (i.e. more resistance) than B73 (Figure 3).

**Figure 3.**
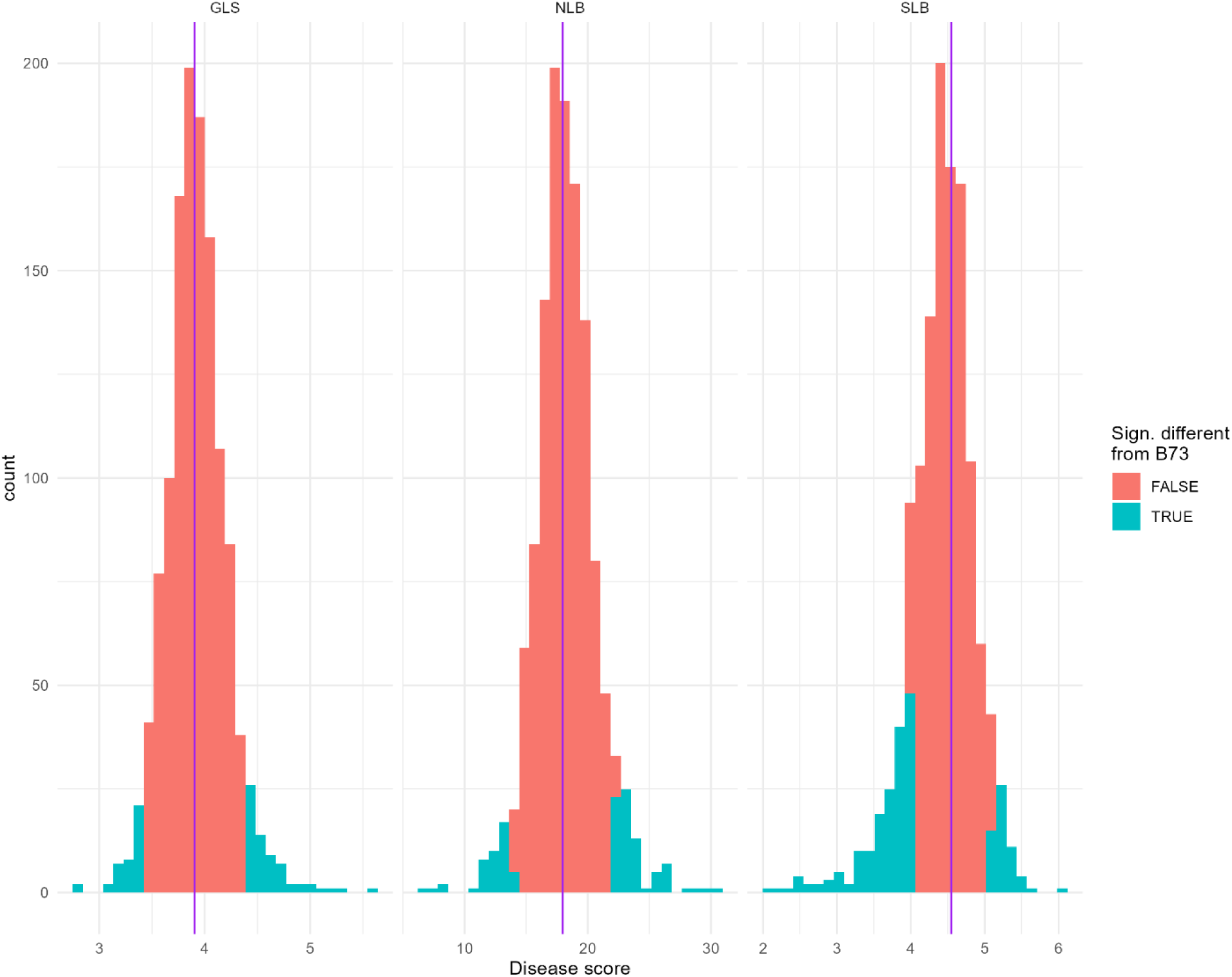
Histograms of mean disease scores (BLUEs) of nNILs for GLS, NLB, and SLB. Blue shaded regions represent nNIL means significantly different (p < 0.05) from the recurrent parent, B73. B73 mean value is indicated with a vertical line. GLS and SLB plots were scored on a 1-9 scale, where 1 corresponds to most resistant and 9 denotes most susceptible NLB plots were scored for 0-100% diseased leaf area with 5% increments

Three nNILs had significantly (p < 0.05) greater disease resistance (i.e less disease and lower disease scores) than B73 and another three NILs had significantly less disease resistance than B73 for all three diseases (Table 4). We term this multiple disease resistance or susceptibility (Wiesner-Hanks and Nelson, 2016). Two of three NILs with greater multiple disease resistance shared an introgressed region from donor CML103 on chromosome 8 (106.96 to 150.11 Mbp). Two of three NILs with greater multiple disease susceptibility shared a small introgressed region on chromosome 3 (9.87 to 14.53 Mbp) but from two different donors (CML247 and CML322)

**Table 4.** nNILs with significantly lower or higher disease scores for all three diseases (GLS, NLB, and SLB) compared to recurrent parent B73, their donor parents (based on sequence matching in introgressed regions) proportion of genome introgressed from donor parent, and introgression block chromosome and B73 reference genome 4.0 positions (in Mbp). GLS and SLB plots were scored on a 1-9 scale, where 1 corresponds to most resistant and 9 denotes most susceptible NLB plots were scored for 0-100% diseased leaf area with 5% increments.

| Line | Donor | GLS score | NLB score | SLB score | % introgression | Introgression chr (Mbp start, end) |
| --- | --- | --- | --- | --- | --- | --- |
| B73 |  | 3.9 | 18.0 | 4.5 | 0 |  |
| *Lines with significantly lower means (greater resistance) for all three disease traits* |  |  |  |  |  |  |
| CML103/B73_NIL-1008 | CML103 | 3.3 | 13.3 | 3.4 | 10 | 1 (0.00, 1.07)<br>1 (63.75,267.32)<br>7 (177.45, 182.38)<br>10 (139.50, 143.57) |
| CML103/B73_NIL-1216 | Tzi8 | 3.4 | 12.0 | 3.9 | 3 | 8 (106.96, 169.37)<br>9 (125.13, 129.61) |
| CML103/B73_NIL-1324A | CML103 | 3.3 | 13.3 | 3.3 | 6 | 3 (224.37, 229.15)<br>5 (65.92, 66.03)<br>8 (0.00, 3.86)<br>8 (23.57, 150.11) |
| *Lines with significantly greater means (less resistance) for all three disease traits* |  |  |  |  |  |  |
| CML247/B73_NIL-1020 | CML247 | 5.3 | 23.2 | 5.4 | 1 | 3 (6.25, 14.53)<br>5 (209.98, 210.10)<br>7 (179.28, 182.38) |
| CML322/B73_NIL-1018A | CML322 | 4.9 | 22.9 | 6.0 | 9 | 3 (9.87, 174.78)<br>4 (7.41, 24.65) |
| M162W/B73_NIL-1201 | M162W | 4.5 | 24.2 | 5.1 | 2 | 4 (193.27, 234.34)<br>9 (0.00, 4.67) |

### QTL mapping

We performed three different tests (described in the materials and methods section) for the effects of introgressions on the disease traits as described in the materials and methods; An “extreme NIL differentiation test”, a “common QTL test” and a “variable donor QTL test”. The extreme NIL differentiation test is significant when introgressions at a marker are enriched in nNILs significantly different from B73. The common QTL test estimates the mean effect of all nNILs with introgression at a marker compared to nNILs without introgressions. The variable donor QTL test estimates unique effects of each donor allele at each QTL relative to the B73 allele within the nNILs.

Fewer than 1% of markers were significantly (Bonferroni corrected *p* < 0.05) enriched in the tail of nNILs with GLS means significantly less than B73, whereas 6% and 4% of markers were significantly enriched in the more resistant phenotypic tails for NLB and SLB. Notable large regions of significant marker enrichment in more resistant tails were observed on chromosomes 1 and 8 for NLB and chromosomes 1, 3, and 8 for SLB (Figure 4). Near the end of chromosome 3, we identified markers enriched for association with the less resistant tail for SLB. The extreme NIL differentiation tests identify markers based on differences between nNILs and B73, whereas the other QTL tests identify trait-associated markers based on their associations within the entire nNIL set. Nevertheless, the same genome regions with extreme QTL peaks were also identified in the QTL scans for common donor allele effects and for variable donor allele effects (Figure 4). The variable donor effect model identified more QTL at higher significance levels than other models.

**Figure 4.**
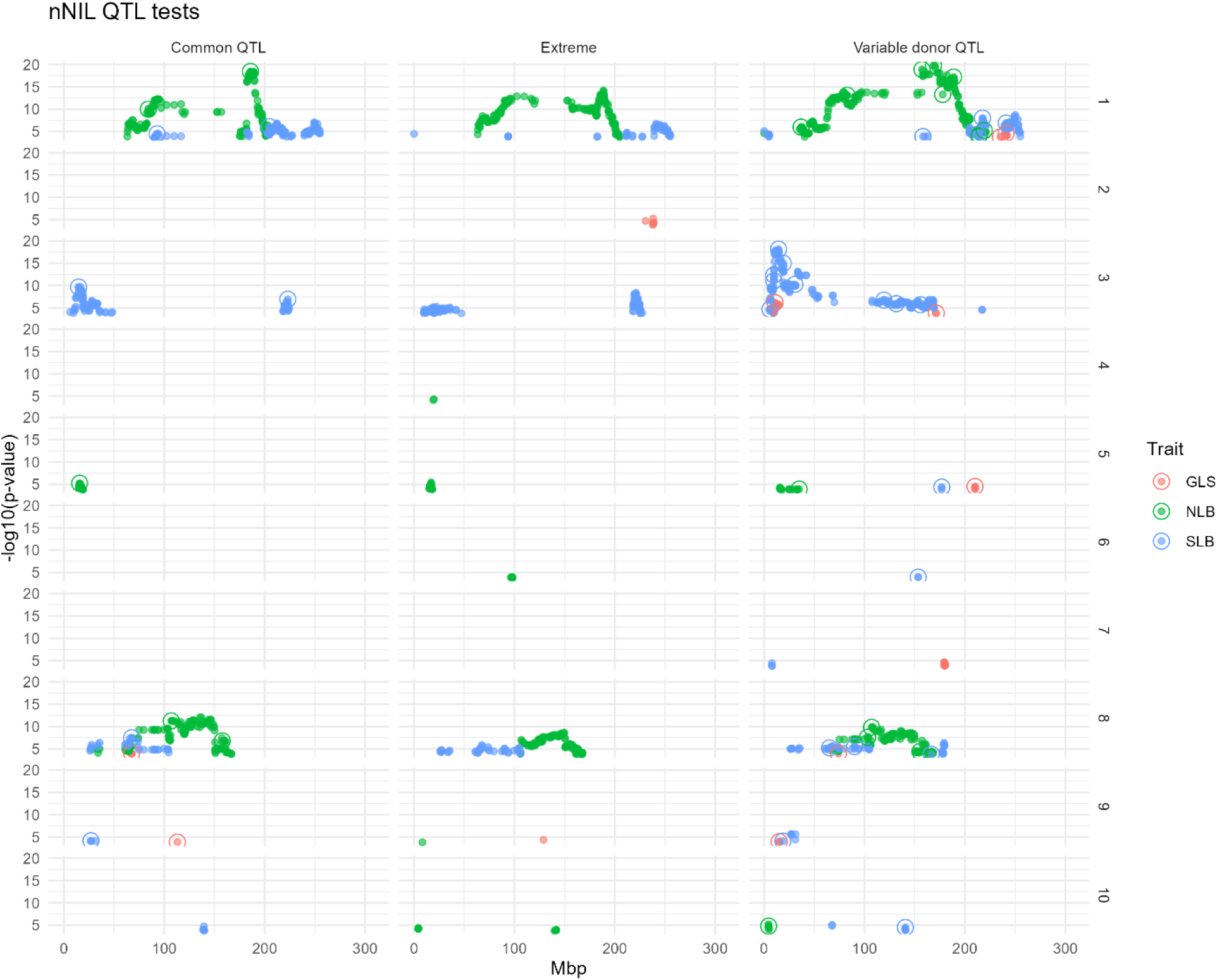
Significant QTL tests for nNILs on three disease traits (GLS, NLB, and SLB) using three different analysis approaches: “Common QTL”, “Extreme NIL differentiation” and “Variable donor QTL”. The Common QTL test estimates the mean effect of all nNILs with introgression at a marker compared to nNILs without introgressions. The Extreme test is significant when introgressions at a marker are enriched in nNILs significantly different from B73. The Variable donor QTL test estimates unique effects of each donor allele at each QTL. Numbers at the right of the figures indicate chromosome numbers

For the common and variable donor QTL tests, we then used model selection to obtain a final model in which markers were retained if they remained significant while other retained markers were included in the model. The final model fits were used to estimate the effects of all retained markers simultaneously. The final multiple QTL models based on the common donor allele effect test for GLS, NLB, and SLB included, respectively, 3, 5, and 6 QTL (Table S3), associated with 5, 22, and 19% of the variation among nNIL means. The final QTL models, based on the variable donor allele effect test for GLS, NLB, and SLB, included, respectively, 7, 11, and 11 QTL associated with 28, 44, and 57% (Table S4) of the phenotypic variation for GLS, NLB, and SLB, respectively.

## Discussion

In this study we describe a set of 1,264 NILs, of which we have genotypic data for 884, developed from crosses between 18 diverse inbred lines and the recurrent parent B73. We demonstrate their utility for QTL detection and the characterization of diverse allelic effects at QTL.

A publication detailing the genotypic and phenotypic characterization of this population was published previously (Morales *et al*., 2020). However, upon subsequent work with some of the individual nNILs, we noted considerable discrepancies between the actual positions of the introgressions and those reported in this paper. Ultimately, we determined that these discrepancies were substantial enough to warrant retraction of the original publication (Morales *et al*., 2024). For the current study we used the identical GBS genotypic and phenotypic data sets. However, we used a different approach to identify the introgressions that did not use imputed data or assume that the reported pedigrees and identities of the donor parents were accurate. In addition, we generated independent genotypic data for 24 of the nNILs and all 19 parents using a SNP genotyping chip. We used these SNP chip data to validate the original GBS dataset and to optimize our introgression calling pipeline.

For 21 of the 24 nNILs for which both GBS and chip genotypic data were available, the introgressions identified were nearly identical (Figure 1). The reason for the discrepancies seen in the other 3 lines are likely a combination of within-line segregation and genotyping with low coverage GBS. First, we originally received BC_5_F_4_ seed, which presumably derived from a single BC_5_F_3_ plant, in which we would expect that almost 40% of the introgressions to be heterozygous and therefore to be segregating in a line derived from it by bulking seed from progeny plants. The DNA used to derive the GBS and the chip genotypic data were from different plants, separated by at least one additional generation of sib-mating. Thus, it is quite likely that in some cases, different sublines had been generated that were fixed for different introgressions. The fact that, even in the three lines that showed substantial discrepancies, some introgressions identified by GBS and chip data were identical (Figure 1), is consistent with this explanation. Second, in some cases, we may have experienced difficulty in distinguishing heterozygous from homozygous introgressions due to low coverage GBS sequence data.

Another potential problem with our current results are the small introgressions of less than 10 Kb that make up 4.7% of all the introgressions identified in nNILs matching their putative donors (and 4.8% in the whole set of genotyped nNILs). The probability of two recombination events occurring so close to each other during the breeding process is very low. Furthermore, in some cases, these small introgressions are shared among multiple nNILs, which would be very unlikely if they were true introgressions. For example, 9 nNILs representing 3 different donors all have introgressions less than 5 kb in length on chromosome 7 around position 3743350 bp and 4 nNILs representing a different group of 4 donors all have 21 bp introgressions on chromosome 4 position 12740845 (File S6). We have retained these introgressions in our datasets in case they are valid and useful, however we suspect that many of them are false positive introgressions that may have arisen from incorrect alignments of short sequencing reads to the reference genome, leading to incorrect signals of introgression that would tend to be clustered in specific genomic regions. It is also possible that introgressions less than 2 kb in size resulted from gene conversions during the breeding process, which were observed at a rate of about 30 gene conversions per recombinant inbred maize line studied by Liu et al. (2018)using about 3-fold higher marker density than we used in this study.

Because of the independent segregation of sublines and the potential problems with small introgressions, we recommend that researchers interested in using this population independently validate introgressions of interest, especially if the introgressions are less than 10 kb. If the desired introgression is not found it may be present in either homozygous or heterozygous state in other individuals of that same line. The phenomenon whereby subfamilies differing for specific introgressions can be differentiated from mostly inbred lines has been described as “heterogeneous inbred families” (HIFs) and can be usefully exploited for higher-resolution mapping and characterization of QTL (Tuinstra *et al*., 1997).

Mapping introgressions using low coverage short read sequences was challenging in this case because of uncertainty introduced from alignment errors, high missing data rates, and uncertainty about the actual pedigrees of the lines. A previous analysis (Morales *et al*., 2020) dealt with the high missing data rate problem by first imputing the raw GBS SNP calls using FILLIN imputation algorithm based on matching small genome segments to a library of diverse maize lines (Swarts *et al*., 2014) . In this case, this imputation method is likely to introduce errors because the donor haplotype may not be the most frequent haplotype that matches the sparse genotype calls in a particular genome region. For some uses, the error rate of this imputation method may be acceptable, but for the current study, initial imputation errors can result in significant subsequent errors in mapping introgressions. Our method avoids this initial imputation step entirely and is robust to high missing data rates.

A previous analysis of a related set of nNILs from the same source population (Kolkman *et al*., 2020) used a population of 412 nNILs to characterize QTL for NLB resistance. While this population was received by a source distinct from the population described here, it was ultimately also derived from Syngenta and, of the 412 NILs used in this study, 51 were named identically to lines in the population described here. Our analyses suggest that while these lines share the same names, the introgressions they carry differ substantially between the two studies. This is likely in significant part due to independent segregation of heterozygous regions discussed above. Kolkman et al. (2020) received the lines at the BC_3_F_3_ generation while the seed we received was at BC_3_F_4_ and in each case subsequent selfing was performed to bulk the seed. Additionally, distinct methods to call introgressions were used in the two studies and Kolkman et al (2020) also used the FILLIN imputation algorithm which may have introduced some errors as detailed above. We recommend therefore that similarly-named lines in these two populations be regarded as distinct.

Similar to Kolkman et al (2020), we found that the introgressions in a substantial number of the lines did not match their reported pedigrees; 20% of lines were matched with high probability to a NAM parental line that disagreed with their pedigree, a further 11% of lines did not match the reported donor parent with certainty and we were not able to make a determination for 2% of lines that had only very small introgression blocks. These mismatches were not accounted for in the previous analysis of this population (Morales *et al*., 2020), which may account for some of the inaccuracies previously observed. Kolkman et al (2020) dealt with uncertainty about the donor parents by comparing each nNIL’s graphical genotype to the recurrent and putative donor parent to check if the donor parent was correct, and by comparing whole genome SNP scores to the 18 possible donor parents to identify a best match. Our method differs in that matches to donor parents are only calculated within called introgression blocks, ignoring most of the genome that is derived from B73 and is not informative about donor parentage. Knowing the donor parent genotype with certainty would make the introgression calling procedure much easier, as the marker set can be reduced to the informative markers for each pedigree. If donor and recurrent parent genotypes are known with certainty, introgression mapping methods that account for sequencing read coverage in each sample can improve the resolution of introgression boundaries (Campos-Martin *et al*., 2023). However, such methods are unreliable if there are errors in the donor parent; our introgression calling method does not require knowledge of the donor parent genotype.

Due to the low frequency of the minor alleles in NIL populations, genetic mapping using NIL populations of this type has lower power for QTL detection compared to population structures with more balanced allele frequencies, such as recombinant inbred line (RIL) populations (Jamann *et al*., 2015). Nevertheless, this type of mapping has been performed many times (e.g. Szalma *et al*., 2007, Lennon *et al*., 2016, Lennon *et al*., 2017). We mapped QTL for SLB, NLB and GLS resistance in the nNIL population. We used three different approaches to identify QTL; An “extreme NIL differentiation test”, a “common QTL test” and a “variable donor QTL test”. The extreme NIL differentiation test, which is conceptually similar to approaches described previously (Deng *et al*., 2000, Wolyn *et al*., 2004), identified introgressions which disproportionately occurred in lines that were at the extremes of the phenotypic distributions in line that were significantly different from B73 . The common QTL test made the assumption that all non-B73 alleles conferred a similar effect relative to the B73 allele. The variable donor QTL test estimated unique effects of each donor allele at each QTL relative to the B73 allele. The variable donor allele effect model appears to be the more realistic approach, as in previous work with the NAM population we noted allelic series at QTL for resistance to these diseases with substantial variation in allelic effects (Kump *et al*., 2011, Poland *et al*., 2011, Benson *et al*., 2015). In agreement with these previous studies, we observed substantial variation in allelic effects at a given QTL among donor alleles, providing more power to explain the observed variation (although also using many more model parameters). The final variable donor effect QTL model also included several clusters of linked markers, such as on chromosome 1 for GLS (Figure S6) and chromosome 3 for NLB (Figure S7). In these cases, the markers tagged different sets of introgressions (although some were shared), potentially capturing the effects of different donors distributed among clusters of linked QTL). This analysis highlights one of the most important potential uses of the nNIL population; its potential use to isolate and characterize varying allelic effects and mechanisms at specific QTL.

The QTL identified are, for the most part, at similar loci to QTL previously identified for these traits. For example, the SLB QTL identified at 14.3 Mbp and 18.7 Mbp (Table S4) are close to SLB resistance gene *ChSK1* (*Zm00014a001903*) (Chen *et al*., 2023) which is at 16,572,093-16,578,681 bp (Mo17-2021) on chromosome 3. A major SLB QTL previously detected in NAM population between 51.8 and 66.9 Mpb on chromosome 8 (Balint-Kurti *et al*., 2007) is also identified in this analysis (Table S4).

Similarly, the *ZmWAK-RLK1* conferring resistance to NLB is located at 156.1 Mb on chromosome 8 (Hurni *et al*., 2015, Yang *et al*., 2021) close to the effect detected at 155.1 Mbp (Table S4) and another NLB resistance gene has been located around 187 Mbp on chromosome 1 (Jamann *et al*., 2014), close to the associated SNP at 188.7 Mbp (Table S4).

In conclusion, we believe that this nNIL population will be of substantial value for identification and characterization of QTL in maize and, particularly for the characterization of allelic series with variable effects.

## Methods

### Population development

We requested 1,264 nNILs from the greater Syngenta AG (Basel, Switzerland) panel from CIMMYT, Mexico. Their reported pedigrees suggested that these had been derived from crosses between one of 18 diverse inbred lines (CML103, CML228, CML247, CML277, CML322, CML333, CML52, CML69, Ki11, Ki3, M162W, Mo17, Mo18W, NC350, NC358, Oh43, Tx303, and Tzi8) acting as the donor parent, and the recurrent inbred parent B73 (Gandhi *et al*., 2008). All 18 donor parents of this population are among founders of the widely-used maize nested association mapping (NAM) population (Gage *et al*., 2020). The pedigrees further suggested that the lines had been backcrossed to B73 for five generations (BC_5_) and that they were then self-pollinated for four generations (BC_5_F_4_). nNILs reported to carry introgressions from the same donor were considered a subpopulation, with each subpopulation comprising 50-108 BC_5_F_4_ nNILs (Table 2). The introgression donors were of tropical, temperate non-stiff stalk, and mixed origin, while B73 is a temperate stiff stalk line (Romay *et al*., 2013). Seed for the 1,264 nNILs can be requested from the corresponding author or from the Maize genetic stock center.

### Field design and inoculation

The entire population was evaluated for GLS, NLB, and SLB severity across four separate year/field replication environments for each disease. For each experiment, we used an augmented incomplete block design, in which each subpopulation was grown separately with 25-plot blocks. nNIL plots were replicated once per environment and randomized within subpopulation. One plot of B73, the recurrent parent, was randomized within each 25-plot block. The NLB and SLB experiments were conducted at the Central Crops Research Station in Clayton, NC. NLB was evaluated in 2015 and 2018 with one replication per year and in 2016 with two replications. SLB was assessed in 2014 with two replications and in 2015 and 2016 with one replication per year. GLS was screened in Andrews, NC in 2014 with one replication and at College Farm Research Station in Blacksburg, VA in 2016 with two replications and in 2017 with one replication.

Inoculum for the GLS (Blacksburg, VA only), NLB, and SLB experiments was prepared from mixtures of isolates of *Cercospora zea-maydis*, *Setosphaeria turcica* (race 0, race 1, race 2,3 and race 2,3,N), and *Cochliobolus heterostrophus* (including 2-16Bm and Hm540), respectively, as previously described (Sermons and Balint-Kurti, 2018, Martins *et al*., 2019). The Andrews, NC site has naturally high and consistent levels of *C. zea-maydis* inoculum present in soil crop residues, which allowed for evaluation of GLS under natural infection.

### Phenotyping

In all SLB experiments and in the 2015 NLB experiment, we took observations on flowering time 2-3 times per week and estimated plot anthesis dates as the first date on which 50% of plants had shed pollen. Ear and plant height were measured in each plot in all SLB experiments and in the 2015 and 2018 NLB experiments (File S7). Plant height was measured as the height to the base of the topmost leaf node. Ear height was measured as the height to the base of the topmost ear. Each plot was visually evaluated for disease severity at two time points per experiment: GLS on July 28 and August 6 in 2014, August 9 and 19 in 2016, and August 9 and 25 in 2017; NLB on July 13 and 22 in 2015, July 17 and 21 in 2016, and July 18 and 26, 2018; SLB on July 14 and 31 in 2014, July 10 and 25 in 2015, and July 11 and 20, 2016 (File S7). GLS and SLB plots were scored on a 1-9 scale, where 1 corresponds to most resistant and 9 denotes most susceptible. This scale is described in Sermons and Balint-Kurti (2018), although here we have reversed the scales described in that publication. NLB plots were scored for 0-100% diseased leaf area with 5% increments. The two disease scores were averaged for each plot that was scored twice. Five nNILs (CML333/B73 NIL-1002, CML333/B73 NIL-1007, Ki11/B73 NIL-1103, Ki11/B73 NIL-1104, Ki3/B73 NIL-1258) exhibited severe lesion-mimic mutant phenotypes (Kim *et al*., 2020) and thus were not included in further analyses on disease resistance.

### Phenotypic data analysis

To estimate genotypic variances and correlations for multiple ratings for a single disease, we fit a linear mixed model to the individual time point data (File S8):

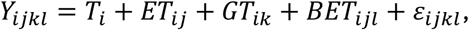

Where *T*_*i*_ is the fixed intercept for the *i*th rating time point (*i* = 1 or 2 for first or second rating time within an environment), *ET*_*ij*_ is the fixed interaction between environment *j* and rating time *i*, *GT*_*ik*_ is the random effect of genotype *k* at rating time *i*, *BET*_*ijkl*_ is the random effect of block *l* in environment *j* and time *i*, and *ε*_*ijkl*_ is the residual effect. Mixed models were fit using the asreml-R package (Butler *et al*., 2017) in R (R Core Team, 2013). We tried to include an interaction term for genotype-by-environment-by-time interaction, but that model failed to converge.

The covariance structures for random effects were:

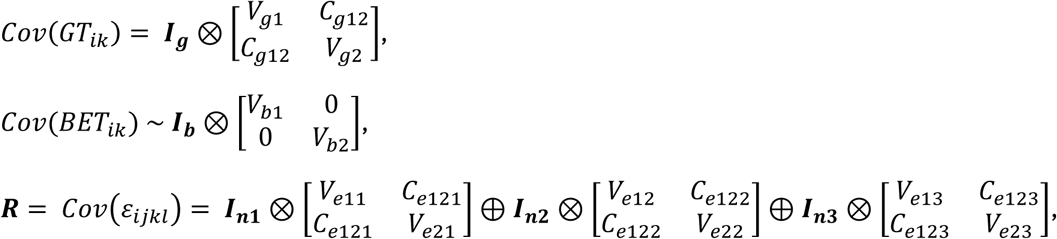

Where ***I**_g_*, ***I**_b_*, and ***I**_ni_* are identity matrices with dimensions equal to the number of genotypes (*g*), the number of blocks (*b*), and the number of plots in environment *i (ni*), respectively; *V*_*g*1_ and *V*_*g*2_ are genotypic variance components at rating times 1 and 2; *C*_*g*12_ is the genotypic covariance between rating times 1 and 2; *V*_*b*1_ and *V*_*b*2_ are the block variances for rating times 1 and 2; *V*_*e*1*i*_ and *V*_*e*2*i*_ are residual variance components at rating times 1 and 2 in environment *i*. ⊕ represents direct sums of matrices and ⊗ represents direct products of matrices. Genotype predictions averaged across environments were made for each rating time separately and also averaged across rating times. Heritability of each rating and of the mean across ratings was estimated as the mean of the reliability of the genotype predictions (Isik *et al*., 2017). The genotypic variance of the mean rating was estimated as 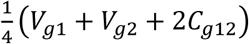.

We also estimated the adjusted means (best linear unbiased estimators) of nNILs across ratings and environments to use for subsequent quantitative trait locus (QTL) testing (Holland and Piepho, 2024) using the following linear model:

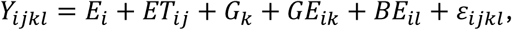

Where terms are defined analogously to the previous model, but genotype main effects (*G*_*k*_) are fit as fixed effects and all other terms were fit as random effects with independent and identical variance structures. We estimated the adjusted genotype means and the matrix of standard errors of their pairwise differences. We then estimated the mean of pairwise differences between B73 and each nNIL and used this difference to determine if an nNIL mean differed from the B73 mean by more than expected by chance at α = 0.05.

### DNA extraction and sequencing

Three seeds per nNIL were planted in a greenhouse at the Cornell University Kenneth Post Laboratory. Approximately one month after germination, 25-50 mg of seedling leaf tissue was collected from each nNIL. Genomic DNA was extracted from fresh leaf tissue using the DNeasy Plant Mini Kit system (Qiagen, Inc., Valencia, CA, USA). GBS (Elshire *et al*., 2011) was performed on 884 unique nNIL DNA samples by BGI Americas Corporation (Cambridge, MA, USA) using 100-base pair paired-end sequencing with the HiSeq 4000 system (Illumina, Inc., San Diego, CA, USA).

### Genotyping-by-sequencing analysis

For proprietary reasons, the BGI team clipped the barcodes on the raw read fastq files before delivering the data to us. In order to use the TASSEL-GBS pipeline (Glaubitz *et al*., 2014) to call single nucleotide polymorphisms (SNPs) on our nNIL panel, we modified the TASSEL 5 (Bradbury *et al*., 2007) base code to bypass the requirement for barcodes. In addition, the TASSEL-GBS pipeline (Glaubitz *et al*., 2014) only accepts single-end reads. In order to meet the single-end read requirement, we generated the reverse complement of the second paired-end fastq file for each set of paired-end fastq files with the FASTX-Toolkit (Hannon, 2009) and then merged the two fastq files.

We made a GBS build from 4,603 lines from the USA maize inbred seed bank that were previously analyzed with GBS by the Panzea Maize Diversity Project (Zhao *et al*., 2006). We accessed the Panzea Maize Diversity Project GBS fastq files (NCBI BioProject accession #PRJNA200550) from the public NCBI Sequence Read Archive (Leinonen *et al*., 2010). The Panzea Maize Diversity Project GBS fastq files were processed with the GBS Discovery Pipeline in TASSEL 5 (Bradbury *et al*., 2007, Glaubitz *et al*., 2014) with the B73 reference genome 4.0 (NCBI BioProject accession #PRJNA10769), a k-mer length filter of 64, and a minimum quality score of 20. The TASSEL GBS Discovery Pipeline yielded a GBS database, which we then used to call SNPs on the Panzea Maize Diversity Project and nNIL fastq files with the GBS Production Pipeline in TASSEL 5 with a k-mer length filter of 64 and a minimum quality score of 20 for the Panzea Maize Diversity Project lines and 10 for the nNILs. This resulted in 564,764 SNPs. The raw GBS data set was filtered to remove fixed SNPs and markers with more than 20% missing data, leaving 93,541 markers (Files S1 and S9). Duplicated lines, and SNPs and lines with more than 2% heterozygosity were then removed, resulting in 64,141 SNPs scored on 884 lines (Files S10).

### Genotyping using a high-density SNP array

As a quality control check, we regenotyped a subset of 24 nNILs and all 25 of the NAM founder lines using a 60K Axiom™ Maize6H Genotyping Array (ThermoFisher cat# 551099) run at North American Genomics (File S2). The DNA used for chip assays of all the mentioned materials was derived from a single plant and isolated using a NucleoSpin PlantII kit (Macherey-Nagel cat# 740770.250). The 24 lines were selected based on their extreme phenotopyes for SLB resistance (both resistant and susceptible) as this was a trait of particular interest.

### Introgression calling

We developed a new pipeline for calling introgressions that did not rely on knowing the donor parent of each nNIL and applied it separately to the original GBS SNP call data and to the newly acquired chip SNP call data for the subset of 24 nNILs genotyped with both methods and checked for agreement between the two data sets.

The new introgression calling pipeline was based on a hidden Markov model (HMM) algorithm that assumes three possible true identity-by-descent (IBD) haplotype states at each SNP position: homozygous B73 haplotype (0), heterozygous (1), or homozygous donor haplotype (2). The starting frequencies for each state were based on the expectations for genotype frequencies in BC5F3:4 introgression lines: 0.011179 homozygous for donor alleles, 0.007813 heterozygous, and 0.981008 homozygous for B73 alleles.

The transition matrix between states was based on the average genome-wide recombination frequency expected between adjacent markers for a set of *m* markers and a total 1500 cM linkage map distance for maize (similar to the mean linkage map length of 23 maize populations: Bauer *et al*., 2013) over the entire process of nNIL development. The recombination frequency over all generations is expected to equal twice the average recombination rate because there are the equivalent of two outbreeding meiotic generations between the F1 and the final backcrossed and selfed progenies. We assumed the probability of double recombinations between adjacent markers was zero. Therefore, we modeled average recombination rates around 2*1500/(100\**m*).

The probability of a transition from true state 0 to true state 1 between adjacent markers was computed as the probability of recombination between adjacent markers times the conditional probability of heterozygosity at the second locus (Table S1). If a recombination occurred between a homozygous recurrent parent locus (state 0) and the adjacent locus, the adjacent locus must be either heterozygous (1) or homozygous donor parent (2) with relative frequencies equal to the expected genotypic class frequency divided by the sum of the heterozygous and homozygous donor class frequencies: (f_het/(f_het + f_homoD) or (f_homD/(f_het + f_homoD). The probability of transitions from true state 1 to either true state 0 or true state 2 was modeled as half the probability of a recombination between adjacent markers because if there is a recombination between two heterozygous loci in a previous generation, the resulting progeny will have either of two homozygous classes with equal frequency at the second locus. The probabilities of transition from true state 2 to other states were modeled as equal to the reciprocal transition probabilities.

Observed states correspond to identity-in-state (IIS) SNP genotypes and were modeled as homozygous for B73 allele (0), heterozygous (1), homozygous for donor allele (2), or missing (state 3; File S10). The emission probability matrix relating true haplotype states to observed allele states involved five parameters (Table S2): missing data rate (*mr*), non-informative rate (*nir*), genotyping error rate for true homozygotes (*ger_m_*), genotyping error rate for true heterozygotes (*ger_t_*), and the proportion of homozygotes mis-called as heterozygotes (*p*) given a genotyping error. Missing data rate was estimated from the data as the proportion of missing SNP calls in the data set averaged over all individuals and loci. Non-informative rate represents the probability that a donor parent’s SNP allele is identical in state to the B73 allele. Genotyping error rates result from sequencing and alignment errors. We modeled genotyping error rates separately for true homozygous and heterozygous SNP genotype classes because these can vary in low-coverage sequencing data. Further, the probabilities that a genotyping call on a true homozygote results in incorrect heterozygous or homozygous calls are not necessarily equal, as they also depend on sequencing coverage. Therefore, we allowed the parameter *p* to represent the proportion of true homozygous calls resulting in heterozygous mis-calls. We used genome-wide averages for all of these parameters, which may not be correct for any specific pair of loci, but greatly simplifies implementation of the HMM.

We implemented the HMM in Python version 3.7.6 with the hmmlearn version 0.3.2 module (https://hmmlearn.readthedocs.io/en/latest/). The chip SNP data included 24,070 markers with physical positions provided in AGPv3 coordinates (Files S2 and S11). Chip SNP positions were uplifted to AGPv4 positions using a bed file with the AGPv3 positions (File S11) and the assembly converter tool at https://plants.ensembl.org/Zea_mays/Tools/AssemblyConverter (Cunningham *et al*., 2015), dropping SNPs lacking autosomal positions on AGPv4 (File S12). The filtered chip marker set included 20,221 markers with very low missing rate (< 0.003). We performed a grid search around HMM parameters, including the non-informative values of 0.001, 0.01, 0.1, 0.3, 0.5, 0.7, and 0.9; genotype calling error rates (both ger_m_ and ger_t_) of 0.0001, 0.001, and 0.01; proportion of homozygous calling errors that result in heterozygous calls of 0.1, 0.25, 0.5, 0.75, and 0.9; and recombination rates equal to the average adjacent marker pair recombination rate times 0.5, 1, or 2. The grid search was implemented using data for B73, donor parents, and the 24 NILs (File S13). The proportions of homozygous and heterozygous introgressions were estimated for B73, donors, and NILs separately (File S3). The parameter settings resulting in the highest proportion of homozygous introgression calls among the donor parents (0.981) while calling zero introgressions on the B73 parent was a non-informative rate of 0.9, genotype call error rate on homozygotes (*ger_m_*) of 0.01 and on heterozygotes (*ger_t_*) of 0.001, *p* = 0.1, and half the average recombination rate (Files S3 and S14). These parameter settings were used to make the introgression calls because results were closest to expectations. Using these settings, the nNILs had average proportion of 0.050 markers called as homozygous introgressions and 0.003 markers as heterozygous. To compare these chip-based introgression calls to GBS-based introgression calls we identified the GBS marker closest to each chip marker and removed chip markers that were associated with redundant closest GBS markers, resulting in a data set with introgression calls projected to 11,326 SNPs from the GBS data set.

We tested the HMM on GBS data of the same 24 NILs analyzed with the chip data set (File S15). The mean missing data rate on this set was 0.086. We performed a grid search over the same non-informative rate and SNP calling error rates as used for the chip SNP data. The expected mean adjacent marker pair recombination rate was recalculated for GBS data to account for the higher number of markers. We evaluated introgression calling using 0.5, 1, or 2 times this mean recombination rate. The GBS-based introgression calls were compared to the chip-based introgression calls projected to the nearest GBS SNP. Results did not vary dramatically among widely different parameter settings, but the proportion of homozygous introgression calls increased as the non-informative rate parameter increased (Files S4 and S16). The parameter setting of 0.9 for non-informative rate, ger_m_ = 0.01, ger_t_ = 0.0001, *p* = 0.9, and average recombination rate between marker pairs (*r* = 0.00047) resulted in the closest agreement between GBS and chip introgression calls (mismatch rate of 0.0007; File S4; Figure S2). At these settings, the GBS data indicated homozygous introgression call rate of 0.044, and heterozygosity rate of 0.003 (File S4); the homozygous introgression proportion was slightly lower than estimated with the chip data (File S3). The mismatch rate between chip and GBS introgression calls was robust to parameter settings (since a very high proportion of calls are B73 homozygous no matter what settings are used), but the proportion of introgression calls was more sensitive to the non-informative rate, increasing from 0.014 to 0.044 as the non-informative rate increased from 0.001 to 0.09 (File S4). These parameter settings were then used to call introgressions on the full set of 884 genotyped nNILs (Files S15 and S17).

Next, we compared SNP calls within introgression regions to compare each NIL to each of the NAM founder inbreds (Files S17 and S18). We used unimputed HapMap 3.2.1 data on the founders (Bukowski *et al*., 2018) to make these comparisons. HapMap data were obtained from Cyverse data commons (Swetnam *et al*., 2024) and projected from B73 AGPv3 to AGPv4 positions. We filtered the HapMap data set to include B73 and donor parents (by pedigree) of the NILs in the testing set. For each data set on the common set of 24 nNILs genotyped both by GBS and chip, we identified markers in common between the NIL data set and the HapMap: 17,548 for chip data and 36,609 for GBS. For each of the subset of 24 NILs, the proportion of its SNP calls identical to each founder SNP calls within introgression segments called using the chip data, ignoring heterozygous calls, was computed (Files S18 and S19). Most comparisons revealed that the chip SNP calls within NIL introgression segments matched best to the donor parent indicated by the NIL pedigree, but several NILs matched more closely to different donor parents, and one had no obvious best match to a donor parent (File S19). We repeated this analysis with the GBS data (File S20) and obtained identical results (File S21); the NILs that matched to a different NAM founder than indicated by their pedigree with chip data matched the same founder with GBS data. We then used the test set of GBS data from the 20 nNIL cases where the best donor match agreed with the pedigree and computed the difference between the match proportion of the first and second best matches to the donors for each line and standardized the difference by the standard deviation of the match proportions for all of the NAM founders except the best match for each nNIL (File S22). Based on these results, we developed a rule that best matched NAM founders could be defined for each nNIL if at least 20 SNPs within introgression blocks were in common between a particular nNIL and NAM founder data sets, the highest match proportion was at least 80% of SNPs, and the difference between the maximum match proportion and the 2^nd^ highest match proportion was greater than 1.5 standard deviations of the match statistics (excluding the maximum match) for a particular line. We then applied this rule to the complete set of 884 nNILs genotyped with GBS, excluding duplicates (Files S22 and S5). For the purpose of estimating introgression block sizes, introgression breakpoints were estimated as the centers of the intervals between first and last contiguous markers and their adjacent flanking non-introgressed markers, respectively.

### QTL tests

We performed three different tests for the effects of introgressions on the disease traits: (1) Extreme NIL differentiation test, (2) Common QTL test, and (3) Variable donor QTL test (File S23) The Extreme NIL differentiation test is significant when introgressions at a marker are enriched in nNILs significantly different from B73. Common QTL test estimates the mean effect of all nNILs with introgression at a marker compared to nNILs without introgressions. Variable donor QTL test estimates unique effects of each donor allele at each QTL. For all tests, we pruned the introgression call data to remove redundant markers.

The extreme NIL differentiation tests was performed by first identifying phenotypically extreme NILs if their mean disease rating differed significantly from the B73 at α = 0.05, and considering positive and negative extreme tails separately. Then, the frequency of introgression at each marker in each tail was tested for deviation from expectations under random chance using Fisher’s exact test (Upton, 1992). We used an empirical estimate of the effective number of marker tests (Galwey, 2009) to define a Bonferroni-threshold for declaring significance for these single marker tests:

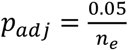, where *n*_*e*_ is the effective number of marker tests.

The common QTL test was based on association between mean disease values and introgressions assuming a common effect among introgression donor parents. We fit a linear model using the lm() function in R for each marker separately:

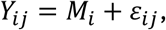

Where *Y*_*ij*_ is the adjusted mean of the *j*th nNIL with introgression call *i* (0,1,or 2) at the marker and *M*_*i*_

is the effect of the *i*th level of introgression dosage at the marker. We also performed forward

regression to select a set of markers to jointly model each disease trait with most significant marker added each step until no further markers could be added at p < 0.01.

The variable donor QTL test was a test of association between mean disease values and introgressions, allowing each donor parent to have a unique QTL effect. We fit a linear model using the lm() function in R for each marker separately:

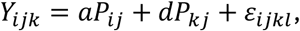

Where *Y*_*ijkl*_ is an adjusted nNIL mean, *aP*_*ij*_ is the additive regression effect of introgression (*i* = 0 or 1, or 2) from the *j*th donor parent P, and *dP*_*jk*_ is the dominance effect of introgression (*k* = 1 for heterozygotes, 0 for others) from the *j*th donor parent P. We performed forward regression to select a set of markers to jointly model each disease trait using Akaike’s Information Criterion (AIC: Akaike, 1974) to select the marker that most improved the model at each step until no further markers improved the AIC.

## Acknowledgements

This work was funded by NSF grants IOS-1127076 to PBK, RJN and JBH and IOS-2154872 to PBK, JBH and TJ. We thank Greg Marshall, Judith Kolkman, and Molly Towne for technical and logistical support, Denise Costich and the CIMMYT Germplasm Bank for providing the nNIL seeds, the staff at the Central Crops Research Station in Clayton, NC for field trial management, Corteva for field space at Andrews for the GLS field trials.

## Supplemental Tables

**Table S1.** Transition matrix for the Hidden Markov Model for calling introgressions in BC5F3:4 generation near-isogenic lines. R is the effective recombination rate over all generations of backcrossing and selfing. Our starting value was 2*1500/(100\**m*)) for a typical total maize linkage map distance of 1500 cM, *m* markers, and a factor of two to reflect about two effective meiosis during the backcrossing and selfing process. f_0 is the expected proportion of homozygous recurrent parent genotypes in the BC5F3:4 generation = 0.980469. f_1 is the expected proportion of heterozygotes in the BC5F3 generation (or, equivalently, segregating lines in the BC5F3:4 generation) = 0.007813. f_2 is the expected proportion of homozygous donor parent genotypes in the BC5F3:4 generation = 0.01179.

|  |  | Ending state |  |  |
| --- | --- | --- | --- | --- |
|  |  | 0 (B73 homozygous) | 1 (heterozygous) | 2 (donor homozygous) |
| Starting state | 0 (B73 homoz) | $1 - R$ | $R \times (f_1 / (f_1 + f_2))$ | $R \times (f_2 / (f_1 + f_2))$ |
| | 1 (het) | $R \times (f_0 / (f_0 + f_2))$ | $1 - R$ | $R \times (f_2 / (f_0 + f_2))$ |
| | 2 (donor homoz) | $R \times (f_0 / (f_0 + f_1))$ | $R \times (f_1 / (f_0 + f_1))$ | $1 - R$ |

**Table S2.** Emission probability matrix for the Hidden Markov Model for calling introgressions in BC5F3:4 generation near-isogenic lines. nir = non-informative rate (probability that a SNP allele in a donor haplotype is identical in state to the recurrent parent SNP allele). ger_m_ = SNP-calling error rate for true homozygotes (probability that a homozygous SNP genotype is called incorrectly). ger_t_ = SNP-calling error rate for true heterozygotes (probability that a heterozygous SNP genotype is called incorrectly). p = proportion of homozygous SNP call errors resulting in heterozygous call. If a homozygous genotype is mis-called, a proportion p is mis-called as heterozygous, whereas a proportion (1 - p) is mis-called as the wrong homozygous genotype. mr = missing rate (proportion of missing data). Initial value estimated as the average missing data proportion. Explanation of individual probabilities: **True state 0 called as SNP state 0** occurs with probability of no genotyping call error on a true homozygote (1-ger_m_) and not missing data (1 – mr). Note that the non-informative rate does not impact calls on true recurrent parent homozygotes. **True state 0 called as SNP state 1** occurs with probability of a genotyping call error on a true homozygote (ger_m_) times the proportion of homozygous genotype call errors that result in heterozygous calls (*p*) and not missing data (1 – mr). **True state 0 called as SNP state 2** occurs with probability of a genotyping call error on a true homozygote (ger_m_) times the proportion of homozygous genotype call errors that result in wrong homozygous calls (1-*p*) and not missing data (1 – mr). **True state 1 called as SNP state 0** occurs if not missing data (1 – mr) and either: the SNP is informative (1 – nir) but there is a genotyping call error with probability ger_t_ wherein 50% of genotyping call errors on heterozygotes result in SNP calls of state 0 (the other half result in SNP calls of state 2) OR the SNP is not informative (nir) and a there is not a genotyping call error on a homozygote (1-ger_m_). **True state 1 called as SNP state 1** occurs if not missing data (1 – mr) and either: the SNP is informative (1 – nir) and there is not a genotyping call error with probability ger_t_ (1 - ger_t_) OR the SNP is not informative (nir) but there is a genotyping call error on a homozygote (ger_m_) resulting in a heterozygous call (*p*). This second case involves two opposite errors that cancel out and result in a correct SNP call. **True state 1 called as SNP state 2** occurs if not missing data (1 – mr) and either: the SNP is informative (1 – nir) but there is a genotyping call error with probability ger_t_ wherein 50% of genotyping call errors on heterozygotes result in SNP calls of state 2 OR the SNP is not informative (nir) but there is a genotyping call error on a homozygote (ger_m_) resulting in wrong homozygous SNP call (1 –*p*). **True state 2 called as SNP state 0** occurs if not missing data (1 – mr) and either: the SNP is informative (1 – nir) but there is a genotyping call error on a homozygote (ger_m_) resulting in wrong homozygous SNP call (1 – *p*) OR the SNP is not informative (nir) and a there is not a genotyping call error on a homozygote (1-ger_m_). **True state 2 called as SNP state 1** occurs if not missing data (1 – mr) and either: the SNP is informative (1 – nir) but there is a genotyping call error on a homozygote (ger_m_) resulting in a heterozygous SNP call (*p*) OR the SNP is not informative (nir) but there is a genotyping call error on a homozygote (ger_m_) resulting in a heterozygous SNP call (*p*). The sum of these two cases is: [((1-nir)*ger_m_*p) + (nir*ger_m_*p)]*(1-mr) = ger_m_*p*(1-mr) **True state 2 called as SNP state 2** occurs if not missing data (1 – mr) and either: the SNP is informative (1 – nir) and no genotyping call error on the true homozygote (1-ger_m_) OR the SNP is not informative (nir) but there is a genotyping call error on a homozygote (ger_m_) resulting in wrong homozygous SNP call (1 –*p*).

|  |  | SNP call state |  |  |  |  |
| --- | --- | --- | --- | --- | --- | --- |
|  |  | 0 (B73 homozygous) | 1 (heterozygous) | 2 (donor homozygous) | 3 (missing data) | Sum |
| True introgression state | 0 (B73 homoz) | $(1 - ger_m) * (1 - mr)$ | $ger_m * p * (1 - mr)$ | $ger_m * (1 - p) * (1 - mr)$ | mr | 1 |
| | 1 (het) | $[((1 - nir) * 0.5 * ger_t) + nir * (1 - ger_m)] * (1 - mr)$ | $[((1 - nir) * (1 - ger_t)) + (nir * ger_m * p)] * (1 - mr)$ | $[((1 - nir) * 0.5 * ger_t) + (nir * ger_m * (1 - p))] * (1 - mr)$ | mr | 1 |
| | 2 (donor homoz) | $[((1 - nir) * ger_m * (1 - p)) + (nir * (1 - ger_m))] * (1 - mr)$ | $ger_m * p * (1 - mr)$ | $[((1 - nir) * (1 - ger_m)) + (nir * ger_m * (1 - p))] * (1 - mr)$ | mr | 1 |

**Table S3.** Effects associated with GLS, NLB, or SLB at markers retained in final multiple QTL models assuming common introgression effects. GLS and SLB were scored on 1 – 9 scales (1 = more resistant), whereas NLB was scored on a percentage disease severity scale. Markers are denoted by an “S” followed by the chromosome number followed by an underscore, followed by the physical position on the chromosome in AGPv4 coordinates. Positive effect scores indicate increased susceptibility relative to the B73 allele and negative scores indicate increased resistance.

| Trait | Marker | Estimate | Std. Error | p-value |
| --- | --- | --- | --- | --- |
| GLS | (Intercept) | 3.93 | 0.01 | 0.0000 |
| GLS | S1_299212099 | 0.09 | 0.03 | 0.0045 |
| GLS | S9_113101198 | -0.13 | 0.03 | 0.0002 |
| GLS | S8_67560496 | -0.11 | 0.03 | 0.0002 |
| NLB | (Intercept) | 18.35 | 0.09 | 0.0000 |
| NLB | S1_185920012 | -1.95 | 0.26 | 0.0000 |
| NLB | S8_106963044 | -1.66 | 0.26 | 0.0000 |
| NLB | S5_15732819 | 0.88 | 0.21 | 0.0000 |
| NLB | S8_157816986 | -1.01 | 0.29 | 0.0006 |
| NLB | S1_84160146 | -0.88 | 0.30 | 0.0031 |
| SLB | (Intercept) | 4.48 | 0.02 | 0.0000 |
| SLB | S3_14748653 | -0.26 | 0.05 | 0.0000 |
| SLB | S8_67560496 | -0.26 | 0.04 | 0.0000 |
| SLB | S1_204753686 | -0.18 | 0.04 | 0.0000 |
| SLB | S3_223075551 | 0.22 | 0.04 | 0.0000 |
| SLB | S9_27020800 | -0.24 | 0.05 | 0.0000 |
| SLB | S1_92896641 | -0.19 | 0.05 | 0.0006 |

**Table S4.** Effects associated with GLS, NLB, or SLB at markers retained in final multiple QTL models with varying donor allele effects. GLS and SLB were scored on 1 – 9 scales (1 = more resistant), whereas NLB was scored on a percentage disease severity scale. Markers are denoted by and “S” followed by the chromosome number followed by an underscore, followed by the physical position on the chromosome in AGPv4 coordinates. Positive effect scores indicate increased susceptibility relative to the B73 allele and negative scores indicate increased resistance.

| Trait | Marker | Donor | Estimate | Std. Error | p-value |
| --- | --- | --- | --- | --- | --- |
| **GLS** | **(Intercept)** |  | **3.92** | **0.01** | **0.0000** |
| GLS | S3_10866532 | CML103 | -0.30 | 0.11 | 0.0080 |
| GLS | S3_10866532 | CML228 | 0.00 | 0.09 | 0.9732 |
| GLS | S3_10866532 | CML247 | 0.68 | 0.13 | 1.0e-07 |
| GLS | S3_10866532 | CML322 | 0.40 | 0.18 | 0.0252 |
| GLS | S3_10866532 | CML333 | 0.07 | 0.06 | 0.2611 |
| GLS | S3_10866532 | CML69 | -0.21 | 0.13 | 0.0938 |
| GLS | S3_10866532 | Ki3 | -0.06 | 0.13 | 0.6386 |
| GLS | S3_10866532 | M162W | 0.06 | 0.13 | 0.6473 |
| GLS | S3_10866532 | Mo17 | 0.03 | 0.09 | 0.7757 |
| GLS | S3_10866532 | Mo18W | -0.06 | 0.13 | 0.6551 |
| GLS | S3_10866532 | NC358 | 0.29 | 0.17 | 0.0880 |
| GLS | S3_10866532 | Oh43 | 0.07 | 0.11 | 0.5462 |
| GLS | S3_10866532 | Tx303 | -0.18 | 0.13 | 0.1465 |
| GLS | S3_10866532 | Tzi8 | 0.01 | 0.07 | 0.9300 |
| GLS | S1_241004561 | CML103 | -0.14 | 0.11 | 0.2147 |
| GLS | S1_241004561 | CML228 | -0.22 | 0.07 | 0.0012 |
| GLS | S1_241004561 | CML277 | -0.20 | 0.18 | 0.2639 |
| GLS | S1_241004561 | CML333 | -0.08 | 0.28 | 0.7815 |
| GLS | S1_241004561 | CML69 | -0.10 | 0.15 | 0.4948 |
| GLS | S1_241004561 | Ki11 | 0.39 | 0.13 | 0.0022 |
| GLS | S1_241004561 | Ki3 | 0.20 | 0.09 | 0.0240 |
| GLS | S1_241004561 | M162W | -0.16 | 0.13 | 0.1960 |
| GLS | S1_241004561 | Mo17 | -0.07 | 0.17 | 0.6820 |
| GLS | S1_241004561 | Oh43 | 0.03 | 0.09 | 0.7490 |
| GLS | S1_241004561 | Tx303 | 0.03 | 0.08 | 0.7371 |
| GLS | S1_241004561 | Tzi8 | -0.53 | 0.28 | 0.0611 |
| GLS | S9_15037724 | CML277 | -0.11 | 0.13 | 0.3666 |
| GLS | S9_15037724 | CML322 | -0.25 | 0.13 | 0.0426 |
| GLS | S9_15037724 | CML333 | -0.21 | 0.07 | 0.0032 |
| GLS | S9_15037724 | CML52 | -0.35 | 0.13 | 0.0048 |
| GLS | S9_15037724 | CML69 | 0.22 | 0.09 | 0.0155 |
| GLS | S9_15037724 | Mo17 | -0.31 | 0.15 | 0.0456 |
| GLS | S9_15037724 | NC350 | 0.14 | 0.07 | 0.0602 |
| GLS | S9_15037724 | NC358 | 0.28 | 0.17 | 0.0960 |
| GLS | S9_15037724 | Oh43 | -0.12 | 0.09 | 0.1609 |
| GLS | S9_15037724 | Tzi8 | -0.14 | 0.06 | 0.0237 |
| GLS | S8_73928265 | CML103 | -0.15 | 0.06 | 0.0091 |
| GLS | S8_73928265 | CML247 | -0.09 | 0.13 | 0.4890 |
| GLS | S8_73928265 | CML277 | -0.07 | 0.06 | 0.2273 |
| GLS | S8_73928265 | CML322 | -0.07 | 0.13 | 0.5532 |
| GLS | S8_73928265 | CML52 | 0.45 | 0.13 | 0.0003 |
| GLS | S8_73928265 | Mo17 | 0.07 | 0.15 | 0.6312 |
| GLS | S8_73928265 | Tx303 | 0.00 | 0.10 | 0.9699 |
| GLS | S8_73928265 | Tzi8 | -0.15 | 0.07 | 0.0342 |
| GLS | S3_171457420 | CML247 | 0.15 | 0.07 | 0.0361 |
| GLS | S3_171457420 | CML277 | 0.28 | 0.09 | 0.0017 |
| GLS | S3_171457420 | CML322 | 0.09 | 0.13 | 0.4708 |
| GLS | S3_171457420 | CML333 | 0.34 | 0.14 | 0.0159 |
| GLS | S3_171457420 | CML69 | 0.08 | 0.09 | 0.3465 |
| GLS | S3_171457420 | Ki11 | 0.06 | 0.07 | 0.4236 |
| GLS | S3_171457420 | M162W | 0.00 | 0.13 | 0.9986 |
| GLS | S3_171457420 | Mo17 | -0.06 | 0.10 | 0.5771 |
| GLS | S3_171457420 | Mo18W | 0.14 | 0.09 | 0.1269 |
| GLS | S3_171457420 | NC358 | -0.32 | 0.11 | 0.0043 |
| GLS | S3_171457420 | Oh43 | 0.10 | 0.14 | 0.5077 |
| GLS | S3_171457420 | Tx303 | -0.03 | 0.13 | 0.8092 |
| GLS | S3_171457420 | Tzi8 | -0.01 | 0.06 | 0.8952 |
| GLS | S1_235322076 | CML103 | -0.04 | 0.09 | 0.6569 |
| GLS | S1_235322076 | CML277 | -0.02 | 0.13 | 0.8857 |
| GLS | S1_235322076 | CML333 | -0.13 | 0.25 | 0.6159 |
| GLS | S1_235322076 | CML69 | 0.18 | 0.13 | 0.1538 |
| GLS | S1_235322076 | Mo17 | -0.03 | 0.12 | 0.7945 |
| GLS | S1_235322076 | Tzi8 | 1.07 | 0.35 | 0.0027 |
| GLS | S5_209976785 | CML228 | 0.32 | 0.13 | 0.0109 |
| GLS | S5_209976785 | CML69 | -0.11 | 0.13 | 0.3768 |
| GLS | S5_209976785 | Mo18W | -0.08 | 0.13 | 0.5321 |
| GLS | S5_209976785 | NC350 | 0.06 | 0.13 | 0.6240 |
| GLS | S5_209976785 | NC358 | 0.10 | 0.09 | 0.2424 |
| GLS | S5_209976785 | Oh43 | -0.03 | 0.13 | 0.7826 |
| GLS | S5_209976785 | Tzi8 | -0.14 | 0.07 | 0.0510 |
| **NLB** | **(Intercept)** |  | **18.36** | **0.08** | **0.0000** |
| NLB | S1_168645419 | CML103 | 1.63 | 2.22 | 0.4618 |
| NLB | S1_168645419 | CML277 | 0.24 | 1.42 | 0.8678 |
| NLB | S1_168645419 | CML322 | -1.95 | 1.42 | 0.1686 |
| NLB | S1_168645419 | CML333 | 1.71 | 1.00 | 0.0893 |
| NLB | S1_168645419 | CML69 | 3.95 | 1.00 | 9.01e-05 |
| NLB | S1_168645419 | Ki11 | -2.69 | 1.55 | 0.0833 |
| NLB | S1_168645419 | M162W | -3.32 | 1.00 | 0.0010 |
| NLB | S1_168645419 | Oh43 | -0.86 | 1.83 | 0.6367 |
| NLB | S1_168645419 | Tzi8 | -2.76 | 2.14 | 0.1984 |
| NLB | S8_106963044 | CML103 | 0.75 | 2.24 | 0.7393 |
| NLB | S8_106963044 | CML277 | 0.11 | 1.12 | 0.9222 |
| NLB | S8_106963044 | CML322 | -0.74 | 1.00 | 0.4617 |
| NLB | S8_106963044 | CML52 | -0.96 | 0.71 | 0.1746 |
| NLB | S8_106963044 | M162W | -5.08 | 1.00 | 5.4e-07 |
| NLB | S8_106963044 | Mo17 | -0.83 | 0.79 | 0.2971 |
| NLB | S8_106963044 | Oh43 | 0.15 | 1.00 | 0.8817 |
| NLB | S8_106963044 | Tx303 | -1.60 | 1.00 | 0.1116 |
| NLB | S8_106963044 | Tzi8 | -4.73 | 0.77 | 1.3e-09 |
| NLB | S1_219232354 | CML103 | -0.90 | 0.71 | 0.2058 |
| NLB | S1_219232354 | CML228 | 0.94 | 2.13 | 0.6572 |
| NLB | S1_219232354 | CML277 | -1.25 | 1.42 | 0.3765 |
| NLB | S1_219232354 | CML52 | -1.42 | 0.58 | 0.0146 |
| NLB | S1_219232354 | CML69 | 1.29 | 2.46 | 0.5997 |
| NLB | S1_219232354 | Ki11 | -0.47 | 1.00 | 0.6378 |
| NLB | S1_219232354 | Mo17 | 0.87 | 0.79 | 0.2748 |
| NLB | S1_219232354 | Tx303 | -0.34 | 1.74 | 0.8439 |
| NLB | S1_219232354 | Tzi8 | 4.68 | 1.52 | 0.0021 |
| NLB | S1_82555693 | CML103 | -0.04 | 1.27 | 0.9734 |
| NLB | S1_82555693 | CML228 | 4.16 | 1.00 | 3.9e-05 |
| NLB | S1_82555693 | CML322 | -0.83 | 1.00 | 0.4066 |
| NLB | S1_82555693 | NC350 | -1.08 | 0.58 | 0.0623 |
| NLB | S1_82555693 | NC358 | -0.89 | 0.71 | 0.2095 |
| NLB | S1_82555693 | Oh43 | -3.48 | 1.74 | 0.0457 |
| NLB | S1_82555693 | Tx303 | -0.23 | 0.58 | 0.6933 |
| NLB | S8_103049205 | CML103 | -2.92 | 2.07 | 0.1578 |
| NLB | S8_103049205 | CML228 | -1.05 | 0.50 | 0.0377 |
| NLB | S8_103049205 | CML277 | -1.73 | 0.50 | 0.0006 |
| NLB | S8_103049205 | Tzi8 | 1.66 | 0.65 | 0.0106 |
| NLB | S1_188722694 | CML103 | -4.97 | 2.01 | 0.0135 |
| NLB | S1_188722694 | CML277 | -1.72 | 1.00 | 0.0875 |
| NLB | S1_188722694 | Ki11 | -2.42 | 1.00 | 0.0162 |
| NLB | S1_188722694 | Mo17 | -1.50 | 0.71 | 0.0346 |
| NLB | S1_188722694 | Oh43 | -1.51 | 1.00 | 0.1321 |
| NLB | S1_188722694 | Tzi8 | -1.66 | 0.76 | 0.0287 |
| NLB | S1_157231888 | CML69 | -2.18 | 1.42 | 0.1247 |
| NLB | S1_157231888 | Oh43 | 4.25 | 1.16 | 0.0003 |
| NLB | S1_177789781 | CML69 | -3.30 | 2.24 | 0.1417 |
| NLB | S1_177789781 | Ki11 | 2.48 | 1.00 | 0.0138 |
| NLB | S1_177789781 | Oh43 | -1.69 | 1.00 | 0.0931 |
| NLB | S5_34841324 | CML103 | 0.24 | 0.38 | 0.5255 |
| NLB | S5_34841324 | CML322 | 2.09 | 0.71 | 0.0033 |
| NLB | S5_34841324 | CML333 | 0.10 | 0.58 | 0.8585 |
| NLB | S5_34841324 | Ki11 | -1.52 | 0.66 | 0.0222 |
| NLB | S5_34841324 | Ki3 | 0.44 | 1.00 | 0.6621 |
| NLB | S5_34841324 | M162W | -0.25 | 1.00 | 0.8012 |
| NLB | S5_34841324 | Mo18W | 0.68 | 1.00 | 0.4967 |
| NLB | S5_34841324 | NC358 | -0.01 | 0.50 | 0.9835 |
| NLB | S5_34841324 | Oh43 | 2.16 | 1.16 | 0.0626 |
| NLB | S5_34841324 | Tx303 | 0.00 | 1.00 | 0.9982 |
| NLB | S5_34841324 | Tzi8 | -1.62 | 1.00 | 0.1070 |
| NLB | S8_155129399 | CML322 | -0.44 | 0.58 | 0.4513 |
| NLB | S8_155129399 | CML333 | -0.71 | 0.71 | 0.3203 |
| NLB | S8_155129399 | CML52 | -0.51 | 1.23 | 0.6784 |
| NLB | S8_155129399 | CML69 | -1.55 | 0.71 | 0.0291 |
| NLB | S8_155129399 | NC350 | -0.43 | 1.00 | 0.6710 |
| NLB | S8_155129399 | Tx303 | -2.01 | 1.42 | 0.1573 |
| NLB | S8_155129399 | Tzi8 | 1.73 | 0.61 | 0.0045 |
| NLB | S1_36755096 | CML277 | -1.11 | 0.50 | 0.0270 |
| NLB | S1_36755096 | Ki11 | -1.67 | 0.72 | 0.0202 |
| NLB | S1_36755096 | Oh43 | 0.87 | 1.00 | 0.3885 |
| NLB | S1_36755096 | Tx303 | -0.43 | 0.71 | 0.5453 |
| NLB | S1_36755096 | Tzi8 | -0.50 | 0.71 | 0.4828 |
| NLB | S10_4470291 | CML228 | 0.30 | 0.71 | 0.6753 |
| NLB | S10_4470291 | CML52 | -0.63 | 1.00 | 0.5278 |
| NLB | S10_4470291 | CML69 | 0.32 | 0.71 | 0.6522 |
| NLB | S10_4470291 | Ki11 | -0.67 | 0.65 | 0.3035 |
| NLB | S10_4470291 | Mo17 | 1.08 | 0.58 | 0.0626 |
| NLB | S10_4470291 | Mo18W | 0.83 | 0.50 | 0.0998 |
| NLB | S10_4470291 | NC350 | 2.16 | 0.71 | 0.0024 |
| NLB | S10_4470291 | Tzi8 | -0.13 | 1.00 | 0.8932 |
| NLB | S1_214380338 | CML103 | 1.39 | 1.27 | 0.2728 |
| NLB | S1_214380338 | CML228 | -1.44 | 2.01 | 0.4735 |
| NLB | S1_214380338 | NC350 | 0.64 | 1.00 | 0.5256 |
| NLB | S1_214380338 | Tx303 | 2.55 | 1.00 | 0.0112 |
| NLB | S1_214380338 | Tzi8 | 0.68 | 0.54 | 0.2027 |
| NLB | S1_213709429 | Mo17 | -1.08 | 0.79 | 0.1750 |
| **SLB** | **(Intercept)** |  | **4.48** | **0.01** | **0.0000** |
| SLB | S3_14337777 | CML103 | -0.29 | 0.34 | 0.3909 |
| SLB | S3_14337777 | CML228 | -0.69 | 0.26 | 0.0072 |
| SLB | S3_14337777 | CML247 | 0.44 | 0.17 | 0.0092 |
| SLB | S3_14337777 | CML322 | 0.68 | 0.24 | 0.0043 |
| SLB | S3_14337777 | CML333 | -0.55 | 0.21 | 0.0081 |
| SLB | S3_14337777 | CML69 | -0.56 | 0.29 | 0.0531 |
| SLB | S3_14337777 | Ki3 | -0.79 | 0.17 | 3.6e-06 |
| SLB | S3_14337777 | M162W | -0.04 | 0.17 | 0.8007 |
| SLB | S3_14337777 | Mo17 | 0.23 | 0.24 | 0.3257 |
| SLB | S3_14337777 | NC358 | 0.15 | 0.24 | 0.5264 |
| SLB | S3_14337777 | Oh43 | 0.06 | 0.17 | 0.7371 |
| SLB | S3_14337777 | Tzi8 | -0.85 | 0.24 | 0.0004 |
| SLB | S1_217472496 | CML103 | -0.19 | 0.27 | 0.4782 |
| SLB | S1_217472496 | CML228 | -0.20 | 0.11 | 0.0784 |
| SLB | S1_217472496 | CML277 | -0.19 | 0.17 | 0.2474 |
| SLB | S1_217472496 | CML52 | 0.00 | 0.10 | 0.9882 |
| SLB | S1_217472496 | CML69 | -0.70 | 0.24 | 0.0034 |
| SLB | S1_217472496 | Ki11 | 0.30 | 0.17 | 0.0761 |
| SLB | S1_217472496 | Mo17 | -0.09 | 0.17 | 0.5881 |
| SLB | S1_217472496 | Tzi8 | -1.03 | 0.18 | 3.7e-08 |
| SLB | S5_176932465 | CML247 | 0.11 | 0.17 | 0.5019 |
| SLB | S5_176932465 | CML277 | -0.93 | 0.17 | 4.8e-08 |
| SLB | S5_176932465 | CML333 | 0.14 | 0.17 | 0.4029 |
| SLB | S5_176932465 | CML52 | -0.57 | 0.17 | 0.0007 |
| SLB | S5_176932465 | Ki11 | 0.12 | 0.11 | 0.2659 |
| SLB | S5_176932465 | NC358 | 0.27 | 0.10 | 0.0058 |
| SLB | S5_176932465 | Oh43 | 0.13 | 0.11 | 0.2164 |
| SLB | S5_176932465 | Tzi8 | 0.01 | 0.21 | 0.9508 |
| SLB | S8_90126501 | CML103 | -0.22 | 0.10 | 0.0261 |
| SLB | S8_90126501 | CML228 | -0.42 | 0.10 | 2.1e-05 |
| SLB | S8_90126501 | CML247 | -0.23 | 0.17 | 0.1724 |
| SLB | S8_90126501 | CML277 | -0.20 | 0.08 | 0.0080 |
| SLB | S8_90126501 | CML322 | -0.14 | 0.17 | 0.3897 |
| SLB | S8_90126501 | CML52 | 0.08 | 0.10 | 0.4270 |
| SLB | S8_90126501 | M162W | 0.18 | 0.17 | 0.2780 |
| SLB | S8_90126501 | Mo17 | -0.90 | 0.34 | 0.0073 |
| SLB | S8_90126501 | Oh43 | 0.28 | 0.17 | 0.0962 |
| SLB | S8_90126501 | Tx303 | -0.70 | 0.25 | 0.0047 |
| SLB | S8_90126501 | Tzi8 | -0.08 | 0.09 | 0.3555 |
| SLB | S10_140616277 | CML103 | -0.02 | 0.24 | 0.9376 |
| SLB | S10_140616277 | CML277 | 0.43 | 0.12 | 0.0003 |
| SLB | S10_140616277 | CML69 | 0.10 | 0.17 | 0.5621 |
| SLB | S10_140616277 | Ki11 | -0.06 | 0.12 | 0.6312 |
| SLB | S10_140616277 | M162W | -0.33 | 0.17 | 0.0470 |
| SLB | S10_140616277 | Mo17 | -0.12 | 0.17 | 0.4603 |
| SLB | S10_140616277 | Mo18W | -0.17 | 0.08 | 0.0414 |
| SLB | S10_140616277 | NC350 | -0.38 | 0.12 | 0.0013 |
| SLB | S10_140616277 | Oh43 | 0.37 | 0.17 | 0.0275 |
| SLB | S10_140616277 | Tzi8 | -0.23 | 0.12 | 0.0521 |
| SLB | S3_9660537 | CML103 | -0.26 | 0.17 | 0.1193 |
| SLB | S3_9660537 | CML228 | -0.37 | 0.26 | 0.1486 |
| SLB | S3_9660537 | CML333 | 0.60 | 0.19 | 0.0021 |
| SLB | S3_9660537 | CML52 | 0.14 | 0.12 | 0.2322 |
| SLB | S3_9660537 | Mo17 | -2.15 | 0.57 | 0.0002 |
| SLB | S3_9660537 | Mo18W | 0.04 | 0.17 | 0.8135 |
| SLB | S3_9660537 | Oh43 | -0.88 | 0.29 | 0.0026 |
| SLB | S3_9660537 | Tx303 | 0.13 | 0.17 | 0.4248 |
| SLB | S3_9660537 | Tzi8 | 0.82 | 0.27 | 0.0021 |
| SLB | S9_18679092 | CML103 | -0.33 | 0.12 | 0.0051 |
| SLB | S9_18679092 | CML228 | 0.23 | 0.34 | 0.4842 |
| SLB | S9_18679092 | CML247 | 0.11 | 0.17 | 0.5043 |
| SLB | S9_18679092 | CML277 | -0.28 | 0.17 | 0.1010 |
| SLB | S9_18679092 | CML322 | -0.58 | 0.17 | 0.0006 |
| SLB | S9_18679092 | CML333 | -0.11 | 0.12 | 0.3754 |
| SLB | S9_18679092 | CML69 | 0.11 | 0.12 | 0.3522 |
| SLB | S9_18679092 | Mo17 | 0.13 | 0.12 | 0.2656 |
| SLB | S9_18679092 | NC350 | 0.24 | 0.17 | 0.1516 |
| SLB | S9_18679092 | NC358 | -0.18 | 0.17 | 0.2969 |
| SLB | S9_18679092 | Oh43 | -0.70 | 0.17 | 3.4e-05 |
| SLB | S9_18679092 | Tzi8 | 0.11 | 0.08 | 0.1940 |
| SLB | S1_158112237 | CML103 | -0.37 | 0.12 | 0.0017 |
| SLB | S1_158112237 | CML277 | -0.11 | 0.17 | 0.5041 |
| SLB | S1_158112237 | CML322 | -0.03 | 0.17 | 0.8538 |
| SLB | S1_158112237 | CML333 | 0.14 | 0.25 | 0.5729 |
| SLB | S1_158112237 | CML69 | 0.17 | 0.17 | 0.3105 |
| SLB | S1_158112237 | Ki11 | -0.35 | 0.08 | 1.4e-05 |
| SLB | S1_158112237 | M162W | -0.21 | 0.17 | 0.2078 |
| SLB | S1_158112237 | Oh43 | 0.12 | 0.09 | 0.1727 |
| SLB | S3_19067827 | CML69 | 0.05 | 0.29 | 0.8762 |
| SLB | S3_19067827 | M162W | -0.29 | 0.17 | 0.0814 |
| SLB | S3_19067827 | Mo17 | 1.15 | 0.43 | 0.0072 |
| SLB | S3_19067827 | Tx303 | -1.68 | 0.43 | 0.0001 |
| SLB | S3_19067827 | Tzi8 | -0.30 | 0.24 | 0.2125 |
| SLB | S3_155684866 | CML103 | -0.28 | 0.18 | 0.1342 |
| SLB | S3_155684866 | CML247 | 0.27 | 0.10 | 0.0060 |
| SLB | S3_155684866 | CML277 | 0.11 | 0.12 | 0.3660 |
| SLB | S3_155684866 | CML322 | 0.10 | 0.17 | 0.5698 |
| SLB | S3_155684866 | CML69 | 0.38 | 0.17 | 0.0249 |
| SLB | S3_155684866 | Ki11 | 0.20 | 0.34 | 0.5460 |
| SLB | S3_155684866 | Ki3 | 0.10 | 0.17 | 0.5479 |
| SLB | S3_155684866 | Mo17 | -1.26 | 0.62 | 0.0408 |
| SLB | S3_155684866 | Oh43 | -0.03 | 0.17 | 0.8419 |
| SLB | S3_155684866 | Tx303 | 3.64 | 0.83 | 1.5e-05 |
| SLB | S3_155684866 | Tzi8 | 0.45 | 0.17 | 0.0073 |
| SLB | S3_5707369 | CML322 | 0.22 | 0.10 | 0.0222 |
| SLB | S3_5707369 | Mo17 | 0.45 | 0.17 | 0.0074 |
| SLB | S3_5707369 | NC350 | -0.13 | 0.17 | 0.4290 |
| SLB | S3_5707369 | NC358 | -0.22 | 0.17 | 0.1895 |
| SLB | S3_5707369 | Tzi8 | 0.06 | 0.17 | 0.7058 |
| SLB | S3_30519885 | CML228 | 0.08 | 0.17 | 0.6260 |
| SLB | S3_30519885 | CML69 | -0.49 | 0.17 | 0.0038 |
| SLB | S3_30519885 | Mo17 | 0.55 | 0.39 | 0.1639 |
| SLB | S3_30519885 | NC350 | -0.45 | 0.17 | 0.0079 |
| SLB | S3_30519885 | NC358 | -0.04 | 0.17 | 0.7998 |
| SLB | S3_30519885 | Tx303 | -2.63 | 0.69 | 0.0002 |
| SLB | S1_241028181 | CML103 | 0.04 | 0.17 | 0.8008 |
| SLB | S1_241028181 | CML333 | -0.36 | 0.17 | 0.0313 |
| SLB | S1_241028181 | CML69 | 0.29 | 0.17 | 0.0853 |
| SLB | S1_241028181 | Ki3 | 0.27 | 0.17 | 0.1065 |
| SLB | S1_241028181 | M162W | -0.13 | 0.17 | 0.4526 |
| SLB | S1_241028181 | Mo17 | 0.89 | 0.39 | 0.0237 |
| SLB | S1_241028181 | Oh43 | 0.12 | 0.12 | 0.3035 |
| SLB | S1_241028181 | Tx303 | 0.52 | 0.14 | 0.0002 |
| SLB | S6_153427535 | CML103 | -0.13 | 0.08 | 0.0918 |
| SLB | S6_153427535 | CML333 | 0.09 | 0.08 | 0.2374 |
| SLB | S6_153427535 | CML52 | 0.14 | 0.17 | 0.4167 |
| SLB | S6_153427535 | CML69 | 0.12 | 0.08 | 0.1047 |
| SLB | S6_153427535 | Mo17 | 0.39 | 0.41 | 0.3432 |
| SLB | S6_153427535 | Mo18W | 0.39 | 0.17 | 0.0218 |
| SLB | S6_153427535 | NC350 | -0.23 | 0.17 | 0.1723 |
| SLB | S6_153427535 | NC358 | -0.07 | 0.17 | 0.6660 |
| SLB | S6_153427535 | Tzi8 | -0.11 | 0.07 | 0.1095 |
| SLB | S8_65142538 | CML103 | -0.18 | 0.15 | 0.2399 |
| SLB | S8_65142538 | Mo17 | 0.76 | 0.44 | 0.0889 |
| SLB | S8_65142538 | Mo18W | -0.20 | 0.12 | 0.0865 |
| SLB | S8_65142538 | Tx303 | -0.17 | 0.12 | 0.1497 |
| SLB | S3_9767602 | CML333 | -0.29 | 0.17 | 0.0873 |
| SLB | S3_119352940 | Ki11 | -0.13 | 0.58 | 0.8217 |
| SLB | S3_119352940 | Mo17 | -0.30 | 0.36 | 0.3974 |
| SLB | S3_119352940 | Mo18W | -0.84 | 0.34 | 0.0125 |
| SLB | S3_119352940 | Tx303 | -0.04 | 0.17 | 0.8064 |
| SLB | S1_212545654 | Tzi8 | -0.13 | 0.08 | 0.0976 |
| SLB | S8_166377016 | CML103 | 0.08 | 0.12 | 0.5040 |
| SLB | S8_166377016 | CML322 | 0.21 | 0.17 | 0.2127 |
| SLB | S8_166377016 | CML69 | 0.20 | 0.10 | 0.0393 |
| SLB | S8_166377016 | Ki11 | 0.00 | 0.13 | 0.9808 |
| SLB | S8_166377016 | NC350 | -0.02 | 0.15 | 0.8891 |
| SLB | S8_166377016 | Tzi8 | -0.18 | 0.08 | 0.0335 |
| SLB | S3_131596725 | Ki11 | 0.55 | 0.34 | 0.1043 |

## Supplemental Figures

**Figure S1.**
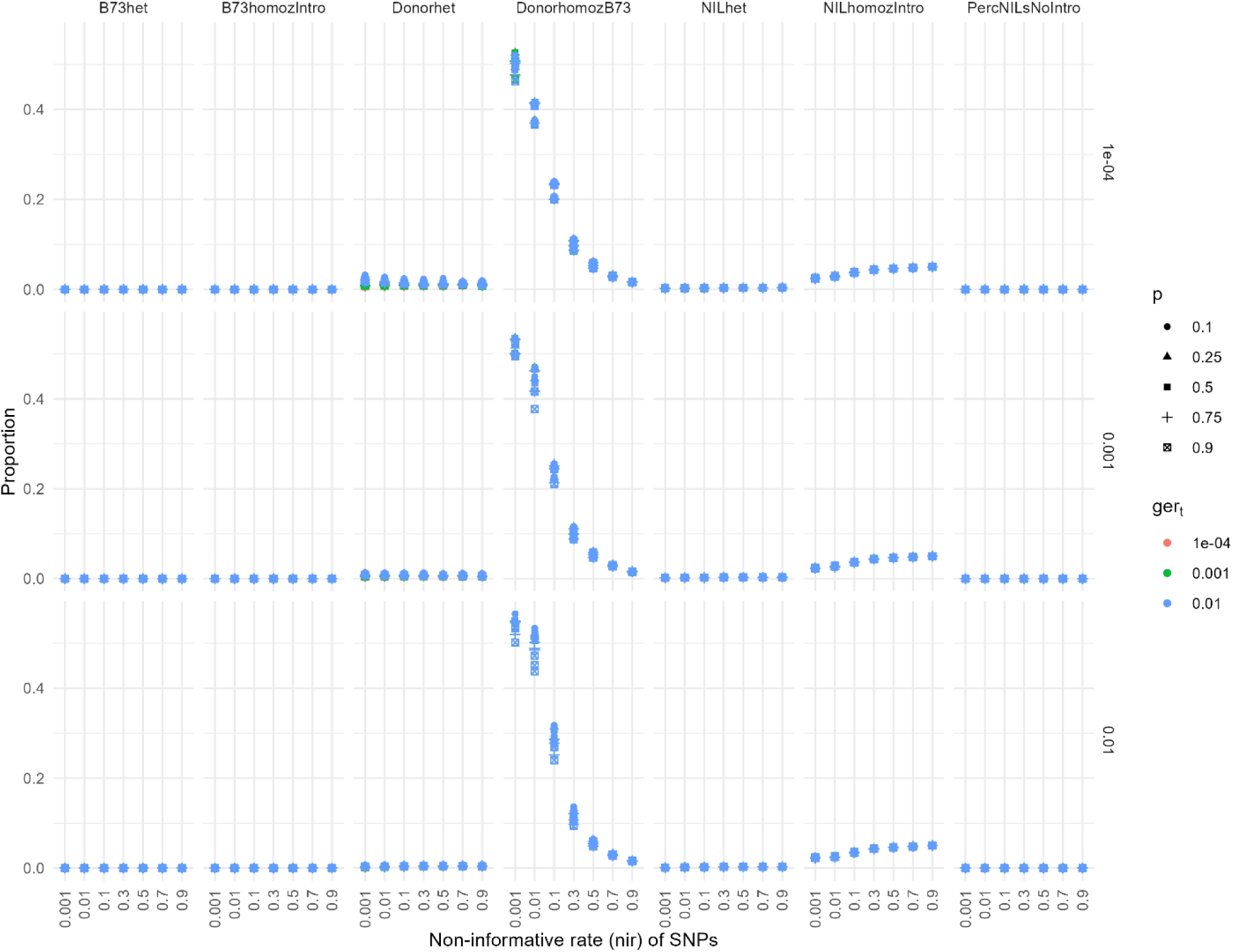
HMM grid search results of chip data. X axis is non-informative rate of SNPs, Y-axis is the proportion of introgression calls that are heterozygous (“B73het”) or homozygous for introgression haplotypes (“B73homozIntro”) in B73 samples; heterozygous (“Donorhet”) or homozygous for introgression haplotypes (“DonorhomozIntro”) in donor parent samples; heterozygous (“NILhet”) or homozygous for introgression haplotypes (“NILhomozIntro”) in nNIL samples; or the proportion of nNILs with no introgression calls (“PercNILSNoIntro”). Rows represent genotyping error rate on homozygotes (*ger_m_*), colors represent genotyping error rate on heterozygotes (*ger_t_*), and shapes represent probability that a homozygous genotyping error results in a heterozygous call (*p*). Three different values of adjacent marker pair recombination rates were tested, but their results were always nearly identical so they are not distinguished by shape or color in this figure.

**Figure S2.**
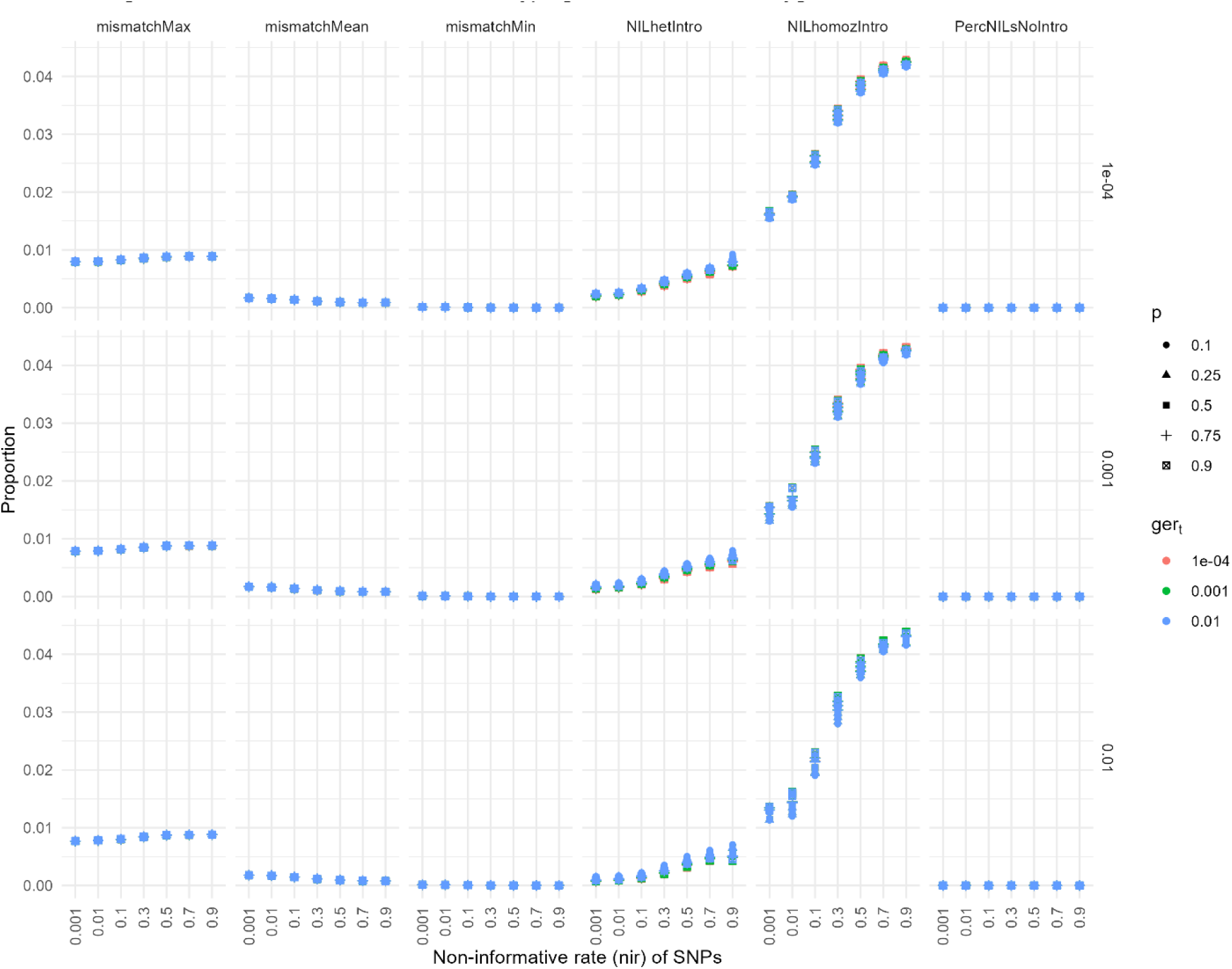
HMM grid search results of GBS data, comparing introgression calls to genotyping chip data on subset of lines genotyped with both methods. X axis is non-informative rate of SNPs, Y-axis is the maximum, mean, or minimum mismatch rate between the two methods across lines (“mismatchMax”, “mismatchMean”, “mismatchMin”), the proportion of heterozygous introgression calls on nNILs (“NILhetIntro”), the proportion of homozygous introgression calls on nNILs (“NILhomozIntro”), or the proportion of nNILs with no introgression calls (“PercNILSNoIntro”). Rows represent genotyping error rate on homozygotes (*ger_m_*), colors represent genotyping error rate on heterozygotes (*ger_t_*), and shapes represent probability that a homozygous genotyping error results in a heterozygous call (*p*). Three different values of adjacent marker pair recombination rates were tested, but their results were always nearly identical so they are not distinguished by shape or color in this figure.

**Figure S3.**
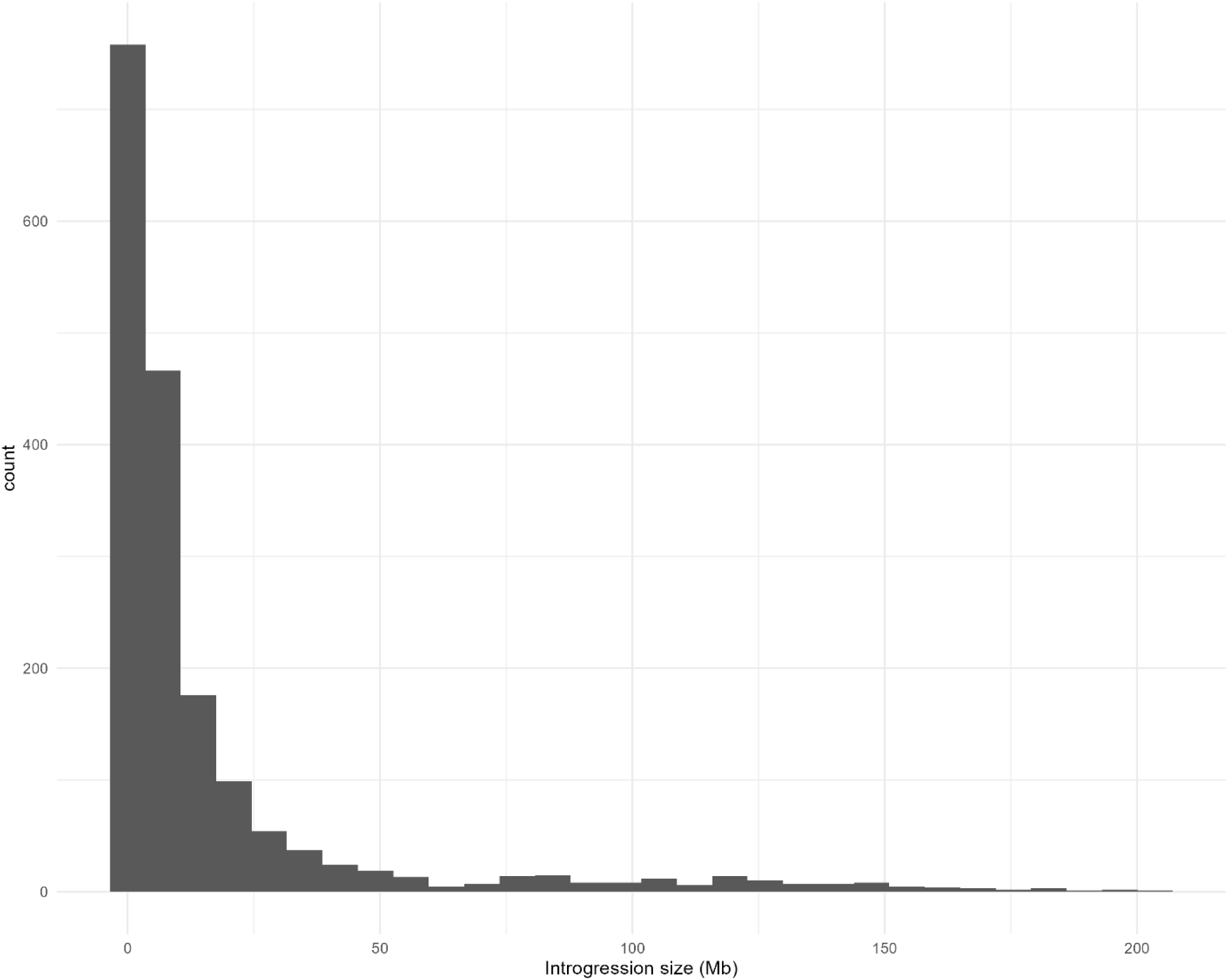
Histogram of individual introgression block sizes in Mbp among the 593 nNILs for which the accurate pedigree was confirmed.

**Figure S4.**
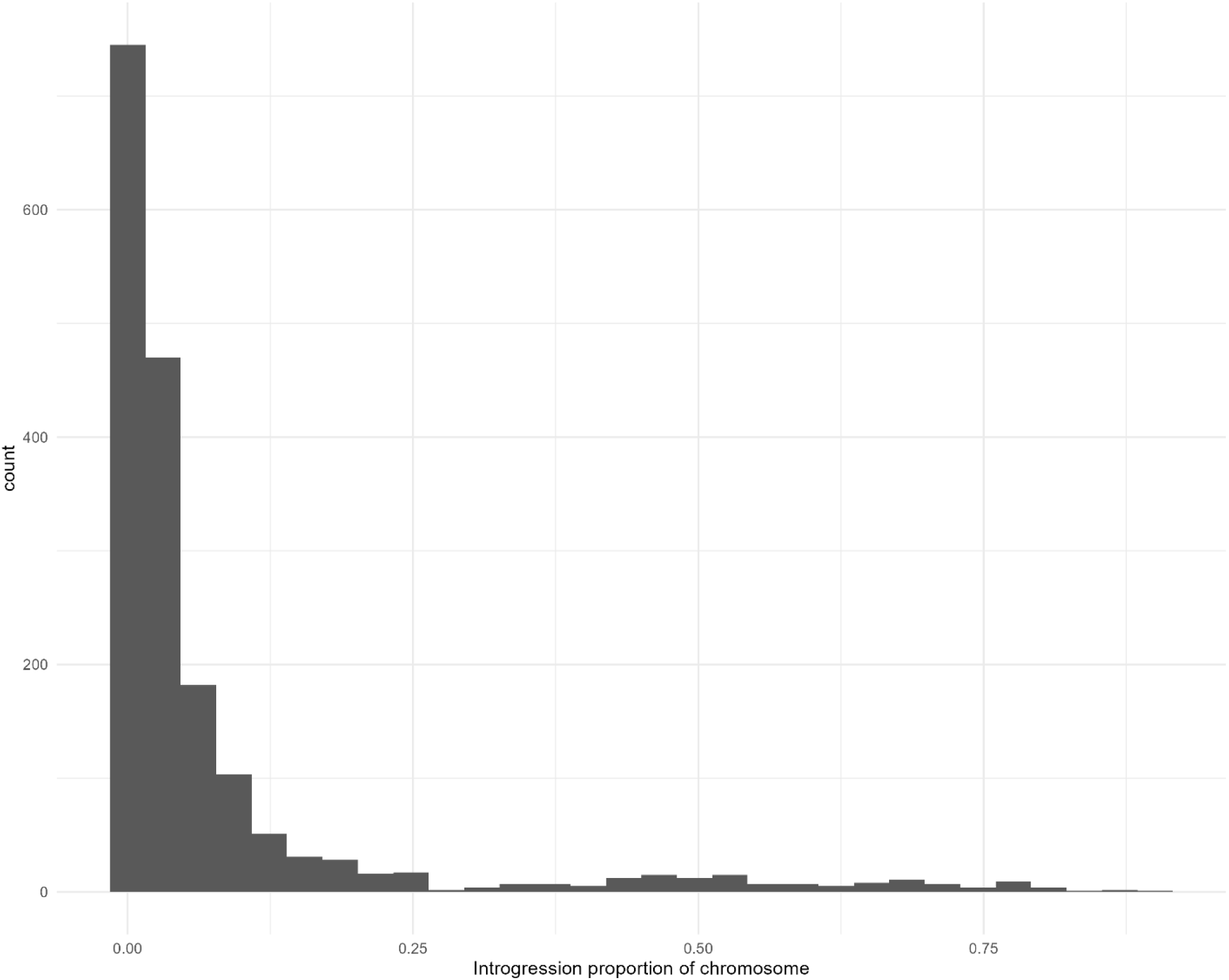
Histogram of individual introgression block sizes as proportions of their respective chromosomes.

**Figure S5.**
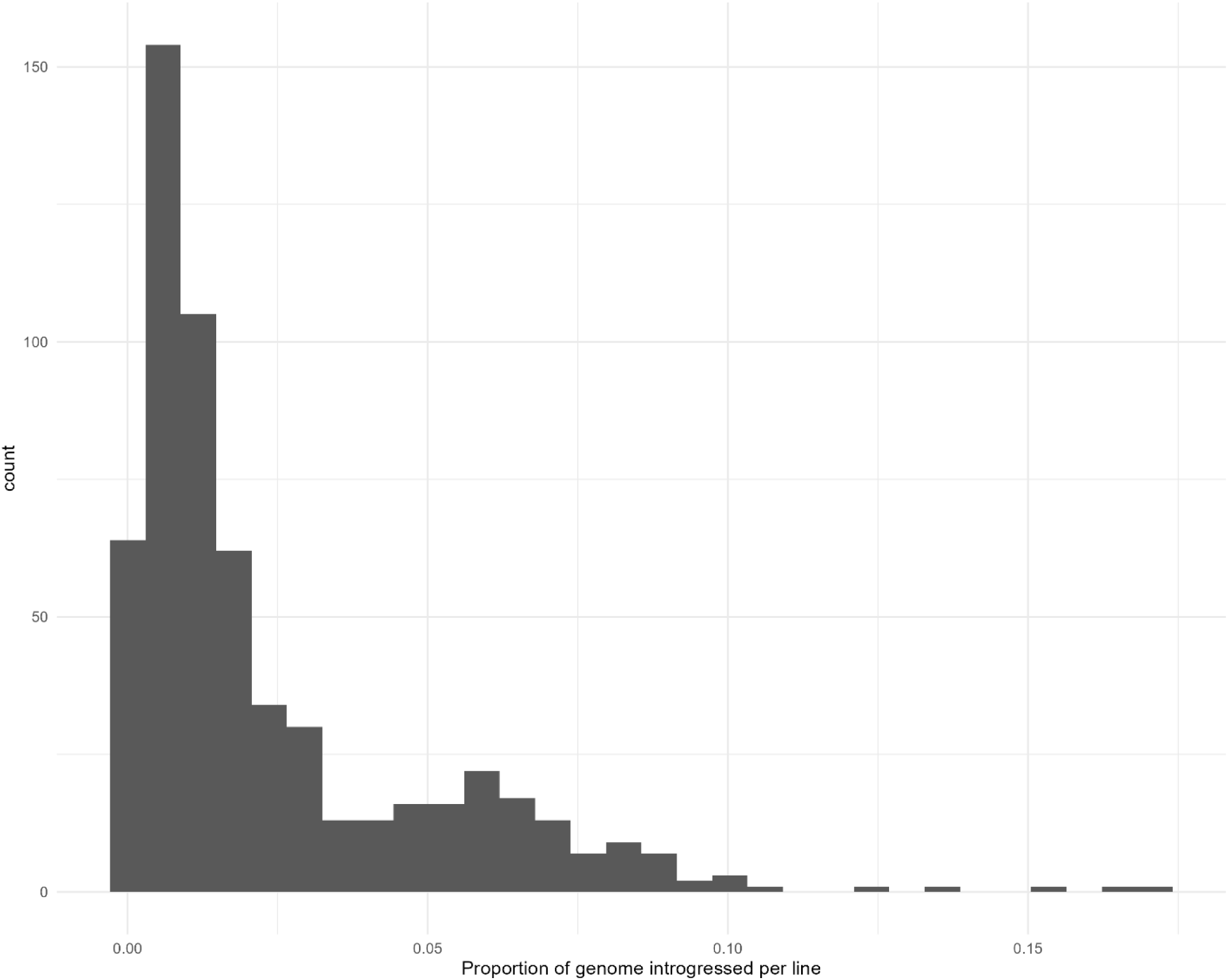
Histogram of proportion of genome introgressed in each line.

**Figure S6.**
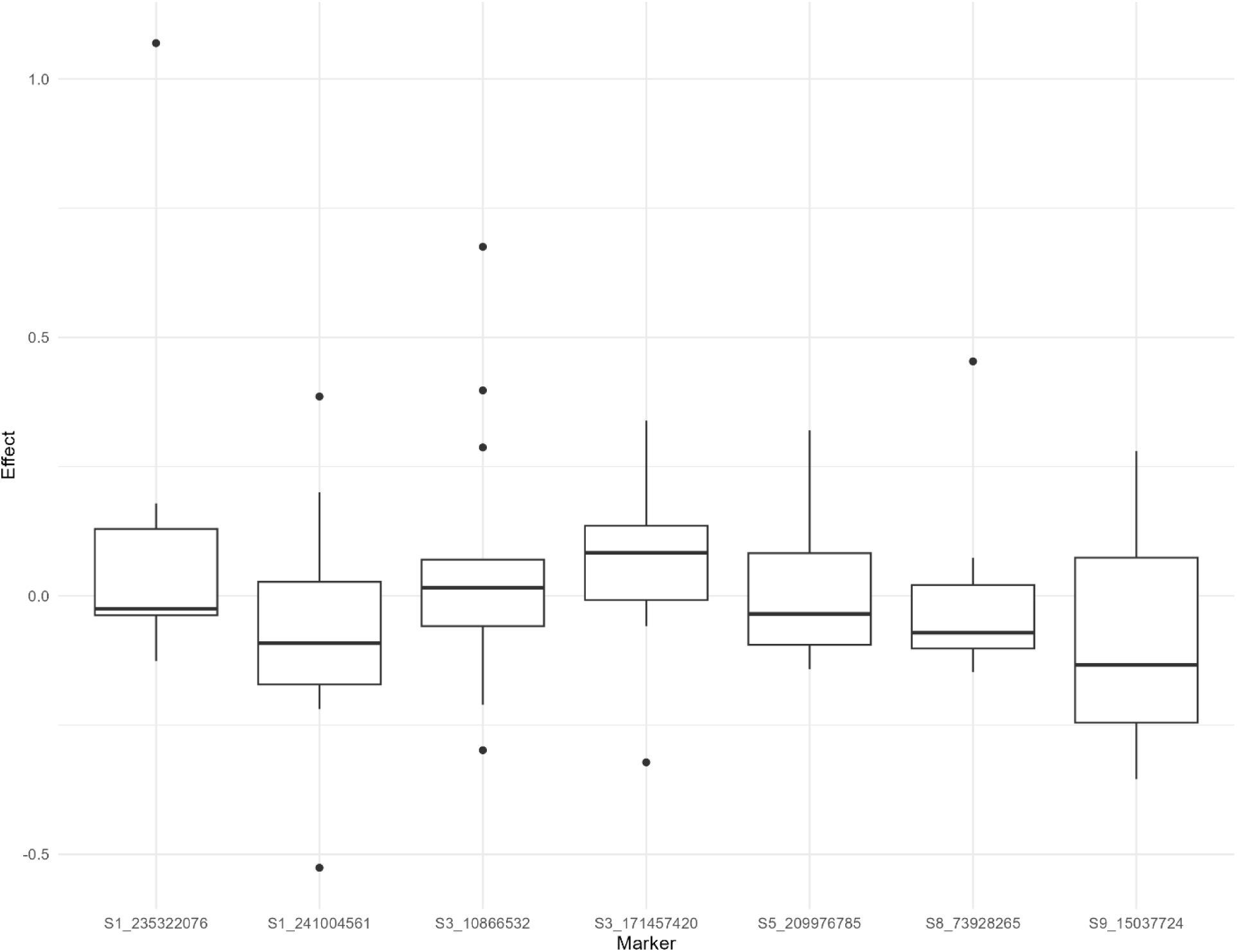
Boxplots of distribution of donor allele effects at QTL selected in the final multiple QTL model with variable donor allele effects for GLS. Negative effects correspond to increased resistance associated with introgressions.

**Figure S7.**
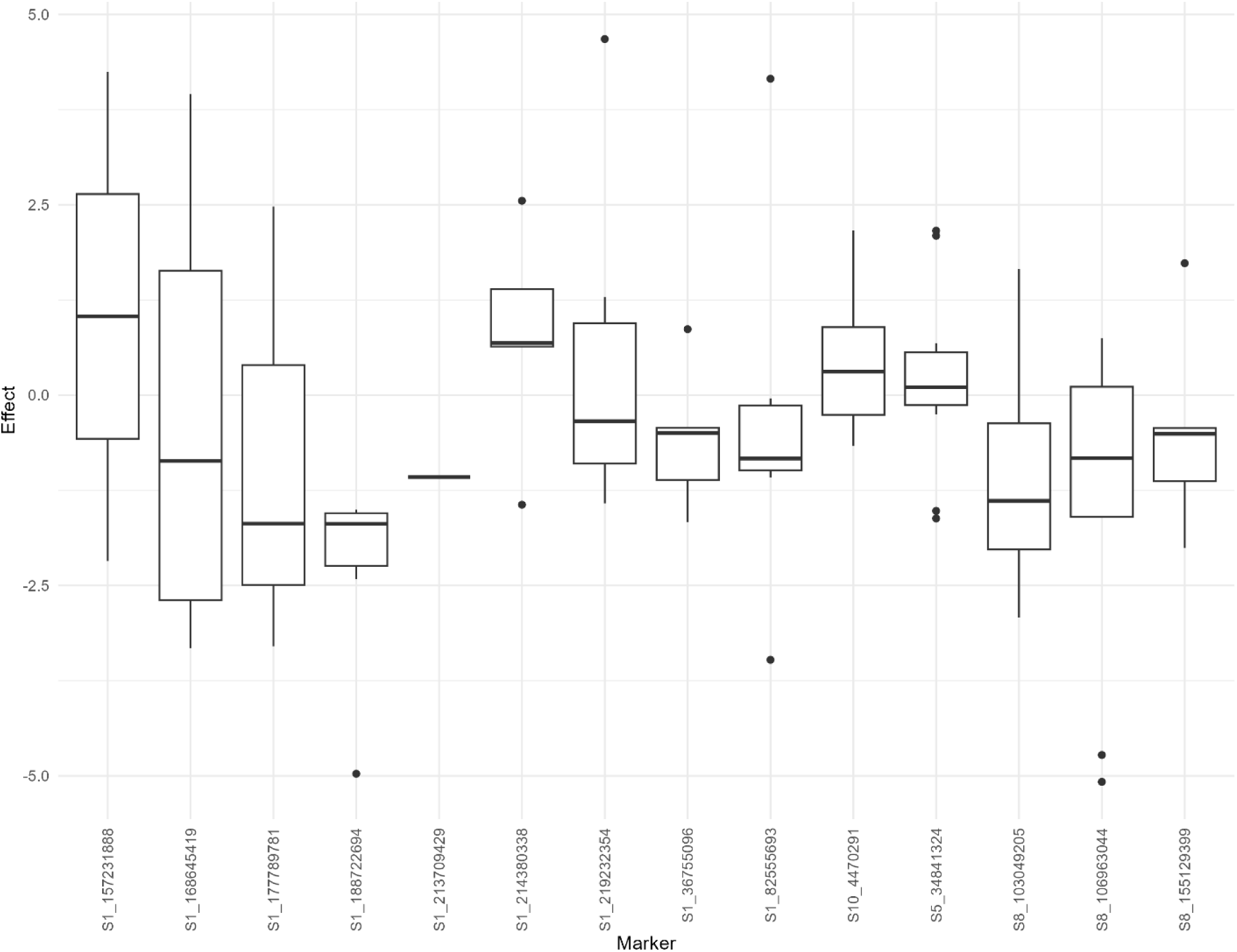
Boxplots of distribution of donor allele effects at QTL selected in the final multiple QTL model with variable donor allele effects for NLB. Negative effects correspond to increased resistance associated with introgressions.

**Figure S8.**
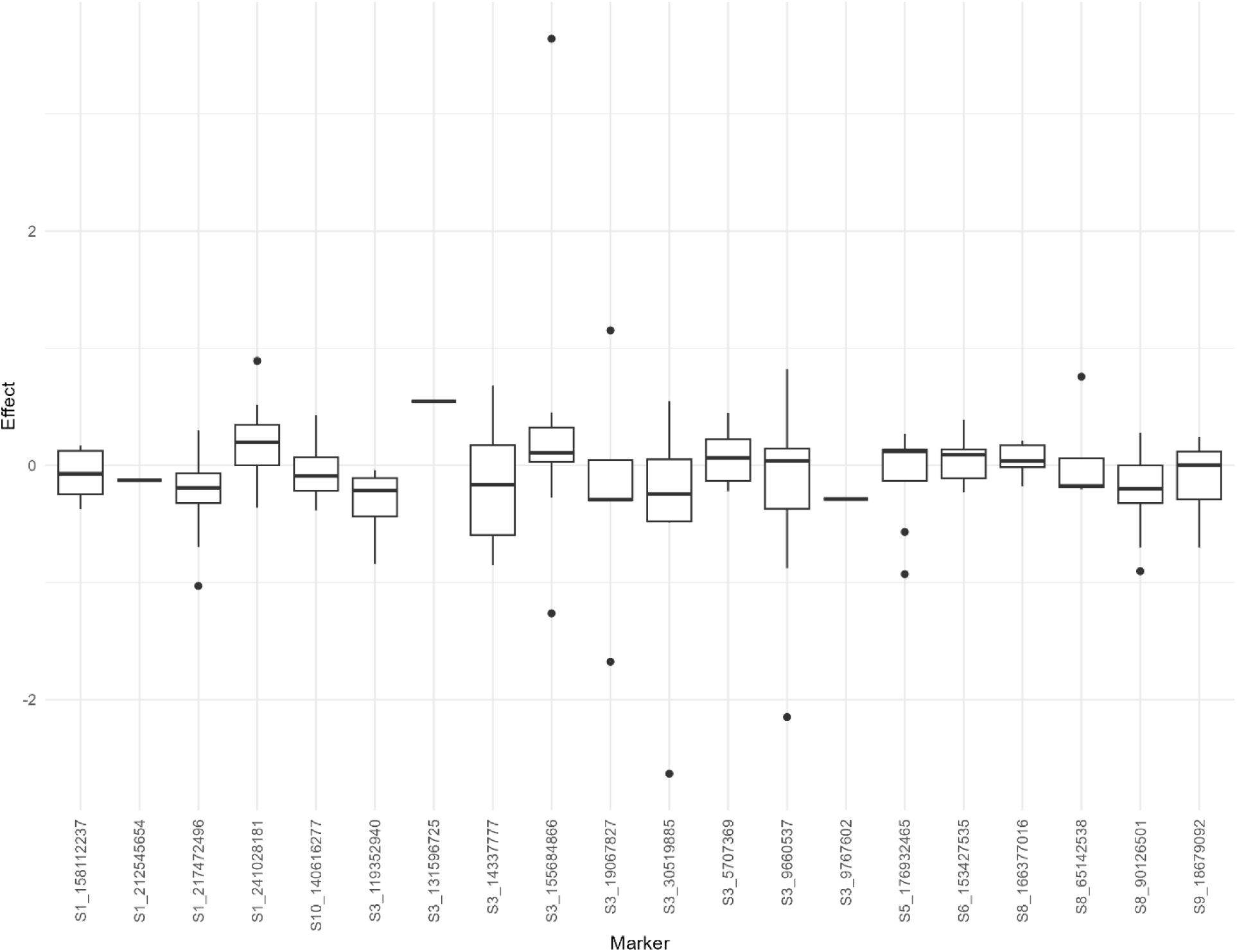
Boxplots of distribution of donor allele effects at QTL selected in the final multiple QTL model with variable donor allele effects for SLB. Negative effects correspond to increased resistance associated with introgressions.

## Supplementary Files

**File S1.** GBS SNP calls on nNILs filtered to remove markers with more than 20% missing data in HapMap format. Marker positions are in B73 v4 reference positions. Sample names were coded by sequence provider and require File S4 to translate to nNIL line names.

**File S2.** Chip genotyping data on NAM parents, a subset of 24 nNILs, and some additional lines in xlsx format. Marker positions are in B73 v3 reference positions.

**File S3.** Summary of introgression calling results of hidden Markov model for different combinations of parameter settings using chip genotyping data. Nir is non-informative rate, germ is genotyping error rate on homozygotes, gert is genotyping error rate on heterozygotes, p is probability that genotyping error on homozygote results in heterozygous call, r is recombination probability between adjacent markers, B73hetMean is proportion of B73 sample introgression calls that are heterozygous, DonorhetMean is proportion of donor line sample introgression calls that are heterozygous, NILhetMean is proportion of nNIL introgression calls that are heterozygous, B73homozIntroMean is proportion of B73 sample introgression calls that are homozygous introgressions, DonorhomozIntroMean is proportion of donor line sample introgression calls that are homozygous introgressions, NILhomozIntroMean is proportion of nNIL sample introgression calls that are homozygous introgressions, and PercNILsNoIntro is the percent of nNIL samples called with no introgressions.

**File S4.** Summary of introgression calling results of hidden Markov model for different combinations of parameter settings using GBS data. Nir is non-informative rate, germ is genotyping error rate on homozygotes, gert is genotyping error rate on heterozygotes, p is probability that genotyping error on homozygote results in heterozygous call, r is recombination probability between adjacent markers, mismatchMin, mismatchMean, and mismatchMax are the minimum, mean, and maximum mismatch rates between GBS calls and optimized chip calls across 24 nNIL samples, NILhomozIntroMean is proportion of nNIL sample introgression calls that are homozygous introgressions, NILhetMean is proportion of nNIL introgression calls that are heterozygous, and PercNILsNoIntro is the percent of nNIL samples called with no introgressions.

**File S5.** Founder matches for each nNIL based on SNP genotypes within introgression blocks and HapMap 3.2.1 data of NAM founders. Putative.donor is the donor indicated by line pedigree,

max.match is the maximum value of matching SNP proportions between the line and all potential donors, founder.match is the donor parent with highest match probability to the line; min.others, mean.others, max.others, and sd.others are minimum, mean, maximum, and standard deviation of match probabilities for all but the best match. The potential donor line columns are the specific match probabilities to each donor.

**File S6.** Summary of each introgression called. Chr is chromosome, Line is nNIL name, pos_leftflank is the position of the flanking marker adjacent to the first marker in the introgression, pos_start is the first SNP position of the introgression, pos_end is the last SNP position of the introgression, pos_righflank is the position of the flanking marker adjacent to the last marker in the introgression, marker_leftflank is the name of the left flanking marker, marker_start is the name of the first SNP, marker_end is the name of the last SNP, marker_rightflank is the name of the right flanking marker, chrom.size is the total length of the chromosome, intro.size is the length of the introgression, intro.prop.chr is the proportion of the chromosome represented by the introgression block. Positions are in AGPv4 B73 reference coordinates.

**File S7.** Phenotype data on a plot-basis. Geno is nNIL or check line name. EHT and PHT are ear and plant heights. DTA is days from planting to anthesis. GLS1 and GLS2 are first and second ratings of gray leaf spot disease (1 – 9 scale, 1 is most resistant). NLB1 and NLB2 are first and second ratings of northern leaf blight (percent leaf area infected). SLB1 and SLB2 are first and second ratings of southern leaf blight (1 – 9 scale, 1 is most resistant).

**File S8**. R markdown code for analysis of plot-level trait data to estimate trait heritabilities, genetic correlations across ratings, and line mean values for subsequent QTL testing.

**File S9.** Translation table for sample names in File S1 from sequencing provider codes to nNIL line names.

**File S10.** Python code to reformat and filter raw SNP data in preparation for hidden Markov model analysis.

**File S11.** Chip genotyping AGPv3 SNP positions in 6-column bed format.

**File S12**. Chip genotyping AGPv4 SNP position in bed format, generated from File S7 and the Ensembl assembly converter tool (https://plants.ensembl.org/Zea_mays/Tools/AssemblyConverter).

**File S13.** Python code to implement hidden Markov model to call introgressions from chip genotyping data, searching over a grid of parameter values.

**File S14.** R markdown code to summarize results of hidden Markov model applied to chip genotyping data for different combinations of parameter settings.

**File S15.** Python code to implement hidden Markov model to call introgressions from GBS data, searching over a grid of parameter values, and comparing results to a common subset of nNILs obtained from chip genotyping data, followed by calling introgressions on the full nNIL set using the optimal set of parameter settings.

**File S16.** R markdown code to summarize results of hidden Markov model applied to GBS data for different combinations of parameter settings.

**File S17.** Full set of nNIL introgression calls. For each marker, calls are 0,1, and 2, representing homozygous B73 haplotype, heterozygote, and homozygous introgression haplotype representing identity-by-descent from the donor parent.

**File S18.** R markdown code to compare chip SNP calls on nNILs to HapMap 3.2.1 NAM founder calls within introgression blocks.

**File S19.** Genotype call match proportions between each nNIL genotyped with SNP chip and NAM founder parents within introgression blocks.

**File S20.** R code to compare GBS SNP calls on nNILs to HapMap 3.2.1 NAM founder calls within introgression blocks.

**File S21.** Genotype call match proportions between each nNIL genotyped with GBS and NAM founder parents within introgression blocks.

**File S22.** R markdown code to summarize nNIL matches to founders within introgression blocks.

**File S23.** R markdown code to perform QTL tests.

## Notes

### Competing Interest Statement

The authors have declared no competing interest.

