## Supplementary material for "A maize near-isogenic line population designed for gene discovery and characterization of allelic effects": File S10

# -*- coding: cp1252 -*-
'''
Read in numericalized genotype calls on nNILs
Data from Morales et al. 2020 Plant J
GBS of nNILs, this is the raw SNP calls BEFORE FILLIN imputation
Note that sample IDs are in BGI format, have to translate those to NIL names (done below)
Numericalized to 1 for major allele homoz., 0 for minor homoz., 0.5 for het
In this script we will change the coding to 0 for major homoz, 1 for het, 2 for minor homoz
And set up and run one HMM model with best guess settings
Later can run the other script to grid search on HMM model settings.
'''
import os as os
import pandas as pd
import numpy as np
import time as time
import matplotlib.pyplot as plt


#os.chdir("Q:/.shortcut-targets-by-id/1FP9BlrAC2EltlqKM3ounRs2COTiTK12G/nNIL genotype data Jim and Tao")
os.chdir("C:/Users/jholland/Box/nNIL genotype data Jim and Tao")

#try to get the whole thing with pandas, check the timing on this

print("time to read into pandas")
start = time.process_time()
geno = pd.read_table("Data/nNIL_raw_SNPs_bgi_id_miss20.txt", sep= "\t", skiprows=1, header='infer')     #
print(time.process_time() - start)


#inspect the pandas df
geno.iloc[:10,:10]

#slice out the first row of marker names
marker_names = geno.columns.to_series()[1:]
marker_names[:5]

#read in the translation file that we can use to convert the BGI sample names to NIL IDs
sampleTranslate = pd.read_table("Data/bgi_nil_id.txt",   header=0,    encoding='windows-1252')

#merge the data frame with sampleTranslate, then drop the BGI sample IDs and use the NIL names
genob = pd.merge(sampleTranslate, geno, how='inner', left_on="bgi_id", right_on="<Marker>")
genob.drop(labels = ['bgi_id', '<Marker>'], axis = 1, inplace=True)

#slice out first columns of sample names
sample_names = genob.iloc[:,0]
sample_names[:5]

#keep just the numeric call columns/rows
genob = genob.iloc[:,1:]
genob.iloc[0:5,0:5]

###MAKE homozygous major allele = 0, het = 1, homoz. minor = 2
genomat = genob.to_numpy() # convert dataframe to numpy numeric array
genomat = np.multiply(np.subtract(genomat, 1), -2)

#get the marker minor allele freqs =  column mean values/2
geno_maf = np.multiply(np.nanmean(genomat, axis = 0), 0.5)
plt.hist(geno_maf)
#get min and max values of maf
np.min(geno_maf)
np.max(geno_maf)
#max is 0.54, I don't understand how it exceeds 0.5. min includes some 0.0 values, let's check those, they could be dropped
highmafs = geno_maf > 0.5
sum(highmafs) #only 21 of these, drop them

#ACTUALLY, we should filter markers with maf > 0.05, this is much higher than expected rate of 0.015
highmafs = geno_maf > 0.05
sum(highmafs) #only 21 of these, drop them

#remove fixed markers
zeromafs = geno_maf == 0
sum(zeromafs) #28641 markers with zero maf, these can be dropped

#filter the array to keep only markers with maf > 0 and maf < 0.1
marker_names_filt = marker_names[np.invert(zeromafs | highmafs)] #invert returns the converse of each element (True -> False)
geno_filt = genomat[:,np.invert(zeromafs | highmafs)]

#check heterozygosity rates at the markers, remove markers with overall heterozygosity > 2%
het_rates = np.nanmean(geno_filt == 1, axis = 0)
np.nansum(het_rates > 0.02)/len(het_rates)

#very few markers have high het rates, we will drop them.
#we expect het rate of 0.007813 in BC5F4 lines, so drop markers with > 2% het rate
geno_filt = geno_filt[:, het_rates < 0.02]
marker_names_filt = marker_names_filt[het_rates < 0.02]


#check heterozygosity rates by lines after filtering problematic markers
het_rates_lines = np.nanmean(geno_filt == 1, axis = 1)
np.nansum(het_rates_lines > 0.02)/len(het_rates_lines)
np.nansum(het_rates_lines > 0.02) #5 lines have high het rates
np.sort(het_rates_lines)[-7:]

#drop lines with high het rates
geno_filt = geno_filt[het_rates_lines < 0.02]
sample_names = sample_names[het_rates_lines < 0.02]

###################################
#WRITE UPDATED GENO FILE TO HARD DRIVE
#USING NUMPY SAVE
###################################
np.save('Q:/.shortcut-targets-by-id/1FP9BlrAC2EltlqKM3ounRs2COTiTK12G/nNIL genotype data Jim and Tao/Output/nNIL_filtered_geno_numpy', geno_filt, allow_pickle=True, fix_imports=True)

#can read this back in and start here if you need to stop and restart
#np.load('Q:/My Drive/TeoNIL/TeoNIL_filtered_geno_numpy.npy', allow_pickle=True)

marker_names_filt.to_csv('Q:/.shortcut-targets-by-id/1FP9BlrAC2EltlqKM3ounRs2COTiTK12G/nNIL genotype data Jim and Tao/Output/nNIL_filtered_marker_list', header = False, index = False)
sample_names.to_csv('Q:/.shortcut-targets-by-id/1FP9BlrAC2EltlqKM3ounRs2COTiTK12G/nNIL genotype data Jim and Tao/Output/nNIL_filtered_sample_list', header = False, index = False)

#Also make a subset of 39 nNILs for which we have array genotyping data and want to make comparisons
subsetList = pd.read_table('ListOf39nNILsChipSeqd.txt', header = None)

geno_filt_subset = geno_filt[sample_names.isin(subsetList.iloc[:,0]),:]
#turns out only 24 sample IDs match the subset, Im guessing the subset list is not properly formatted. But we can just use these 24

#update the sample_names to include just the subset for initial analysis below
sample_names = sample_names[sample_names.isin(subsetList.loc[:,0].tolist())]

np.save('Q:/.shortcut-targets-by-id/1FP9BlrAC2EltlqKM3ounRs2COTiTK12G/nNIL genotype data Jim and Tao/Output/nNIL_filtered_geno_numpy_24subset', geno_filt_subset, allow_pickle=True, fix_imports=True)
