## Supplementary material for "A maize near-isogenic line population designed for gene discovery and characterization of allelic effects": File S13

# -*- coding: utf-8 -*-
"""
Created on Tue May 21 17:36:31 2024

@author: jholland

Read data from SNP chip on nNILs and putative parents
Convert marker positions from AGP V3 to AGP V4
Run the HMM to call introgressions from the chip data
These data are different from GBS because much higher quality
So alignment/sequencing errors should be very low rate
But still have problem of non-informative markers that confuse the introgression calls
And for this purpose we do not want to assume the putative donor is correct.
Rather, make the introgression calls, then compare individual SNP calls within introgression blocks to the possible parents

"""

import os as os
import sys as sys
import pandas as pd
import numpy as np

# adding folder with calIntrogressions module to the system path
sys.path.insert(0, 'Q:/.shortcut-targets-by-id/1FP9BlrAC2EltlqKM3ounRs2COTiTK12G/nNIL genotype data Jim and Tao/Code')
import callIntrogressions as ci

os.chdir("Q:/.shortcut-targets-by-id/1FP9BlrAC2EltlqKM3ounRs2COTiTK12G/nNIL genotype data Jim and Tao")

#GET THE CHIP GENOTYPING DATA FOR A SUBSET OF LINES THAT WE WILL USE AS GOLD STANDARD
chip_header = pd.read_excel("Data/Chip data of NAM parents and nNILs.xlsx",  header = None, nrows = 1)
chip_lines = chip_header.loc[0,9:] #get the line names

chip = pd.read_excel("Data/Chip data of NAM parents and nNILs.xlsx", header = 0, skiprows = 1)

# fix column names
chip.rename(columns={'#CHROM':'chr', 'POS (V3)':'pos_V3'}, inplace = True)
chip.rename(columns = dict(zip(chip.columns[9:], chip_lines)), inplace = True)

# chip data file has V3 positions, update to V4 to match the GBS data of Morales et al
chipV3pos = pd.read_table("Output/nNIL_chip_SNP_positions_V3_6col.bed", header = None)
chipV3pos.columns = ['chr', 'startV3', 'pos_V3',  'name', 'score', 'strand']
chipV3pos.drop(columns = ['startV3', 'score', 'strand'], inplace = True)
chipV4pos = pd.read_table("Data/nNIL_chip_SNP_positions_converted_to_V4.bed", header = None)

chipV4pos.columns = ['chr_V4', 'startV4', 'pos_V4',  'name', 'score', 'strand']
chipV4pos.drop(columns = ['startV4', 'score', 'strand'], inplace = True)

chipV3toV4 = pd.merge(chipV3pos, chipV4pos, how = 'inner', on = "name")
                                                  
#find snps in V3 but not in V4 just for curiosity
#set(chipV3pos['name']).difference(set(chipV4pos['name']))
#yep, they are just missing from the V4 positions

#remove any SNPs not in chr 1 - 10, some map to contigs in V4, get rid of them
chipV3toV4 = chipV3toV4.loc[chipV3toV4.chr_V4.isin(['1', '2', '3', '4', '5', '6', '7', '8', '9', '10'])]
chip = chip.loc[chip.chr.isin(['1', '2', '3', '4', '5', '6', '7', '8', '9', '10'])]

#chipV3toV4 = chipV3toV4.astype({'pos_V3':'int64'}) #'chr':'int64', 
#merge the position translation data frame with the chip data frame
chip = chip.astype({'chr':'int64'})
chip = chip.merge(chipV3toV4, on = ['chr', 'pos_V3'])
chip['name'] = 'S' + chip['chr_V4'].astype('str') + '_' + chip['pos_V4'].astype('str')


#transpose the data frame so lines on rows and markers on columns
chipT = chip.iloc[:,9:(chip.shape[1]-3)].transpose()
chipT.columns = chip['name']

#drop the extra lines that I am not going to analyze
chipT.drop(index = ['H100', 'Ki3', 'NC262', 'NC304', 'DRIL32.90 ', 'DRIL32.095',
'DRIL52.055', 'DRIL62.078', 'Mo17', 'NIL-1030', 'NIL-1290'], inplace = True)

#convert the genotype calls to numeric 0,1,2
def converter(x):
    if x == '0/0': return('0')
    if x == '0/1' or x == '1/0': return('1')
    if x == '1/1': return('2')
    if x == './.' or x == 'nan' or pd.isna(x): return('3') #3 will be na value
  
chipT2 = chipT.applymap(converter)
#convert to numpy array
chipNp = chipT2.to_numpy(dtype='int64')

#Any markers scored as non-zero on B73 samples? (But also < 3! because na is now 3)
B73_nobs = np.sum(chipNp[0:2,] != 3, axis = 0)
B73_nobs[B73_nobs == 0] = 1 #replace 0 counts with so next division will work, we will not drop these markers
B73_afs = np.divide(np.sum(chipNp[0:2,], axis = 0) - 3*np.sum(chipNp[0:2,] == 3, axis = 0),(2*B73_nobs)) #have to use max 1 or sum in case sum is zero, we don't want to drop those markers
sum(B73_afs > 0) #349

# check introgression rates in NILs at markers remaining. If some are high, maybe the B73 recurrent parent does not match the reference
# in which case we should drop those markers

chipNIL = chipNp[chipT.index.str.contains(r'(NIL)'), ]
NIL_nobs = np.sum(chipNIL != 3, axis = 0)
NIL_nobs[NIL_nobs == 0] = 1 #replace 0 counts with so next division will work, we will not drop these markers
NIL_afs = np.divide(np.sum(chipNIL, axis = 0) - 3*np.sum(chipNIL == 3, axis = 0),(2*NIL_nobs)) #have to use max 1 or sum in case sum is zero, we don't want to drop those markers
sum(NIL_afs > 0.20) #424


#remove markers with non-zero scores on either B73 sample,
#remove markers with NIL mafs > 0.2, I think these may be sites where recurrent B73 doesn't match the referece
#also drop one of the B73 samples
chipNp = chipNp[1::,np.logical_and(B73_afs == 0, NIL_afs < 0.20)]

chipSamples = chipT.index.to_series()
chipSamples = chipSamples[1:,]
chipMarkers = chipT.columns.to_series()[np.logical_and(B73_afs == 0, NIL_afs < 0.20)]

###############################
#Find overlapping markers between chip and GBS
gbsMarkers = pd.read_table('Output/nNIL_filtered_marker_list', header = None)
gbsMarkers = gbsMarkers.iloc[:,0]

chipMarkersInGBS = chipMarkers[chipMarkers.isin(gbsMarkers)]
#yikes only 9 are exact matches!!!!

#so, instead, we need to identify NEAREST GBS markers to each gbsMarker and make a common name 
chrnames = gbsMarkers.str.extract(r'(S\d*)')
chrnames = chrnames.iloc[:,0].str.replace('S','').astype('int64')

posnames = gbsMarkers.str.extract(r'(\d*$)')
posnames = posnames.iloc[:,0].astype('int64')

gbsMarkersDF = pd.DataFrame({'name' : gbsMarkers, 'chr': chrnames, 'pos':posnames})

chipMarkersDF = chip[['name', 'chr_V4', 'pos_V4']]
chipMarkersDF.drop(chipMarkersDF.loc[~chipMarkersDF['name'].isin(chipMarkers)].index, inplace = True)
chipMarkersDF.rename(columns = {'chr_V4':'chr', 'pos_V4':'pos'}, inplace = True)
chipMarkersDF.chr = chipMarkersDF.chr.astype('int64')

#now iterate through chip markers, searching for closest position match on same chrom from GBS markers
def closestMatch(row):
    gbs = gbsMarkersDF.loc[gbsMarkersDF.chr == row['chr'],:]
    dist = abs(gbs.pos - row['pos'])
    dist.reset_index(inplace = True, drop = True)
    return(gbs.iloc[dist.idxmin(),0])

chipMarkersDF['closestGBS'] = chipMarkersDF.apply(closestMatch, axis=1)
chipMarkersDF['closest_gbs_pos'] = chipMarkersDF.closestGBS.str.extract(r'(\d*$)')
chipMarkersDF.closest_gbs_pos = chipMarkersDF.closest_gbs_pos.astype('int64')
chipMarkersDF['distance'] = abs(chipMarkersDF['closest_gbs_pos'] - chipMarkersDF['pos'])
chipMarkersDF.to_csv("Output/nNIL chip markers and closest GBS markers.csv",  index = False)

###############################


###############################
#Implement HMM introgression caller on chip genotyping data

#estimate probability that a non-B73 haplotype has a different SNP call than B73
#we can do this by measuring the average frequency of non-B73 calls among putative donor parents
donorCalls = chipNp[~chipSamples.str.contains(r'(NIL)') & ~chipSamples.str.contains(r'B73'),:] 
#for computing non-informative rate, we have to re-convert '3' values to NAs
donorCalls = donorCalls.astype('float')
donorCalls[donorCalls == 3] = np.nan
donor_maf = np.multiply(np.nanmean(donorCalls, axis = 1), 0.5)
maf = np.nanmean(donor_maf)

#compute the 'non-informative rate' as 1 - maf among donors (the expected rate at which a non-B73 haplotpye has an IIS B73 SNP)
nir = 1 - maf

#recode NA values as integer 3
#note chipNp is a numpy array not pandas df.
#as such it has nan values not NaN
#we will encode our HMM with output value 3 as one observable state
chipNp = np.nan_to_num(chipNp, copy=True, nan=3)

#estimate missing data rate
#missing data rate empirically sets the probability that any observed state is missing, we assume it's constant probability for all true states
missing = chipNp == 3
missing_rate = missing.sum().sum()/(chipNp.shape[0]*chipNp.shape[1])


#The average recombination frequency between adjacent SNPs assuming total map length of 1500 cM:
#multiply by two in numerator because we have effectively two meioses in the backcrossing and selfing 
#breeding method (similar to recombination freq doubling in RILs)
avg_r = 2*1500/(100*len(chipMarkers)) 

#Get the chr for each SNP
#get the SNP position information from the name

chroms = chipMarkers.replace("_.+$", "", regex = True)
chroms.replace("^S", "", regex = True, inplace = True)
chroms.reset_index(inplace = True, drop = True)
chroms[:10]

#split the geno data frame into ten separate dataframes, one for each chromosome
#each chromosome will need to be processed separately in next step

markers_by_chrom = {}
for i in range(1,11):
    markers_by_chrom[i] = chroms[chroms == str(i)].index

# create indices for parental lines vs NILs
# we need this info to separately summarize introgression call results for the two groups
B73Index = chipSamples.index[chipSamples == "B73"]
parentIndices = chipSamples.index[~chipSamples.str.contains(r'(NIL)') & ~chipSamples.str.contains(r'B73')]
NILIndices = chipSamples.index[chipSamples.str.contains(r'(NIL)') | chipSamples.str.contains(r'B73 ') ]


# make a function to report qc stats for each parameter setting
def evaluate_chip_results(NIL_calls, chipSamples, B73Index, parentIndices, NILIndices, nir, germ, gert, p, r):
    
    #get the %het calls in B73
    perc_het_B73 = np.count_nonzero(NIL_calls[chipSamples.isin(B73Index),:] == 1, axis = 1)/NIL_calls.shape[1]
       
    #get the %het calls per donor parent line
    perc_het_donor = np.count_nonzero(NIL_calls[chipSamples.isin(parentIndices),:] == 1, axis = 1)/NIL_calls.shape[1]
 
    #get the %het calls per NIL
    perc_het_NIL = np.count_nonzero(NIL_calls[chipSamples.isin(NILIndices),:] == 1, axis = 1)/NIL_calls.shape[1]  
 
    #get the % homozygous introgression calls in B73
    perc_homoz_intro_B73 = np.count_nonzero(NIL_calls[chipSamples.isin(B73Index),:] == 2, axis = 1)/NIL_calls.shape[1]
       
    #get the % homozygous introgression calls per donor parent line
    perc_homoz_intro_donor = np.count_nonzero(NIL_calls[chipSamples.isin(parentIndices),:] == 2, axis = 1)/NIL_calls.shape[1]
 
    #get the % homozygous introgression calls per NIL
    perc_homoz_intro_NIL = np.count_nonzero(NIL_calls[chipSamples.isin(NILIndices),:] == 2, axis = 1)/NIL_calls.shape[1]
    
    #get % of NILs that have no introgressions
    perc_lines_no_intro_NIL = 1 - np.count_nonzero(np.count_nonzero(NIL_calls[chipSamples.isin(NILIndices),:] > 0, axis = 1))/len(NILIndices)
                                         
    #create a dictionary to hold the current introcalls and summaries
    # current_results = {"calls":NIL_calls, "NILhet":perc_het, "NILhetMean":np.mean(perc_het),
    #                    "NILhomoz":perc_homoz_intro, "NILhomozIntroMean":np.mean(perc_homoz_intro), 
    #                    "PercNILsNoIntro":perc_lines_no_intro}

    current_results = {"Method":"Chip",
                       "nir":nir,
                       "germ":germ,
                       "gert":gert,
                       "p":p,
                       "r":r,
                       "B73hetMean":np.mean(perc_het_B73),
                       "DonorhetMean":np.mean(perc_het_donor),
                       "NILhetMean":np.mean(perc_het_NIL),
                       "B73homozIntroMean":np.mean(perc_homoz_intro_B73), 
                       "DonorhomozIntroMean":np.mean(perc_homoz_intro_donor), 
                       "NILhomozIntroMean":np.mean(perc_homoz_intro_NIL), 
                       "PercNILsNoIntro":perc_lines_no_intro_NIL}
    return(current_results)


#f(donor_hom) = 0.011179
f_2 = 0.011179
#f(het) = 0.007813
f_1 = 0.007813
#f(B73 homoz) = 1 - others

#Run the function on a range of nir, scer, and avg_r values
parameter_tests = {}
for nir in [0.001, 0.01, 0.1, 0.3, 0.5, 0.7, 0.9]:
    for germ in [0.0001, 0.001, 0.01]:
        for gert in [0.0001, 0.001, 0.01]:
            for p in [0.1, 0.25, 0.5, 0.75, 0.9]:
                for r in [avg_r/2, avg_r, avg_r*2]:
                    calls = ci.call_intros(chipNp, markers_by_chrom, nir, germ, gert, p, missing_rate, r, f_1, f_2, return_calls = True)
                    summary = evaluate_chip_results(calls, chipSamples, B73Index, parentIndices, NILIndices, nir, germ, gert, p, r)
                    parameter_tests["nir " +str(round(nir,5)) + " germ " + str(round(germ,4)) + " gert " + str(round(gert,4)) + " p " + str(round(p,3)) + " r " + str(round(r,6))] = summary
                    #call_intros will return a list of tuples, each tuple will have two data frames, one with the calls, other with parameter settings
        

summaryDF = pd.DataFrame.from_dict(parameter_tests, orient='index')    

# summaryDF['nir'] = summaryDF.index.str.extract("(\d+\.\d+).+(\d+\.\d+).+(\d+\.\d+)")[0].values
# summaryDF['scer'] = summaryDF.index.str.extract("(\d+\.\d+).+(\d+\.\d+).+(\d+\.\d+)")[1].values
# summaryDF['r'] = summaryDF.index.str.extract("(\d+\.\d+).+(\d+\.\d+).+(\d+\.\d+)")[2].values

# #reorder columns
# newcols = ['nir', 'scer', 'r', 'B73homozIntroMean', 'B73hetMean', 'DonorhetMean', 'DonorhomozIntroMean',
#            'NILhetMean', 'NILhomozIntroMean', 'PercNILsNoIntro', ]
# summaryDF = summaryDF[newcols]


#Write the summaryDF to a csv so we can visualize results in R
summaryDF.to_csv('Output/nNIL_chipdata_HMMgridSearchSummary.csv', index=False)


# choose parameter settings that give the best results and make final calls:
finalModel = ci.call_intros(geno = chipNp, marker_dict = markers_by_chrom, nir = 0.9, germ = 0.01, gert = 0.001, p = 0.1, mr = missing_rate, r = avg_r/2, f_1 = f_1, f_2 = f_2, return_calls = True)

#convert numpy array of calls to data frame
finalModel = pd.DataFrame(finalModel,
                            index = chipSamples,
                            columns = chipMarkers)
finalModel.index.name = "Line"

finalModel.to_csv('Output/nNIL_chipdata_HMM_introgressionCalls.csv')

#last step is to project the introgression calls onto the nearest GBS markers
# drop chip markers that project to the same GBS marker as another chip marker
chipMarkersUnique = chipMarkersDF.groupby(['chr', 'closestGBS'], as_index = False).first()
chipMarkersUnique.sort_values(by = ['chr', 'pos'], inplace = True)

NIL_calls_projected = finalModel.loc[chipSamples.isin(NILIndices),chipMarkers.isin(chipMarkersUnique['name'])]
#now rename the markers to the nearest GBS marker, not exactly correct, but should be good approximation for IBD calls
NIL_calls_projected.columns = chipMarkersUnique['closestGBS']
NIL_calls_projected.index.name = "Line"

NIL_calls_projected.to_csv('Output/nNIL_chipdata_HMM_introgressionCalls_project_to_GBS_markers.csv')
