## Supplementary material for "A maize near-isogenic line population designed for gene discovery and characterization of allelic effects": File S14

---
title: "HMM grid search results chip data"
author: "Jim Holland"
date: "2024-08-16"
output: html_document
---

```{r setup, include=T}
knitr::opts_chunk$set(echo = TRUE)
library(tidyverse)
#base.path = "Q:/.shortcut-targets-by-id/1FP9BlrAC2EltlqKM3ounRs2COTiTK12G/nNIL genotype data Jim and Tao/Output/"
base.path = "C:/Users/jholland/Box/nNIL genotype data Jim and Tao/Output/"
```


Get the grid search results for HMM on chip data.  
Compute the percent of genome called as homoz non-introgression in donor lines as 1 - (homoz + het introgression). We expect these values to be low. This way we can graph all of the statistics in a single plot (otherwise homoz introgression values are near 1, other values are near zero and scaling is awkward).
```{r}
chip.results = read.csv(paste0(base.path, "nNIL_chipdata_HMMgridSearchSummary.csv")) %>%
  mutate(DonorhomozB73 = 1 - DonorhetMean - DonorhomozIntroMean) %>%
  select(-Method) 
results.long = pivot_longer(chip.results, cols = -c("nir", "germ", "gert", "p", "r"), names_to = "Statistic", values_to = "Proportion") %>%
  mutate(nir = as.factor(nir),
         germ = as.factor(germ),
         gert = as.factor(gert),
         p = as.factor(p),
         r = as.factor(r),
         Statistic = sub("Mean", "", Statistic))
```


Graph results for statistics expected to be near zero
```{r}
results.near.zero = results.long %>%
  filter(Statistic %in% c("B73homozIntro", "B73het", "Donorhet", "NILhet", "NILhomozIntro", "PercNILsNoIntro", "DonorhomozB73"))
ggplot(results.near.zero, aes(x = nir, y = Proportion)) +
  geom_point(aes(colour = gert, shape = p)) +
  facet_grid(rows = vars(germ), cols = vars(Statistic)) +
  scale_color_discrete(name=c(expression(ger[t]))) +
  theme_minimal() +
  theme(axis.text.x = element_text(angle = 90, vjust = 0.5, hjust=1))  + 
  xlab("Non-informative rate (nir) of SNPs") +
  ggtitle("HMM grid search results of chip data. Genotyping error rate on homozygotes in rows") 
```
Write plot to a file
```{r}
ggsave(filename = paste0(base.path, "nNIL chip data hmm grid search results.tiff"), width = 10, height = 8, units = "in", dpi = 300, bg = "white")
ggsave(filename = paste0(base.path, "nNIL chip data hmm grid search results.png"), width = 10, height = 8, units = "in", dpi = 300, bg = "white")

```

Find the optimal parameter settings
```{r}
best.settings = chip.results %>%
  arrange(desc(DonorhomozIntroMean)) %>%
  slice_head(n = 5)
```
