## Supplementary material for "A maize near-isogenic line population designed for gene discovery and characterization of allelic effects": File S15

# -*- coding: utf-8 -*-
"""
Created on Thu May 23 21:22:36 2024

@author: jholland

Read data from GBS on nNILs 
Grid search around parameters
Run the HMM to call introgressions, compare to introgressions already called from the chip data on a subset
Compute identity of calls between GBS and chip to find best parameter settings

"""
import os as os
import sys as sys
import pandas as pd
import numpy as np


# adding folder with calIntrogressions module to the system path
sys.path.insert(0, 'Q:/.shortcut-targets-by-id/1FP9BlrAC2EltlqKM3ounRs2COTiTK12G/nNIL genotype data Jim and Tao/Code')

import callIntrogressions as ci

os.chdir("Q:/.shortcut-targets-by-id/1FP9BlrAC2EltlqKM3ounRs2COTiTK12G/nNIL genotype data Jim and Tao")

#read in GBS data on full NIL set
gbs = np.load('Output/nNIL_filtered_geno_numpy.npy', allow_pickle=True)

marker_names = pd.read_csv('Output/nNIL_filtered_marker_list', header = None).loc[:,0]
sample_names = pd.read_csv('Output/nNIL_filtered_sample_list', header = None).loc[:,0]

#read in the introgression calls on NILs based on chip data
#but marker positions have been updated to V4 and only the set closest to GBS markers retained
#so markers mostly don't match GBS positions exactly but are nearest and renamed to match nearest GBS marker name

NIL_calls_chip = pd.read_csv('Output/nNIL_chipdata_HMM_introgressionCalls_project_to_GBS_markers.csv', index_col = 0)

#subset the chip calls to the NILs also included in the GBS data
NIL_calls_chip = NIL_calls_chip.loc[NIL_calls_chip.index.isin(sample_names),:]

#sort the chip calls by sample name in index
NIL_calls_chip.sort_index(inplace = True)

#subset the GBS data to the same lines in the chip data
gbs2 = gbs[sample_names.isin(NIL_calls_chip.index),:]
sample_names_subset = sample_names[sample_names.isin(NIL_calls_chip.index)]

#check that sample order is identical between GBS and chip samples 
all(sample_names_subset == NIL_calls_chip.index) #if not True, stop and fix!

#convert chip dataframe to numpy array
chip = np.array(NIL_calls_chip)

#keep track of the column names (markers) in a list since numpy array does not have column names
chip_markers = NIL_calls_chip.columns.to_list()

#COMPUTE MAF ON FULL DATA SET
#AND ESTIMATE PROPORTION OF NON-INFORMATIVE CALLS = 'nir'

#most sites will match B73 even if they are introgressed because teo and maize share most alleles
# rate of SNP being non-informative is ?
# we expect across the genome that donor allele frequency in BC5 NILs should be 0.0156
#we actually observe this average allele frequency after filtering:
geno_maf = np.multiply(np.nanmean(gbs2, axis = 0), 0.5)
maf = np.nanmean(geno_maf)
print("non-reference allele frequency across all nNILs: " + str(maf))

#compute the 'non-informative rate' as the proportional difference between the expected maf and the observed maf
nir = max((0.0151 - maf)/0.0151, 0.001) #if <=0 set to a small number.
print("estimate of non-informative SNP rate based on proportional difference between expected and observed non-reference allele frequency: ")
print(str(nir))


#recode NA values as integer 3
#note chipNp is a numpy array not pandas df.
#as such it has nan values not NaN
#we will encode our HMM with output value 3 as one observable state
gbs2 = np.nan_to_num(gbs2, copy=True, nan=3)

#estimate missing data rate
#missing data rate empirically sets the probability that any observed state is missing, we assume it's constant probability for all true states
missing = gbs2 == 3
missing_rate = missing.sum().sum()/(gbs2.shape[0]*gbs2.shape[1])


#for the HMM we need a vector of hidden states, a vector of observed states (in our case the same thing), 
#a transition probability matrix and an emmision probability matrix

states = [0, 1, 2] #"B73", "het",  "donor"

#The average recombination frequency between adjacent SNPs assuming total map length of 1500 cM:
#multiply by two in numerator because we have effectively two meioses in the backcrossing and selfing 
#breeding method (similar to recombination freq doubling in RILs)
avg_r = 2*1500/(100*len(marker_names)) #factor of 2 here is because we have ~ equivalent of 2 generations of meioses during backcrossing and selfing to homozygosity, per Haldane and Waddington
print("average recombination frequency between adjacent markers: " + str(avg_r))


#Get the chr for each SNP
#get the SNP position information from the name

chroms = marker_names.replace("_.+$", "", regex = True)
chroms.replace("^S", "", regex = True, inplace = True)
chroms.reset_index(inplace = True, drop = True)
chroms[:10]

#split the geno data frame into ten separate dataframes, one for each chromosome
#each chromosome will need to be processed separately in next step

markers_by_chrom = {}
for i in range(1,11):
    markers_by_chrom[i] = chroms[chroms == str(i)].index

# make a function to compare NIL calls from gbs and chip data sets
def compare_gbs_chip(NIL_calls, chip_data, chip_markers, nir, germ, gert, p, r):

    
    #compare the mismatch rate between NIL_calls and chip data
    NIL_calls_sub = NIL_calls[:,marker_names.isin(chip_markers)]
    NIL_vs_chip = NIL_calls_sub != chip_data
    
    #compute mismatch rate by line
    mismatches = np.nanmean(NIL_vs_chip, axis = 1)/NIL_vs_chip.shape[0]
    
    #compute min, mean, max mismatch rates
    mismatchMin = np.min(mismatches)
    mismatchMean = np.mean(mismatches)
    mismatchMax = np.max(mismatches)

    #summarize introgression calls using all markers (no need to subset to chip set here)
    #get the %het calls per NIL
    perc_het_NIL = np.count_nonzero(NIL_calls == 1, axis = 1)/NIL_calls.shape[1]  
 
    #get the % homozygous introgression calls per NIL
    perc_homoz_intro_NIL = np.count_nonzero(NIL_calls == 2, axis = 1)/NIL_calls.shape[1]
    
    #get % of NILs that have no introgressions
    perc_lines_no_intro_NIL = 1 - np.count_nonzero(np.count_nonzero(NIL_calls > 0, axis = 1))/NIL_calls.shape[0]
                                         
    current_results = {
                       "nir":nir,
                       "germ":germ,
                       "gert":gert,
                       "p":p,
                       "r":r,       
                       "mismatchMin":mismatchMin,
                       "mismatchMean":mismatchMean,
                       "mismatchMax":mismatchMax,
                       "NILhetMean":np.mean(perc_het_NIL),
                       "NILhomozIntroMean":np.mean(perc_homoz_intro_NIL), 
                       "PercNILsNoIntro":perc_lines_no_intro_NIL}
    

    return(current_results)


#f(donor_hom) = 0.011179
f_2 = 0.011179
#f(het) = 0.007813
f_1 = 0.007813
#f(B73 homoz) = 1 - others

#Run the function on a range of nir, scer, and avg_r values
parameter_tests = {}
for nir in [0.001, 0.01, 0.1, 0.3, 0.5, 0.7, 0.9]:
    for germ in [0.0001, 0.001, 0.01]:
        for gert in [0.0001, 0.001, 0.01]:
            for p in [0.1, 0.25, 0.5, 0.75, 0.9]:
                for r in [avg_r/2, avg_r, avg_r*2]:
                    calls = ci.call_intros(gbs2, markers_by_chrom, nir, germ, gert, p, missing_rate, r, f_1, f_2, return_calls = True)

                    summary = compare_gbs_chip(calls, chip, chip_markers, nir, germ, gert, p, r)
                    parameter_tests["nir " +str(round(nir,5)) + " germ " + str(round(germ,4)) + " gert " + str(round(gert,4)) + " p " + str(round(p,3)) + " r " + str(round(r,6))] = summary
                    #call_intros will return a list of tuples, each tuple will have two data frames, one with the calls, other with parameter settings
        

summaryDF = pd.DataFrame.from_dict(parameter_tests, orient='index')    
#extract the summary values only from the parameter_tests object
 
# summaryDF['nir'] = summaryDF.index.str.extract("(\d+\.\d+).+(\d+\.\d+).+(\d+\.\d+)")[0].values
# summaryDF['scer'] = summaryDF.index.str.extract("(\d+\.\d+).+(\d+\.\d+).+(\d+\.\d+)")[1].values
# summaryDF['r'] = summaryDF.index.str.extract("(\d+\.\d+).+(\d+\.\d+).+(\d+\.\d+)")[2].values

#reorder columns
newcols = ['nir', 'germ', 'gert', 'p', 'r', 'mismatchMin', 'mismatchMean', 'mismatchMax', 'NILhomozIntroMean', 'NILhetMean', 'PercNILsNoIntro']
summaryDF = summaryDF[newcols]


#Write the summaryDF to a csv so we can visualize results in R
summaryDF.to_csv('Output/nNIL_gbs_vs_chip_data_HMMgridSearch.csv', index=False)

#################################### 
#Save GBS introgression calls from best parameter setting for 24 subset
#################################### 

#nir = 0.9 germ = 0.01 gert = 0.0001 p = 0.9 and r = avg_r seems to give lowest mismatch rate
finalModel = ci.call_intros(geno = gbs2, marker_dict = markers_by_chrom, nir = 0.9, germ = 0.01, gert = 0.0001, p = 0.9, mr = missing_rate,  r = avg_r, f_1 = f_1, f_2 = f_2, return_calls = True)

#convert numpy array of calls to data frame
finalModelDF = pd.DataFrame(finalModel,
                           index = sample_names_subset,
                           columns = marker_names)

finalModelDF.to_csv('Output/nNIL_gbs_HMM_introgressionCalls_24subset.csv')

#################################### 
#Save GBS introgression calls from best parameter setting for 24 subset, markers nearest to chip set only
#################################### 

#subset the markers to make comparison to the chip calls easier
finalModelSub = pd.DataFrame(finalModel[:,marker_names.isin(chip_markers)],
                           index = sample_names_subset,
                           columns = marker_names[marker_names.isin(chip_markers)])

finalModelSub.to_csv('Output/nNIL_gbs_HMM_introgressionCalls_chip_marker_24subset.csv')


#################################### 
#Save GBS introgression calls from best parameter setting for FULL set of nNILs
#################################### 

#use final parameter settings to call introgressions on the full set of nNILs
gbs = np.nan_to_num(gbs, copy=True, nan=3)

#estimate missing data rate on full set and use this value for final introgression calling
#missing data rate empirically sets the probability that any observed state is missing, we assume it's constant probability for all true states
missing = gbs == 3
missing_rate = missing.sum().sum()/(gbs.shape[0]*gbs.shape[1])

#nir = 0.9 germ = 0.01 gert = 0.0001 p = 0.9, r = avg_r, and mean missing rate obtained from full set used for final introgression calls
finalModelFull = ci.call_intros(geno = gbs, marker_dict = markers_by_chrom, nir = 0.9, germ = 0.01, gert = 0.0001, p = 0.9, mr = missing_rate,  r = avg_r, f_1 = f_1, f_2 = f_2, return_calls = True)

finalModelFullDF = pd.DataFrame(finalModelFull,
                           index = sample_names,
                           columns = marker_names)

finalModelFullDF.to_csv('Output/nNIL_gbs_HMM_introgressionCalls_full_set.csv')
