## Supplementary material for "A maize near-isogenic line population designed for gene discovery and characterization of allelic effects": File S18

---
title: "Compare Chip SNPs in introgressions to HapMap3 subset of nNILs"
author: "Jim Holland"
date: "2024-06-17"
output: html_document
---

```{r setup, include=FALSE}
knitr::opts_chunk$set(echo = TRUE)
library(data.table)
library(tidyverse)
library(vcfR)
base.path = "Q:/.shortcut-targets-by-id/1FP9BlrAC2EltlqKM3ounRs2COTiTK12G/nNIL genotype data Jim and Tao/"
```

Get chip introgression calls on a subset of nNILs
```{r}
chip.intro = read.csv(paste0(base.path, "Output/nNIL_chipdata_HMM_introgressionCalls_project_to_GBS_markers.csv")) 
names(chip.intro)[1] = 'sampleID'
chip.intro = chip.intro %>% mutate(
  sampleID = gsub(" ", "_", sampleID))
```

Get chip SNP calls on same nNILs
```{r}
chip.snp = readxl::read_xlsx(paste0(base.path,"Data/Chip data of NAM parents and nNILs.xlsx"), col_names = T, skip = 1)
chip.snp.header = readxl::read_xlsx(paste0(base.path,"Data/Chip data of NAM parents and nNILs.xlsx"), col_names = F, n_max = 1)
chip.snp.header = unlist(chip.snp.header[1,]) #make it a vector
chip.snp.header.dups = duplicated(chip.snp.header) #find the duplicated column names
chip.snp.header[chip.snp.header.dups] = paste0(chip.snp.header[chip.snp.header.dups], "_2") #give B73 and Ki3 replicates unique identifiers

#check if any duplicate column names remain
any(duplicated(chip.snp.header))

chip.snp.header = gsub(" ", "_", chip.snp.header) #replace spaces with _ in NIL names
names(chip.snp)[10:ncol(chip.snp)] = chip.snp.header
```
Chip data are not base calls, they are ref/alt calls, need to reformat all of this to IUPAC base calls. Also drop indels and some weird markers that are all missing data
```{r}
#First drop markers that are INDELs
chip.snp = chip.snp %>%
    filter(nchar(REF) == 1 & nchar(ALT) == 1) %>%
  filter(REF != "N" | REF != "." & ALT != "N" & ALT != ".")

vcf_to_calls = function(x, ref, alt){
  if (is.na(x)) {return(NA)}
  if (x == "0/0") {return(ref)}
  if (x == "1/1") {return(alt)}
  if (x == "0/1" | x == "1/0" | x == "0|1" | x == "1|0") {
    het = paste0(ref, "/", alt)
    if (het == "A/C" | het == "C/A") {return("M")}
    if (het == "A/G" | het == "G/A") {return("R")}
    if (het == "A/T" | het == "T/A") {return("W")}
    if (het == "C/G" | het == "G/C") {return("S")}
    if (het == "C/T" | het == "T/C") {return("Y")}
    if (het == "G/T" | het == "T/G") {return("K")}
  }
  return(NA) #if all else fails
}


vcf_to_calls_by_row = function(row){
  #row is one row of the chip.snp data frame
  ref = row["REF"]
  alt = row["ALT"]
  calls = row[10:length(row)]
  sapply(calls, vcf_to_calls, ref = ref, alt = alt) 
}

chip.snp.calls = apply(chip.snp, 1, vcf_to_calls_by_row)
#this returns matrix rows = samples, cols = markers
row.names(chip.snp.calls) = colnames(chip.snp)[-c(1:9)]
colnames(chip.snp.calls) = paste0("S", chip.snp$`#CHROM`, "_", chip.snp$`POS (V3)`)
```

chip SNP calls are in AGPv3 positions. The hapmap 3 data are ALSO in AGPv3 positions, so we do NOT need to update marker positions. Yay!


Note, this data set includes some NAM founders as well as some nNILs.  
Could compare both of these groups to hapmap data. NAM founders on whole genome, nNILs only inside introgressions.

######################################################################

First compare NAM founders between chip and hapmap3 data sets across whole genome

Make bed file in AGP_V3 coordinates to use to subset HapMap 3 founder SNPs. 3 column format needed because other formats are misread by bcftools. Drop any SNPs that are not on chromosome assemblies in V3. 3-column format start is SNP position, end is at SNP position + 1 (but note I am constantly confused by this, other formats seem to differ) 
```{r}
bedChip = chip.snp %>% select(`#CHROM`, `POS (V3)`)
names(bedChip) = c('chrV3', 'startV3')
bedChip = bedChip %>%
  mutate(
    chrV3 = as.numeric(chrV3),
    startV3 = as.numeric(startV3),
    endV3 = startV3 + 1,
    name = paste0("S", chrV3, "_", startV3),
    score = 1,
    strand = "+"
  )  %>% 
  filter(!is.na(chrV3)) %>%
  select(chrV3, startV3, endV3)
#write.table(bedChip, paste0(base.path,"Data/nNIL_chip_SNP_positions_v3_6col.bed"),
#            row.names = F, col.names = F, quote = F, sep = "\t")
```
See HPC script "Y:\nNILs\nNIL_introgression_comparisons_to_founder_sequences.txt" for bcftools commands used to subset the HapMap3 SNPs to the nNIL founders and only the sites overlapping the chip SNPs. There are 18k SNPs in common between them.

Here is the subsetted GBS HapMap vcf with only the nNIL parents and only the SNPs that overlap with the chip markers
```{r}
founderVCF = read.vcfR("Y:/nNILs/HapMap3/nNIL_founders_chipSNPs_Hmp321_AGPv3.vcf.gz", verbose = FALSE )
```

Get the info in v3 on each SNP, get the original ordering of SNPs in vcf
```{r}
founder_meta = founderVCF@fix
```

Get the founder calls
```{r}
founder_calls =extract.gt(founderVCF, return.alleles = T)
founder_calls[founder_calls == "."] = NA
```

Convert the biallelic calls in founder_calls to single character IUPAC values, this is format used by the gbs_snp_v4 data frame.
```{r}
biallelic_to_IUPAC = function(x){
  if (is.na(x)) {return(x)}
  if (x == "A/A") {return("A")}
  if (x == "C/C") {return("C")}
  if (x == "G/G") {return("G")}
  if (x == "T/T") {return("T")}
  if (x == "A/C" | x == "C/A") {return("M")}
  if (x == "A/G" | x == "G/A") {return("R")}
  if (x == "A/T" | x == "T/A") {return("W")}
  if (x == "C/G" | x == "G/C") {return("S")}
  if (x == "C/T" | x == "T/C") {return("Y")}
  if (x == "G/T" | x == "T/G") {return("K")}
  return(NA) #in case no match
}
```

```{r}
founder_calls_iupac = apply(founder_calls, 1:2, biallelic_to_IUPAC)
colnames(founder_calls_iupac) = sub("282set_", "", colnames(founder_calls_iupac))
```

Now subset the chip SNP calls on nNIL lines to the same set of SNPs. 
Here, we have a problem that the position numbering system is off by 1 between the two data sets.
Add 1 to position of chip markers to make them equal to hapmap markers

Make sure missing values of N are set to NA. Also a few rare values need to be set to missing
```{r}
chip.snp.markers = apply(chip.snp[,1:2], 1, function(x) paste(x[1], as.numeric(x[2]) + 1, sep = "_")) #add 1 to marker position
hapmap.markers = apply(founder_meta[,1:2], 1, function(x) paste(x[1], x[2], sep = "_"))
chip.snp.subset = chip.snp.calls[,chip.snp.markers %in% hapmap.markers]
row.names(chip.snp.subset) = colnames(chip.snp)[-c(1:9)]
colnames(chip.snp.subset) = chip.snp.markers[chip.snp.markers %in% hapmap.markers]
all(colnames(chip.snp.subset) == hapmap.markers)
```
Subset the chip.snp.subset to only founder lines
```{r}
chip.snp.founders = chip.snp.subset[!grepl("NIL|DRIL|B73_\\(3.04\\)_L2|B73_JL1|B73_LL1|H100|NC262|NC304|Oh7B", rownames(chip.snp.subset)),]
```

Match rate ignoring hets between NAM founders in chip data and hapmap data, across whole genome
```{r}
compare_SNPs_whole_genome_nohets = function(x){
  line = rownames(chip.snp.founders)[x]
  chips = unlist(chip.snp.founders[line,])
  #set hets to missing
  hom.calls = c("A", "C", "G", "T")
  chips[!chips %in% hom.calls] = NA
  founders2 = apply(founder_calls_iupac, 1:2, function(x) if(x %in% hom.calls) {x} else {NA})
  
    compare = apply(founders2, 2, FUN = function(y) {sum(y == chips, na.rm = T)/sum(!is.na(y) & !is.na(chips))})
  return(compare)
}

 whole.list = lapply(1:nrow(chip.snp.founders), compare_SNPs_whole_genome_nohets)
 names(whole.list) = rownames(chip.snp.founders)
 whole.df = do.call(rbind, whole.list)
 write.csv(whole.df, file = paste0(base.path, "Output/founders_chip_vs_HapMap_whole_genome.csv"), row.names = T, quote = F)
```


Now compare the chip SNP calls inside introgressions to the hapmap SNP calls.
The introgression calls are in V4 positions.
```{r}
intros = read.csv(paste0(base.path, "Output/nNIL_chipdata_HMM_introgressionCalls.csv"))
```

Convert introgression markers to V3 positions. Use the V4 bed file with names reflecting V3 positions to make the translation.
ALSO NOTE ADD 1 TO v3 POSITION TO MATCH CHIP MARKER POSITIONS!
```{r}
v4_to_v3 = read.table(paste0(base.path,"Data/nNIL_chip_SNP_positions_converted_to_V4.bed"), header = F)
colnames(v4_to_v3) = c("chrV4", "startV4", "endV4", "nameV3", "score", "strand")
v4_to_v3 = v4_to_v3 %>%
  separate(nameV3, into = c("chrV3", "posV3"), sep = "_", remove = T) %>%
  mutate(chrV3 = as.numeric(sub("S", "", chrV3)),
         posV3 = as.numeric(posV3) + 1, #ADD ONE TO v3 POSITION!
         nameV3 = paste0("S", chrV3, "_" , posV3), #UPDATE V3 MARKER NAME!!
         chrV4 = as.numeric(chrV4),
         startV4 = as.numeric(startV4)) %>%
  filter(!is.na(chrV4))
```

Parse intro marker names into v4 chr and pos, then merge with the v4 to v3 translation table. 
Also subset keep only markers in the chip.snp.subset data frame
```{r}
intro.markers = data.frame(nameV4 = colnames(intros)[-c(1)]) %>%
  separate(nameV4, into = c("chrV4", "endV4"), sep = "_", remove = F) %>%
  mutate(chrV4 = as.numeric(sub("S", "", chrV4)),
         endV4 = as.numeric(endV4)) %>%
  inner_join(v4_to_v3, by = c("chrV4", "endV4")) %>%
  filter(nameV3 %in% paste0("S", colnames(chip.snp.subset)))
```


Have to matchup exactly intro.markers, order of columns of intros, and order of cols of chip.snp.calls, and order of rows of founder_calls_iupac. Start by transposing introgression calls matrix, merge with SNP info, then sort by V3 position
```{r}
introsT = as.data.frame(t(intros[,-c(1)]))
colnames(introsT) = intros[,1]
introsT$nameV4 = row.names(introsT)
introsT = inner_join(intro.markers, introsT, by = 'nameV4') %>%
  arrange(chrV3, posV3)
```

Check that markers are ordered the same in intro.markers and introsT
```{r}
all(intro.markers$nameV4 == introsT$nameV4)
```
Rename the NIL lines to be consistent
```{r}
colnames(introsT) = sub("B73 NIL", "B73_NIL", colnames(introsT))
```
Keep only introgression calls on the nNILs
```{r}
introsT = introsT %>%
  select(contains("V3"), contains("V4"), contains("B73_NIL"))
```

Finally, get the order of SNPs identical among the three data frames: founder_calls_iupac, introsT, and chip.snp.calls. Founder gbs hapmap and chip SNPs are in V3, but introgressions are in V4. 
```{r}
chip.snp.subset = chip.snp.subset[, paste0("S", colnames(chip.snp.subset)) %in% introsT$nameV3]
all(colnames(chip.snp.subset) == introsT$nameV3)
```
```{r}
all(paste0("S", colnames(chip.snp.subset))  %in% introsT$nameV3)
```
```{r}
all(introsT$nameV3 %in% paste0("S", colnames(chip.snp.subset)) )
```
So, markers in introsT are a subset of chip.snp.calls. Let's reduce chip.snp.calls to the overlapping set of markers and keep only the nNIL samples
```{r}
chip.snp.nNILs = chip.snp.subset[grepl("B73_NIL",row.names(chip.snp.subset)),paste0("S", colnames(chip.snp.subset)) %in% introsT$nameV3] 
dim(chip.snp.nNILs)
```
Now, check ordering
```{r}
all(introsT$nameV3 == paste0("S",colnames(chip.snp.nNILs)) )
```
Finally, check the ordering of markers in the founders
```{r}
all(introsT$nameV3 == row.names(founder_calls_iupac) )
```
No; Subset the founder calls to the overlapping SNPs. Have to add or remove the "S" at beginning of marker name to match between sets
```{r}
nNIL_markers_overlap = intersect(paste0("S", row.names(founder_calls_iupac)), introsT$nameV3)
length(nNIL_markers_overlap)
```
```{r}
introsT = introsT %>%
  filter(nameV3 %in% nNIL_markers_overlap)
dim(introsT)
```

```{r}
founder_calls_iupac2 = founder_calls_iupac[paste0("S", row.names(founder_calls_iupac)) %in% nNIL_markers_overlap,]
dim(founder_calls_iupac2)
```
```{r}
chip.snp.nNILs = chip.snp.nNILs[,paste0("S",colnames(chip.snp.nNILs)) %in% nNIL_markers_overlap]
dim(chip.snp.nNILs)
```
Re-check ordering one last time
```{r}
all(introsT$nameV3 == paste0("S", row.names(founder_calls_iupac2)))
```
```{r}
all(introsT$nameV3 == paste0("S", colnames(chip.snp.nNILs)))
```
OK! All the data frames are sorted properly!!!

make a function to apply to each line in chip set.
This function ignores hets in either nNIL or founder within introgression blocks
```{r}
compare_SNPs_in_intros_nohets = function(x){
  line = colnames(introsT[,-c(1:7)])[x]
  intro = as.logical(unlist(introsT[, colnames(introsT) == line]))
  chip = chip.snp.nNILs[line, intro]
  founders = founder_calls_iupac2[intro,] 

  #set hets to missing
  hom.calls = c("A", "C", "G", "T")
  chip[!chip %in% hom.calls] = NA
  founders2 = apply(founders, 1:2, function(x) if(x %in% hom.calls) {x} else {NA})
  
  compare = apply(founders2, 2, FUN = function(y) {sum(y == chip, na.rm = T)/sum(!is.na(y) & !is.na(chip))})
  return(compare)
}

nNIL_compare_list = lapply(1:ncol(introsT[,-c(1:7)]), compare_SNPs_in_intros_nohets)
names(nNIL_compare_list) = colnames(introsT)[-c(1:7)]
nNIL_compare.df = do.call(rbind, nNIL_compare_list)
write.csv(nNIL_compare.df, file = paste0(base.path, "Output/nNIL_subset_CHIP_vs_founders_inside_introgressions.csv"), row.names = T, quote = F)

```
