## Supplementary material for "A maize near-isogenic line population designed for gene discovery and characterization of allelic effects": File S20

---
title: "SNP_info_nNILs"
author: "Jim Holland"
date: "2024-06-17"
output: html_document
---

```{r setup, include=FALSE}
knitr::opts_chunk$set(echo = TRUE)
library(data.table)
library(tidyverse)
library(vcfR)
#base.path = "Q:/.shortcut-targets-by-id/1FP9BlrAC2EltlqKM3ounRs2COTiTK12G/nNIL genotype data Jim and Tao/"
base.path = "C:/Users/jholland/Box/nNIL genotype data Jim and Tao/"
```

# Founder HapMap Data 
Read in the founder HapMap SNPs in v3
```{r}
founderVCF = read.vcfR(paste0(base.path, "Data/nNIL_founders_consistent_gbsSNPs_v3.vcf.gz"), verbose = FALSE )
```

Get the info in v3 on each SNP, get the original ordering of SNPs in vcf
```{r}
founder_meta = founderVCF@fix
founder_meta = cbind(founder_meta, ord = 1:nrow(founder_meta))
```

Get the set of SNPs overlapping between nNIL gbs markers and HapMap3 markers with consistent REF/ALT alleles (identified in Step 4)
```{r}
consistentSNPs = read.csv(paste0(base.path, "Output/nNIL_gbs_SNPs_congruent_w_HapMap_v3v4v5coords_info.csv"))
```

Match the v3 position info with the v4 position info
```{r}
founder_meta_v4 = merge(founder_meta[,c("CHROM", "POS", "ord")], consistentSNPs, by.x = c("CHROM", "POS"), by.y = c("chrV3", "endV3")) %>%
  mutate(ord = as.numeric(ord)) %>%
  arrange(ord)
```

Verify the sorted V3 order matches the order of SNPs in founder matrix
```{r}
all(founder_meta_v4$POS == founder_meta[,"POS"])
```
Get the founder calls
```{r}
founder_calls =extract.gt(founderVCF, return.alleles = T)
founder_calls[founder_calls == "."] = NA
```

We verified above that the sorting of SNP names in founder_meta_v4 matches the founder VCF, so we can use the V4 SNP names to replace the V3 based row names.
```{r}
row.names(founder_calls) = founder_meta_v4$nameV4
colnames(founder_calls) = sub("282set_", "", colnames(founder_calls))
```

Convert the biallelic calls in founder_calls to single character IUPAC values, this is format used by the gbs_snp_v4 data frame.
```{r}
biallelic_to_IUPAC = function(x){
  if (is.na(x)) {return(x)}
  if (x == "A/A") {return("A")}
  if (x == "C/C") {return("C")}
  if (x == "G/G") {return("G")}
  if (x == "T/T") {return("T")}
  if (x == "A/C" | x == "C/A") {return("M")}
  if (x == "A/G" | x == "G/A") {return("R")}
  if (x == "A/T" | x == "T/A") {return("W")}
  if (x == "C/G" | x == "G/C") {return("S")}
  if (x == "C/T" | x == "T/C") {return("Y")}
  if (x == "G/T" | x == "T/G") {return("K")}
  return(NA) #in case no match
}
```

```{r}
founder_calls_iupac = apply(founder_calls, 1:2, biallelic_to_IUPAC)
```

# nNIL data

Get nNIL GBS SNPs in original V4 positions uplifted to V3 This is a subset after filtering to < 20% missing data in the vcf.
```{r}
gbs_v3 = read.table(paste0(base.path, "Data/nNIL_gbs_snps_converted_V3.bed"))
names(gbs_v3) = c("chrV3", "startV3", "endV3", "nameV4", "score", "strand")
gbs_v3 = gbs_v3 %>% separate(nameV4, into = c("chrV4", "posV4"), sep = "_", remove = F) %>%
  mutate(chrV4 = as.integer(sub("S", "", chrV4)))%>%
  select(-score, -strand)

```

Check GBS nNIL SNP info directly from hapmap meta data column alleles. Assuming first allele is reference
```{r}
gbs_snp_v4 = 
read.table(paste0(base.path,"Data/raw_snps_bgi_id.hmp.txt"), comment.char = "", header = T)
colnames(gbs_snp_v4)[1:4] = c("nameV4", "alleles", "chrV4", "posV4")
```

Now subset the GBS SNP calls on nNIL lines to the same set of SNPs. Make sure missing values of N are set to NA. Also a few rare values need to be set to missing
```{r}
gbs_snp_v4_consistent = gbs_snp_v4[gbs_snp_v4$nameV4 %in% row.names(founder_calls),]
gbs_snp_v4_consistent[gbs_snp_v4_consistent == "-"] = NA
gbs_snp_v4_consistent[gbs_snp_v4_consistent == "N"] = NA
gbs_snp_v4_consistent[gbs_snp_v4_consistent == "O"] = NA
```

change NIL line names from BGI codes to standard NIL line names
```{r}
name_translator = read.table(paste0(base.path, "Data/bgi_nil_id.txt"), sep = "\t", header = T, fileEncoding = "latin1")
name_translator = name_translator %>% 
  mutate(bgi_id = gsub("-", ".", bgi_id),
    nil_id = sub("B73.NIL", "B73 NIL", nil_id))
name_translator_vec = name_translator$nil_id
names(name_translator_vec) = name_translator$bgi_id
colnames(gbs_snp_v4_consistent)[12:ncol(gbs_snp_v4_consistent)] = name_translator_vec[colnames(gbs_snp_v4_consistent)[12:ncol(gbs_snp_v4_consistent)]]
```


Get the introgression calls on the nNILs
```{r}
intros = read.csv(paste0(base.path, "Output/nNIL_gbs_HMM_introgressionCalls_full_set.csv"))
```

Subset to the consistent set of SNPs in V4 positions that match the founder hapmap
```{r}
intros_sub = intros[,gbs_snp_v4_consistent$nameV4]
row.names(intros_sub) = intros[,1]
```

Keep only NILs whose line names match the gbs line names
```{r}
intros_sub = intros_sub[row.names(intros_sub) %in% colnames(gbs_snp_v4_consistent),]
```


Finally, get the order of SNPs identical among the three data frames: founder_calls_iupac, gbs_snp_v4_consistent, and intros_sub. Since the founder gbs and the nNIL gbs were aligned to different reference genomes, their ordering may vary and we need to make them identical
```{r}
all(rownames(founder_calls_iupac) == gbs_snp_v4_consistent$nameV4)
```
Resort the founder call SNPs to match nNILs. Need to parse the names into chrom and position to do this, then sort by NUMERIC chromosome and position
```{r}
founder_call_snps = strsplit(rownames(founder_calls_iupac), "_")
founder_call_snps = as.data.frame(do.call(rbind, founder_call_snps))
colnames(founder_call_snps) = c("chrV4", "posV4")
founder_call_snps$chrV4 = sub("S", "", founder_call_snps$chrV4)
founder_call_snps$nameV4 = rownames(founder_calls_iupac)
founder_calls_iupac = cbind(founder_call_snps, founder_calls_iupac)
founder_calls_iupac = as.data.frame(founder_calls_iupac) %>%
  mutate(chrV4 = as.numeric(chrV4),
         posV4 = as.numeric(posV4)) %>%
  arrange(chrV4, posV4)
```

Check that the new sorting matches the nNIL gbs marker order exactly
```{r}
all(founder_calls_iupac$nameV4 == gbs_snp_v4_consistent$nameV4)
```
Yay!!

Now check that intros_sub data.frame also has matching SNP order
```{r}
all(colnames(intros_sub) == gbs_snp_v4_consistent$nameV4)
```

Match rate ignoring hets within introgression blocks
```{r}
compare_SNPs_in_intros_nohets = function(x){
  line = row.names(intros_sub)[x]
  intro = as.logical(unlist(intros_sub[line,]))
  gbs = gbs_snp_v4_consistent[intro,line]
  founders = founder_calls_iupac[intro,-c(1:3)] #drop first three columns, they are meta-data
  if (length(gbs) <= 20){return(rep(NA, length(founders)))} #if <= 20 markers introgressed, too small to make meaningful match comparisons, so return a vector of NAs same length as the founders for this line.

  #set hets to missing
  hom.calls = c("A", "C", "G", "T")
  gbs[!gbs %in% hom.calls] = NA
  founders2 = apply(founders, 1:2, function(x) if(x %in% hom.calls) {x} else {NA})
  
  compare = apply(founders2, 2, FUN = function(y) {sum(y == gbs, na.rm = T)/sum(!is.na(y) & !is.na(gbs))})
  return(compare)
}

compare.list = lapply(1:nrow(intros_sub), compare_SNPs_in_intros_nohets)
names(compare.list) = row.names(intros_sub)
compare.df = do.call(rbind, compare.list)
write.csv(compare.df, file = paste0(base.path, "Output/nNIL_gbs_vs_founders_inside_introgressions.csv"), row.names = T, quote = F)
```
