## Supplementary material for "A maize near-isogenic line population designed for gene discovery and characterization of allelic effects": File S22

---
title: "Untitled"
author: "Jim Holland"
date: "2024-07-22"
output: html_document
---
```{r}
library(tidyverse)
base.path = "Q:/.shortcut-targets-by-id/1FP9BlrAC2EltlqKM3ounRs2COTiTK12G/nNIL genotype data Jim and Tao/"
```

# Test set: NAM founder match proportions inside called introgressions from chip data
Get the founder match results for the 24 nNILs included in the testing set from chip and GBS data and compare results
```{r}
chip_founder_matches = read.csv(paste0(base.path, "Output/nNIL_subset_CHIP_vs_founders_inside_introgressions.csv"))
colnames(chip_founder_matches)[1] = "Line"
```

Remove the underscore in line names in chip set to match with GBS set
```{r}
chip_founder_matches = chip_founder_matches %>%
  mutate(Line = gsub("_", " ", Line))
```

# NAM founder match proportions inside called introgressions from GBS data
```{r}
gbs_founder_matches = read.csv(paste0(base.path, "Output/nNIL_gbs_vs_founders_inside_introgressions.csv"))
colnames(gbs_founder_matches)[1] = "Line"
```

subset the GBS matches to the common test lines
```{r}
gbs_founder_matches_subset = gbs_founder_matches %>% filter(Line %in% chip_founder_matches$Line)
```

subset the chip matches to the common test lines (some lines in chip set are NOT in gbs)
```{r}
chip_founder_matches_subset = chip_founder_matches %>% filter(Line %in% gbs_founder_matches$Line)
```

Turn chip and gbs founder match dfs into matrices, make sure rows and columns are sorted identically
```{r}
chip.mat = as.matrix(chip_founder_matches_subset[,-c(1)])
row.names(chip.mat) = chip_founder_matches_subset$Line
chip.mat = chip.mat[order(row.names(chip.mat)), order(colnames(chip.mat))]
```

```{r}
gbs.mat = as.matrix(gbs_founder_matches_subset[,-c(1)])
row.names(gbs.mat) = gbs_founder_matches_subset$Line
gbs.mat = gbs.mat[order(row.names(gbs.mat)), order(colnames(gbs.mat))]

```
Check that rows are sorted identically
```{r}
all(rownames(chip.mat) == rownames(gbs.mat))
```

Check that cols are sorted identically
```{r}
all(colnames(chip.mat) == colnames(gbs.mat))
```
Overall correlation between founder match scores within introgressions
```{r}
cor(as.vector(chip.mat), as.vector(gbs.mat))
```
Check the correlations by NIL
```{r}
compare.founder.matches = sapply(1:nrow(chip.mat), FUN = function(x) cor(chip.mat[x,], gbs.mat[x,]))
names(compare.founder.matches) = row.names(chip.mat)
compare.founder.matches
```
Find the max founder match for ALL nNILs from GBS data
```{r}
find_best_match_num = function(row){
  if (all(is.na(row))) {return(NA)}
  x = row[which.max(row)]
  return(x)
}

find_best_match_name = function(row){
  nms = names(row)
  if (all(is.na(row))) {return(NA)}
  x = nms[which.max(row)]
  return(x)
}
besties = apply(gbs_founder_matches[,-c(1)], 1, find_best_match_num)
besties.names = unlist(apply(gbs_founder_matches[,-c(1)], 1, find_best_match_name))
best.df = data.frame(Line = gbs_founder_matches$Line, max.match = besties, founder.match = besties.names)
```

Also find max match from chip data
```{r}
besties.chip = apply(chip_founder_matches_subset[,-c(1)], 1, find_best_match_num)
besties.chip.names = unlist(apply(chip_founder_matches_subset[,-c(1)], 1, find_best_match_name))
best.chip.df = data.frame(Line = chip_founder_matches_subset$Line, max.match.chip = besties.chip, founder.match.chip = besties.chip.names)
```

Merge best matches from GBS and chip data, find the cases where best founder match disagrees
```{r}
best.both = merge(best.df, best.chip.df, by = "Line")
best.both %>% filter(founder.match != founder.match.chip)
```
Also find cases, where founder.match from gbs and chip data agree, but they disagree with pedigree
```{r}
best.both %>% mutate(pedigree.founder = sub("\\/.+", "", Line)) %>% filter(founder.match != pedigree.founder | founder.match.chip != pedigree.founder)
```
Now for each row, get the summary stats for the OTHER match proportions from GBS
```{r}
summary.non.match = function(row){
  if(all(is.na(row))) {return(c(min = NA, mean = NA, max = NA, sd = NA))}
  row[which.max(row)] = NA #set max to NA
  return(c(min = min(row,na.rm = T), mean = mean(row, na.rm = T),
         max = max(row, na.rm = T), sd = sd(row, na.rm = T)))
}

non.matches = apply(gbs_founder_matches[,-c(1)], 1, summary.non.match)
non.matches.df = as.data.frame(t(non.matches))
non.matches.df$Line = gbs_founder_matches$Line
```

Check the distribution of founder match probabilities for all except best match for each line
```{r}
non.match.mat = t(apply(gbs_founder_matches[,-c(1)], 1, function(x) {x[which.max(x)] = NA; return(x)}))
hist(as.vector(non.match.mat))
```
```{r}
summary(as.vector(non.match.mat))
```

Compare to distribution of best match
```{r}
hist(best.df$max.match)
```
```{r}
summary(best.df$max.match)
```

merge best and summary of non-matches
```{r}
best.and.non.matches = merge(best.df, non.matches.df, by = "Line")
```

compute the difference between best match and max match of the non-matches, also scale it to the sd, and look at the distribution of these stats.  
best.vs.2 = difference between best match and 2nd best match  
best.vs.2.std = best.vs.2 in terms of standard deviations of all but the best match
```{r}
best.and.non.matches = best.and.non.matches %>%
  mutate(best.vs.2 = max.match - max,
         best.vs.2.std = best.vs.2/sd)
```

check the distributions
```{r}
summary(best.and.non.matches[,c("best.vs.2", "best.vs.2.std")])
```

```{r}
hist(best.and.non.matches$best.vs.2)
```


```{r}
hist(best.and.non.matches$best.vs.2.std)
```


Plot the relationship between max.match and the difference between max and 2nd best match
```{r}
plot(best.and.non.matches$max.match, best.and.non.matches$best.vs.2,
     xlab = "Highest founder match proportion for each nNIL",
     ylab = "Highest - 2nd highest founder match")
```

Plot the relationship between max.match and the standardized difference between max and 2nd best match
```{r}
plot(best.and.non.matches$max.match, best.and.non.matches$best.vs.2.std,
     xlab = "Highest founder match proportion for each nNIL",
     ylab = "Highest - 2nd highest founder match standardized")
```
This is not very clear.

Declare a match when best match is > 0.8 and more than 1.5 SD greater than the 2nd best match proportion.

# Full set of nNILs

Using the rule set above, we can check the matches for the complete set of nNILs:
```{r}
all.matches = gbs_founder_matches %>%
  mutate(putative.donor = sub("\\/.*", "", Line)) %>%
  select(Line, putative.donor, everything())
```

Find the best match for each NIL.
Note that we have some NULL values in the resulting lists and unlist drops them. Convert NULLs to NAs before unlisting
```{r}
best.matches = apply(all.matches[,-c(1:2)], 1, find_best_match_num)
best.matches = lapply(best.matches, function(x) if (length(x) == 0){NA} else {x})
best.matches = unlist(best.matches)
best.matches.names = apply(all.matches[,-c(1:2)], 1, find_best_match_name)
best.matches.names = lapply(best.matches.names, function(x) if (length(x) == 0){NA} else {x})
best.matches.names = unlist(best.matches.names)
best.matches.df = cbind(all.matches[,c("Line", "putative.donor")], max.match = best.matches, founder.match = best.matches.names)
```

Summarize the non-best matches for each NIL
```{r}
non.matches.all = apply(all.matches[,-c(1:2)], 1, summary.non.match)
non.matches.all.df = as.data.frame(t(non.matches.all))
```
Merge info on best matches and other matches
```{r}
best.matches.all.df = cbind(best.matches.df, non.matches.all.df) %>%
  mutate(founder.match = case_when(
    max.match <= 0.8 ~ "Uncertain",
    (max.match - max)/sd < 1.5 ~ "Uncertain",
    T ~ founder.match
  ))
```


There are 18 lines that are NA for max.match. I checked them and their data look fine, but they have no introgressions called (or only a few SNPs called as introgressions).  Add one more filter, which is that an nNIL needs at least 20 SNPs remaining in the SNP set matching HapMap for match to be declared. If less than 20 SNPs, set founder.match to "IntrosTooSmall"  
Get the introgression calls on the nNILs
```{r}
intros = read.csv(paste0(base.path, "Output/nNIL_gbs_HMM_introgressionCalls_full_set.csv"))
```

Subset to the consistent set of SNPs in V4 positions
```{r}
consistentSNPs = read.csv(paste0(base.path, "Output/nNIL_gbs_SNPs_congruent_w_HapMap_v3v4v5coords_info.csv"))
intros_sub = intros[,colnames(intros) %in% consistentSNPs$nameV4]
row.names(intros_sub) = intros[,1]
```

Keep only NILs in the matches df
```{r}
intros_sub = intros_sub[row.names(intros_sub) %in% best.matches.all.df$Line,]
```

Summarize the number of SNPs scored as homozygous introgressions in the subset
```{r}
intro.num = apply(intros_sub, 1, function(x) sum(x ==2))
summary(intro.num)
```
Lines with <= 20 SNPs in introgression blocks should not be considered for matching to founders
```{r}
NILs.small.intros = names(intro.num)[intro.num <= 20]
length(NILs.small.intros)
```
Modify the 'founder.match' to "IntrosTooSmall" in the summary data frame
```{r}
best.matches.all.df = best.matches.all.df %>%
  mutate(founder.match = ifelse(Line %in% NILs.small.intros, "IntrosTooSmall", founder.match))
```


One or more founder names have a spelling difference between HapMap and NIL line names, have to correct that before comparing putative.donor to founder.match
```{r}
all(best.matches.all.df$putative.donor %in% best.matches.all.df$founder.match)
```
```{r}
unique(best.matches.all.df$putative.donor[!best.matches.all.df$putative.donor %in% best.matches.all.df$founder.match])
```
Just need to fix spelling of Ki3 in founder.match column
```{r}
best.matches.all.df = best.matches.all.df %>%
  mutate(founder.match = ifelse(founder.match == "KI3", "Ki3", founder.match))
```


How many matches are uncertain, and how many disagree with pedigree?
```{r}
missing.matches = best.matches.all.df %>% filter(is.na(putative.donor)) #0 NAs in putative.donor column
uncertains = best.matches.all.df %>% filter(founder.match == "Uncertain")
wrong = best.matches.all.df %>%
  filter((founder.match != putative.donor) & (! founder.match %in% c("Uncertain", "IntrosTooSmall")))
too.small = best.matches.all.df %>% filter(founder.match == "IntrosTooSmall")
```

Proportion of uncertain matches
```{r}
nrow(uncertains)/(nrow(best.matches.all.df))
```
Proportion of wrong matches
```{r}
nrow(wrong)/(nrow(best.matches.all.df))
```
Proportion of NILs with introgressions too small
```{r}
nrow(too.small)/(nrow(best.matches.all.df))
```

Combine all the info and write to a file
```{r}
final.df = merge(best.matches.all.df, all.matches, by = c("Line", "putative.donor")) %>%
  rename(min.others = min,
         mean.others = mean,
         max.others = max,
         sd.others = sd)
write.csv(final.df, paste0(base.path, "Output/Founder_matches_all_nNILs.csv"), row.names = F, quote = F)
```

Summarize the mismatches by pedigree:
```{r}
mismatch.by.pedigree = final.df %>% 
  mutate(mismatch = (putative.donor != founder.match) & ! founder.match %in% c("Uncertain", "IntrosTooSmall"))  %>%
  group_by(putative.donor) %>%
  summarize(N = n(), proportion.mismatch = sum(mismatch)/n())
mismatch.by.pedigree
```
```{r}
write.csv(mismatch.by.pedigree, paste0(base.path, "Output/Founder_mismatch_proportion_by_pedigree.csv"),row.names = F, quote = F)
```
