## Supplementary material for "A maize near-isogenic line population designed for gene discovery and characterization of allelic effects": File S23

---
title: "nNIL introgression qtl tests"
author: "Jim Holland"
date: "2024-11-21"
output: html_document
---

```{r setup}
knitr::opts_chunk$set(echo = TRUE)
library(tidyverse)
```

Get the data with pruned introgression markers
```{r}
blues_intros = read.csv("C:/Users/jholland/Box/nNIL genotype data Jim and Tao/GWAS/nNIL_BLUEs_pruned_introgression_markers.csv") |>
  select(-ends_with("B73"))
```

Get list of marker names
```{r}
markers = colnames(blues_intros)[-c(1:7)]
```

Get the corrected donor information for each nNIL, drop nNILs with uncertain donors
```{r}
nNIL.donors = read.csv("C:/Users/jholland/Box/nNIL genotype data Jim and Tao/Output/Founder_matches_all_nNILs.csv") |>
  select(Line, donor = founder.match) |>
  filter(!donor  %in% c("IntrosTooSmall", "Uncertain", "B73") ) |>
  mutate(Geno = sub(" ", "_", Line))
```

Merge blues with corrected donor info
```{r}
blues_intros2 = merge(blues_intros, nNIL.donors, by = 'Geno')
```


Make a function to fit QTL tests. 
Model 1 fits common effect for all donor alleles at each introgression
```{r}
QTL_test_common_effect = function(df, trait, marker){
  y = df[[trait]]
  x = as.factor(df[[marker]])
  mod = lm(y ~ x)
  mod.sum = summary(mod)$coefficients
  mod.sum = as.data.frame(mod.sum[-c(1),c("Estimate", "Pr(>|t|)"), drop = F])
  colnames(mod.sum)[2] = "p.value"
  mod.sum[["Marker"]] = marker
  mod.sum[["Intro_Dosage"]] = row.names(mod.sum)
  mod.sum[["Trait"]] = trait
  return(mod.sum)
}
```

Run the tests
```{r}
results.common.effect.list = list()
for (t in c("GLS", "NLB", "SLB")){
  for (m in markers){
    results.common.effect.list[[paste0(t, "_", m)]] = QTL_test_common_effect(blues_intros2, t, m)
  }
}
```

Combine results
```{r}
qtl.results.common.effect.df = do.call(bind_rows, results.common.effect.list)
```

Check p-value distribution
```{r}
ggplot(qtl.results.common.effect.df, aes(x = p.value)) + geom_histogram(aes(colour = Trait)) +
  facet_wrap(facets = vars(Trait)) +
  theme_minimal()
```
Count significant tests
```{r}
bonf = 0.05/297 #this is computed in xQTL script
qtl.results.common.effect.df |> group_by(Trait) |>
  summarise(N.sign = sum(p.value < bonf))
```

Plot QTL results from model with common donor effects
```{r}
qtl.common.effect.results.sign.df = qtl.results.common.effect.df |> filter(p.value < bonf) |>
  separate(Marker, into = c("chr", "pos"), sep = "_", remove = F) |>
  mutate(chr = as.numeric(sub("S", "", chr)),
         pos = as.numeric(pos),
         Mbp = pos/1000000,
         log10p = -log10(p.value))
```

```{r}
qtl.common.effect.hits.plot = ggplot(qtl.common.effect.results.sign.df, aes(x = Mbp, y = log10p)) +
  geom_point(aes(colour = Trait)) +
  facet_wrap(facets = vars(chr), scales = "free_y") +
  ylab("-log10(p-value)") +
  ggtitle("nNIL introgressions associated with disease traits\ncommon donor effect model") +
  theme_minimal()
qtl.common.effect.hits.plot 
```


Write out results of common QTL effect tests
```{r}
write.csv(qtl.common.effect.results.sign.df , file = "C:/Users/jholland/Box/nNIL genotype data Jim and Tao/GWAS/nNIL_common_introgression_effect_significant_results.csv", row.names = F, quote = F)
```


# Function for forward regression common effect qtl tests
```{r}
forwardQTL = function(df, trait, marker.df, p.thresh = 0.01){
  #df is data frame of trait data and marker scores
  #trait is character string of trait name
  #marker.df is a data.frame with two columns: Marker with character marker names and p.value which are p.values from initial QTL screen
  minp.marker = marker.df |> filter(p.value < p.thresh) 
  if (nrow(minp.marker) == 0) {return(NULL)}
  minp.marker = minp.marker |> slice_min(order_by = p.value, n = 1, with_ties = F) |> pull(Marker)
  marker.df = marker.df |> filter(Marker != minp.marker)

  mod = lm(as.formula(paste(trait, " ~ 1 + ", minp.marker)), data = df) #base model
  best.mod = mod
  while (min(marker.df$p.value < 0.01)){
    results.list = list()
    for (m in marker.df$Marker){
      new.mod = update(best.mod, as.formula(paste("~ . +", m)))
      if (m %in% row.names(summary(new.mod)$coefficients)){ #if new marker causes singularity it will not be in rownames of summary, so have to check for that
      results.list[[m]] = data.frame(Marker = m, p.value = summary(new.mod)$coefficients[m,4])} 
    }
    marker.df = do.call(rbind, results.list) |> filter(p.value < p.thresh) 
    minp.marker = slice_min(marker.df, order_by = p.value, n = 1, with_ties = F) 
    if (nrow(minp.marker) == 0) {return(best.mod)}
    minp.marker = minp.marker |> pull(Marker)
    marker.df = marker.df |> filter(Marker != minp.marker)
    best.mod = update(best.mod, as.formula(paste("~ . +", minp.marker)))
    }
return(best.mod)
}
```


Forward regression to select subset of markers associated with each trait.  
Filter markers to include only those with p < 0.01 as potential markers to include in model
```{r}
gls.multimod = forwardQTL(blues_intros2, "GLS", qtl.common.effect.results.sign.df[qtl.common.effect.results.sign.df$Trait == "GLS",c("Marker", "p.value")])
nlb.multimod = forwardQTL(blues_intros2, "NLB", qtl.common.effect.results.sign.df[qtl.common.effect.results.sign.df$Trait == "NLB",c("Marker", "p.value")])
slb.multimod = forwardQTL(blues_intros2, "SLB", qtl.common.effect.results.sign.df[qtl.common.effect.results.sign.df$Trait == "SLB",c("Marker", "p.value")])
```

Summarize final multiple QTL model fits
```{r}
summary(gls.multimod)
```
```{r}
summary(nlb.multimod)
```
```{r}
summary(slb.multimod)
```

Make a 2nd function to test introgression*donor interactions. 
Now we have to treat introgression dosage as a numeric value (regressor) with common intercept at 0.  
Otherwise, if marker value is a factor, this model will fit a separate level for the non-introgressed lines for each donor, which is not what we want.
Once we have a list of markers-traits of interest, we can get their effect estimates later.
```{r}
Intro_Donor_test = function(df, trait, marker){
  y = df[[trait]]
  a = as.numeric(df[[marker]])
  d = 1 - abs(a - 1)# d is 1 for het and 0 for others
  donor = as.factor(df[['donor']])
  mod = aov(y ~ a:donor + d:donor)
  mod.df = summary(mod)[[1]]
  mod.a = mod.df[grepl("^a:donor|donor:a$", trimws(row.names(mod.df))),]
  
  #check if any d effects in model result
  if (any(grepl("^d:donor|donor:d$", trimws(row.names(mod.df))))){
  mod.d = mod.df[grepl("^d:donor|donor:d$", trimws(row.names(mod.df))),]
  DF_dom = mod.d[1,"Df"]
  pval.dom = mod.d[1,"Pr(>F)"]} else { 
    #handle case when no dominance effects in results
    DF_dom = 0
    pval.dom = NA}
  
  coefs = mod$coefficients
  results.df = cbind(data.frame('Marker' = marker, 'Trait' = trait, 'DF_add' = mod.a[1,"Df"], 'DF_dom' = DF_dom, pval.add = mod.a[1,"Pr(>F)"], pval.dom = pval.dom), t(coefs))
  return(results.df)
}
```
Run it
```{r}
donor.intro.results.list = list()
for (t in c("GLS", "NLB", "SLB")){
  for (m in markers){
    donor.intro.results.list [[paste0(t, "_", m)]] = Intro_Donor_test(blues_intros2, t, m)
  }
}
```

```{r}
qtl.donor.intro.results.df = do.call(bind_rows, donor.intro.results.list)
```

Check p-value distribution
```{r}
ggplot(qtl.donor.intro.results.df, aes(x = pval.add)) + geom_histogram(aes(colour = Trait)) +
  facet_wrap(facets = vars(Trait)) +
  theme_minimal()
```
Display significant QTL hits across the genome
```{r}
qtl.donor.intro.results.sign.df = qtl.donor.intro.results.df |> filter(pval.add < bonf) |>
  separate(Marker, into = c("chr", "pos"), sep = "_", remove = F) |>
  mutate(chr = as.numeric(sub("S", "", chr)),
         pos = as.numeric(pos),
         Mbp = pos/1000000,
         log10p = -log10(pval.add))
```


```{r}
qtl.donor.hits.plot = ggplot(qtl.donor.intro.results.sign.df, aes(x = Mbp, y = log10p)) +
  geom_point(aes(colour = Trait)) +
  facet_wrap(facets = vars(chr), scales = "free_y") +
  ylab("-log10(p-value)") +
  ggtitle("nNIL introgressions associated with disease traits\nmarker-by-donor interaction model") +
  theme_minimal()
qtl.donor.hits.plot  
```
Write out results of donor-by-introgression QTL effect tests
```{r}
write.csv(qtl.donor.intro.results.sign.df , file = "C:/Users/jholland/Box/nNIL genotype data Jim and Tao/GWAS/nNIL_donor_by_introgression_interaction_significant_results.csv", row.names = F, quote = F)
```

# Function for forward regression donor*QTL interaction tests
```{r}
forwardQTL.donor = function(df, trait, marker.df){
  #df is data frame of trait data and marker scores
  #trait is character string of trait name
  #marker.df is a data.frame with two columns: Marker with character marker names and p.value which are p.values from initial QTL screen
  minp.marker = marker.df |> slice_min(order_by = p.value, n = 1, with_ties = F) |> pull(Marker)
  marker.df = marker.df |> filter(Marker != minp.marker)
  donor = as.factor(df[['donor']])


  mod = lm(as.formula(paste0(trait, " ~ 1 + ", minp.marker, ":donor")), data = df) #base model
  best.mod = mod
  best.aic = AIC(mod)
  keep.on = T
  while (keep.on){
    results.list = list()
    for (m in marker.df$Marker){
      new.mod = update(best.mod, as.formula(paste0("~ . +", m, ":donor")))
      results.list[[m]] = data.frame(Marker = m, aic = AIC(new.mod))} 
    marker.df = do.call(rbind, results.list) 
    minAIC.marker.df = slice_min(marker.df, order_by = aic, n = 1, with_ties = F) 
    if (nrow(minAIC.marker.df) == 0) {return(best.mod)}
    if (minAIC.marker.df$aic >= best.aic) {return(best.mod)}
    minAIC.marker = minAIC.marker.df |> pull(Marker)
    marker.df = marker.df |> filter(Marker != minAIC.marker)
    best.mod = update(best.mod, as.formula(paste0("~ . + ", minAIC.marker, ":donor")))
    best.aic = minAIC.marker.df |> pull(aic)
    }
return(best.mod)
}
```

Forward regression to select subset of markers associated with each trait.  
Filter markers to include only those with pval.add < 0.01 as potential markers to include in model
```{r}
qtl.donor.intro.results.sign.gls = qtl.donor.intro.results.sign.df |>
  filter(Trait == "GLS") |>
  rename(p.value = pval.add)
gls.donor.multimod = forwardQTL.donor(blues_intros2, "GLS", qtl.donor.intro.results.sign.gls)


qtl.donor.intro.results.sign.nlb = qtl.donor.intro.results.sign.df |>
  filter(Trait == "NLB") |>
  rename(p.value = pval.add)
nlb.donor.multimod = forwardQTL.donor(blues_intros2, "NLB", qtl.donor.intro.results.sign.nlb)


qtl.donor.intro.results.sign.slb = qtl.donor.intro.results.sign.df |>
  filter(Trait == "SLB") |>
  rename(p.value = pval.add)

slb.donor.multimod = forwardQTL.donor(blues_intros2, "SLB", qtl.donor.intro.results.sign.slb )
```

Summarize final multiple QTL model fits
```{r}
summary(gls.donor.multimod)
```
```{r}
summary(nlb.donor.multimod)
```

```{r}
summary(slb.donor.multimod)
```

Combine final multi-QTL models and write to hard drive
```{r}
gls.common = as.data.frame(summary(gls.multimod)$coefficients)
gls.common$Marker = row.names(gls.common)
gls.common$Trait = "GLS"

nlb.common = as.data.frame(summary(nlb.multimod)$coefficients)
nlb.common$Marker = row.names(nlb.common)
nlb.common$Trait = "NLB"

slb.common = as.data.frame(summary(slb.multimod)$coefficients)
slb.common$Marker = row.names(slb.common)
slb.common$Trait = "SLB"

all.multiQTL.common = rbind(gls.common, nlb.common, slb.common)
write.csv(all.multiQTL.common, file = "C:/Users/jholland/Box/nNIL genotype data Jim and Tao/GWAS/Common_Effect_MultiQTL_Results.csv", row.names = F, quote = F)
```

```{r}
gls.donor = as.data.frame(summary(gls.donor.multimod)$coefficients)
gls.donor$Marker = row.names(gls.donor)
gls.donor$Trait = "GLS"

nlb.donor = as.data.frame(summary(nlb.donor.multimod)$coefficients)
nlb.donor$Marker = row.names(nlb.donor)
nlb.donor$Trait = "NLB"

slb.donor = as.data.frame(summary(slb.donor.multimod)$coefficients)
slb.donor$Marker = row.names(slb.donor)
slb.donor$Trait = "SLB"

all.multiQTL.donor = rbind(gls.donor, nlb.donor, slb.donor) |>
  filter(!is.na(Estimate))
write.csv(all.multiQTL.donor, file = "C:/Users/jholland/Box/nNIL genotype data Jim and Tao/GWAS/Donor_Effect_MultiQTL_Results.csv", row.names = F, quote = F)
```
